# An acylsugar-deficient *Nicotiana benthamiana* strain for aphid and whitefly research

**DOI:** 10.1101/2020.08.04.237180

**Authors:** Honglin Feng, Lucia Acosta-Gamboa, Lars H. Kruse, Jake D. Tracy, Seung Ho Chung, Alba Ruth Nava Fereira, Sara Shakir, Hongxing Xu, Garry Sunter, Michael A. Gore, Clare L. Casteel, Gaurav D. Moghe, Georg Jander

**Affiliations:** Boyce Thompson Institute, Ithaca NY, USA; Plant Breeding and Genetics Section, School of Integrative Plant Science, Cornell University, Ithaca NY, 14853, USA; Plant Biology Section, School of Integrative Plant Science, Cornell University, Ithaca NY, 14853, USA; Plant-Microbe Biology and Plant Pathology Section, School of Integrative Plant Science, Cornell University, Ithaca NY, 14853, USA; Department of Biology, University of Texas San Antonio, San Antonio TX, 78249, USA; Michael Smith Laboratories, University of British Columbia, Vancouver, BC, V6T 1Z4, Canada; Gembloux Agro-Bio Tech Institute, the University of Liege, Gembloux, Belgium; College of Life Science, the Shaanxi Normal University, Xi’an, China

## Abstract

*Nicotiana benthamiana* is used extensively as a platform for transient gene expression and as a model system for studying plant-virus interactions. However, many tobacco-feeding generalist herbivores, including *Myzus persicae* (green peach aphid), *Bemisia tabaci* (whitefly), *Macrosiphum euphorbiae* (potato aphid), *Heliothis virescens* (tobacco budworm), *Trichoplusia ni* (cabbage looper), and *Helicoverpa zea* (corn earworm), grow poorly on *N. benthamiana*, limiting its utility for research on plant-insect interactions. Using CRISPR/Cas9, we generated knockout mutations in two *N. benthamiana* acylsugar acyltransferases, *ASAT1* and *ASAT2*, which contribute to the biosynthesis of insect-deterrent acylsucroses. Whereas *asat1* mutations reduced the abundance of two predominant acylsucroses, *asat2* mutations caused almost complete depletion of foliar acylsucroses. The tested hemipteran and lepidopteran species survived, gained weight, and/or reproduced significantly better on *asat2* mutant plants than on wildtype *N. benthamiana*. Furthermore, both *asat1* and *asat2* mutations reduced the water content and increased the temperature of leaves, indicating that foliar acylsucroses can protect against desiccation. Two experiments demonstrated the utility of the *N. benthamiana asat2* mutant line for insect bioassays. Transmission of turnip mosaic virus by *M. persicae* was significantly improved by an *asat2* mutation. Tobacco rattle virus constructs were used for virus-induced gene silencing of *acetylcholinesterase*, *squalene synthase*, *toll-like receptor 7*, and *tubulin-specific chaperon D* genes in *B. tabaci*, an experiment that would have been difficult with wild-type *N. benthamiana* due to high insect mortality. Additionally, the absence of acylsugars in *asat2* mutant lines will simplify transient expression assays for the functional analysis of acylsugar biosynthesis genes from other Solanaceae.

## Introduction

*Nicotiana benthamiana*, a wild tobacco species that is native to Australia, is commonly used by plant molecular biologists as model system for laboratory research. Susceptibility to a wide variety of plant viruses has made *N. benthamiana* a popular model for fundamental studies of plant-virus interactions (Goodin et al., 2008). Scientists have developed *N. benthamiana* as a transgene expression powerhouse by engineering viral vectors to express heterologous genes, including fluorescent reporter genes to visualize cell structures (Bally et al., 2018). Antibodies, biofuel compounds, and other protein and metabolite products have been produced in *N. benthamiana* (Arntzen, 2015; Powell, 2015). Virus-induced gene silencing (VIGS), which is employed to study gene function in a variety of plant species, was originally developed in *N. benthamiana* (Velásquez et al., 2009; Hayward et al., 2011). Recently, high-efficiency CRISPR/Cas9 gene editing of germline cells using virus-encoded guide RNA (gRNA) was demonstrated for the first time in *N. benthamiana* (Ellison et al., 2020). Although *N. benthamiana* is hyper-susceptible to many plant viruses, it is not a good host for three virus-transmitting Hemiptera, *Myzus persicae* (green peach aphid; Thurston, 1961; Hagimori et al., 1993), *Bemisia tabaci* (whitefly; Simon et al., 2003), and *Macrosiphum euphorbiae* (potato aphid) that otherwise grow well on cultivated tobacco (*Nicotiana tabacum*).

The poor growth of generalist insect herbivores on *N. benthamiana* may be attributed in part to glandular trichomes. These epidermal secretory structures on the leaf surface of ∼30% of vascular plants (Weinhold and Baldwin, 2011; Glas et al., 2012) have been found to play a crucial defensive role in several ways: as a physical obstacle for insect movement on the plant surface (Cardoso, 2008), entrapment (Simmons et al., 2004), synthesis of volatiles and other defensive metabolites (Laue et al., 2000; Schilmiller et al., 2010; Glas et al., 2012), and production of proteins that repel herbivores (phylloplane proteins, e.g. T-phylloplanin; Shepherd and Wagner, 2007). In addition to their defensive functions, glandular trichomes also protect plants from abiotic stresses such as transpiration water loss and UV irradiation (Karabourniotis et al., 1995).

There are two main types of glandular trichomes on *N. benthamiana* leaves, large swollen-stalk trichomes and small trichomes that are capped by a secretory head with one, two, or four cells (Slocombe et al., 2008). The large trichomes have been shown to secrete phylloplane proteins in *N. tabacum*. The small trichomes, which are the most abundant on tobacco leaf surfaces, secrete exudates, including acylsugars (Wagner et al., 2004; Slocombe et al., 2008). Detached trichomes, a mixture of the large and small trichomes, from *N. benthamiana* are able to synthesize acylsugars (Kroumova and Wagner, 2003), and the secretory head cells alone are able to synthesize acylsugars in *N. tabacum* (Kandra and Wagner, 1988).

Acylsugars, generally sucrose or glucose esterified with aliphatic acids of different chain lengths (Figure 1), are abundant insect-deterrent metabolites produced by Solanaceae glandular trichomes (Arrendale et al., 1990; Slocombe et al., 2008; Moghe et al., 2017). Specific acylsugars are associated with aphid-resistant *Nicotiana* species, while not being detected in more susceptible species in this genus (Hagimori et al., 1993). Furthermore, relative to cultivated tomatoes (*Solanum lycopersicum*), acylsugars provide wild tomatoes (*Solanum pennellii*) greater resistance against *M. persicae* (Rodriguez et al., 1993). Diacylsucrose protects crops against tobacco aphids (*Myzus persicae nicotianae*), *B. tabaci*, and two-spotted spider mites (*Tetranychus urticae*) (Chortyk et al., 1996; Alba et al., 2009). The synthetic sucrose octanoate (an analog of *Nicotiana gossei* sugar esters) is effective in the field against Asian citrus psyllids (*Diaphorina citri*), citrus leafminer (*Phyllocnistis citrella*), and a mite complex (including Texas citrus mite, red spider mite, and rust mite) (McKenzie and Puterka, 2004; McKenzie et al., 2005). Interestingly, acylsucroses in *Nicotiana attenuata* are metabolized to volatile fatty acids by neonate *Manduca sexta* (tobacco hornworm) larvae, thereby tagging these larvae and attracting predatory ants, *Pogonomyrmex rugosus* (Weinhold and Baldwin, 2011).

**Figure 1.**
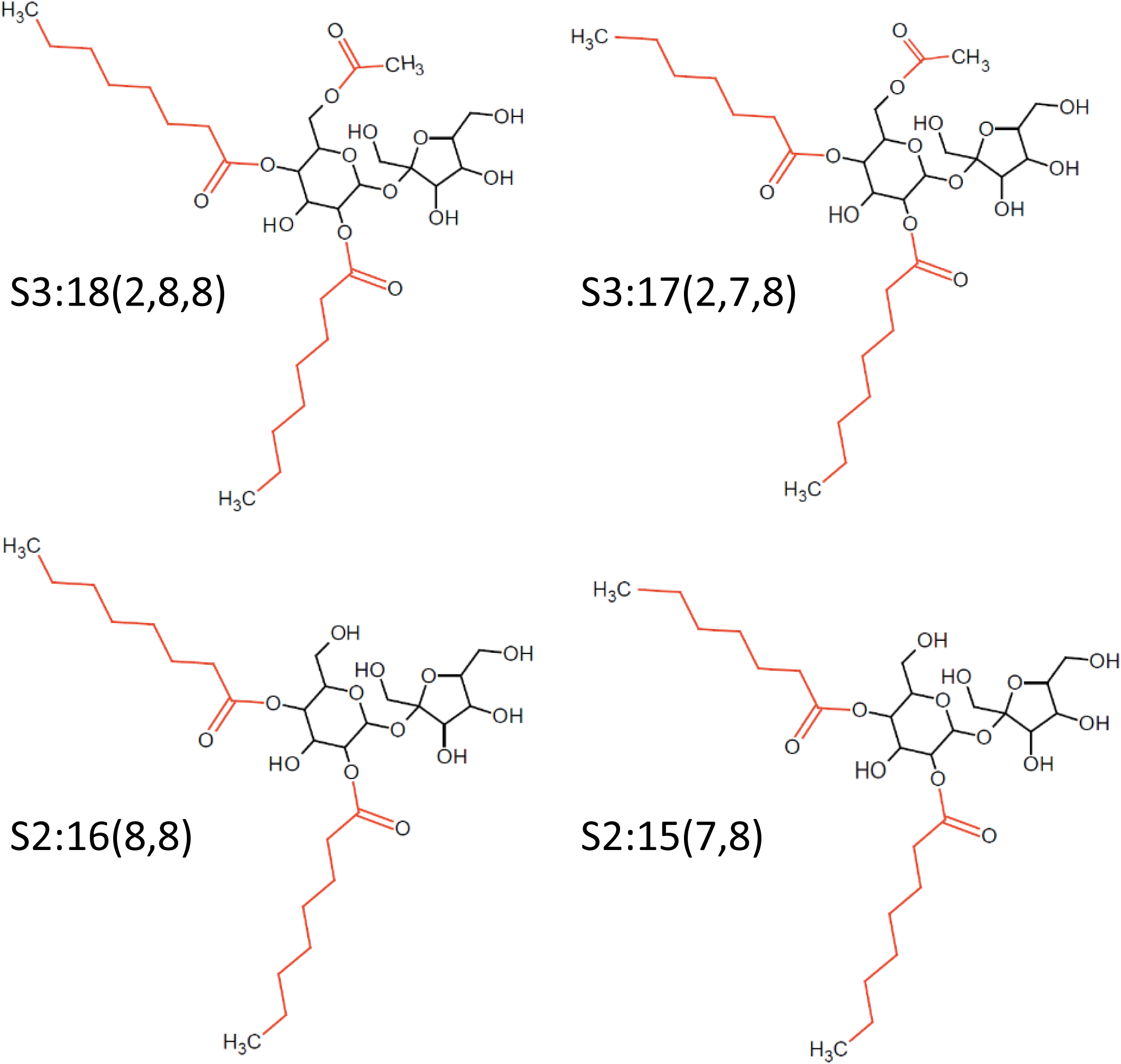
**Predominant acylsugars in *Nicotiana benthamiana*** Acylsucroses S3:17(2,7,8), S3:18(2,8,8), S2:16(8,8) and S2:15(7,8) are present in *N. benthamiana*. In the acylsugar structure names, S refers to sucrose, followed by the number of acyl chains, the total length of acyl chains, and the length of each individual chain in parentheses. Although the presence of C2, C7, and C8 chain lengths is confirmed, the specific positions of the acyl chains on the sucrose molecule are hypothesized based on previous observations of acylsugars in *Nicotiana alata* (Moghe et al, 2017), the predicted evolution of the acylsugar biosynthetic pathway, and enzyme promiscuities in the Solanaceae family.

Acylsugars and leaf surface lipids more generally may contribute to plant drought tolerance. Transcriptomic studies of drought-tolerant *S. pennellii* populations showed that lipid metabolism genes are among those that are most responsive to drought stress (Gong et al., 2010; Egea et al., 2018). Additionally, the abundance of acylsugars on the leaf surface in a native *S. pennellii* population was associated with drought tolerance (Fobes et al., 1985). Acylsugars also are reported to provide protection against drought stress conditions in *Solanum chilense* (O’ Connell et al., 2007). Similarly, abundant accumulation of acylsugars with C7-C8 acyl groups in the desert tobacco (*Nicotiana obtusifolia*) was suggested to provide this species with high drought tolerance for its desert environment (Kroumova et al., 2016). Although the mechanism is not completely understood, it has been proposed that the polar lipids reduce the surface tension of adsorbed dew water, thereby allowing the leaves absorb more condensed water on the surface (Fobes et al., 1985).

More recently, enzymes involved in the biosynthesis of acylsugars have been identified. Four acylsugar acyltransferases (ASATs), *Sl*ASAT1, *Sl*ASAT2, *Sl*ASAT3, and *Sl*ASAT4, have been biochemically characterized in cultivated tomato (Fan et al., 2016). *Sl*ASAT1 catalyzes the first step of sucrose acylation, using sucrose and acyl-CoA to generate monoacylsucroses via pyranose R_4_ acylation (Fan et al., 2016). *Sl*ASAT2 uses the product of *Sl*ASAT1 (*e.g.* S1:5(iC5^R4^)) and acyl-CoA donor substrates (*e.g.* iC4-CoA, aiC5-CoA, nC10-CoA, and nC12-CoA) to generate diacylsucroses (Fan et al., 2016). Further, *Sl*ASAT3 uses the diacylsucroses generated by *Sl*ASAT2 to make triacylsucroses by acylating the diacylsucrose five-membered (furanose) ring (Fan et al., 2016). Then, *Sl*ASAT4 makes tetraacylsucroses by acetylating triacylsucroes using C2-CoA (Schilmiller et al., 2012). Notably, *Sl*ASAT4 (formerly *Sl*ASAT2) is specifically expressed in the trichomes, where acylsugar acetylation occurs (Schilmiller et al., 2012). The expression and activity of ASATs varies among different plant species (including the order of the ASAT reactions in the pathway), which likely contributes to the observed trichome chemical diversity (Kim et al., 2012). Although ASATs have been most intensively studied in tomato, *ASAT* genes also have been annotated in the available *Nicotiana* genomes (Gaquerel et al., 2013; Van et al., 2017; Egan et al., 2019). However, *ASAT* genes in *N. benthamiana* have not been annotated and characterized previously, and their functions in protection against insect pests and desiccation remained unknown.

The goal of this study was to confirm the role of acylsugars in *N. benthamiana* resistance to insect feeding, as well as to create an insect-susceptible *ASAT* mutant line to facilitate use of *N. benthamiana* for laboratory research on plant-insect interactions. We identified two *ASAT* genes in *N. benthamiana*, *NbASAT1 and NbASAT2*. Using CRISPR/Cas9 to create mutant lines, we showed that knockout of both *NbASAT1* and *NbASAT2* reduced acylsugar content. *NbASAT2* mutations allowed increased survival, growth, and/or reproduction of six tested insect species. Additionally, decreased water content and elevated leaf temperature in *NbASAT1* and *NbASAT2* mutants indicated that *N. benthamiana* acylsugars contribute to protection against desiccation.

## Results

### Identification of *ASAT1* and *ASAT2* in N. benthamiana

Using reciprocal comparisons to confirmed Solanaceae *ASAT* genes (Moghe et al., 2017), we identified three highly homologous sequences in the *N. benthamiana* genome: Niben101Scf02239Ctg025, Niben101Scf22800Ctg001, and Niben101Scf14179Ctg028 (Bombarely et al., 2012) (gene identifiers are from annotations at solgenomics.net). Whereas Niben101Scf02239Ctg025 and Niben101Scf22800Ctg001 were annotated as a full-length coding sequences with strong coverage in available RNAseq datasets, Niben101Scf14179Ctg028 was annotated as a pseudogene because it appears to be a fragment of the predicted cDNA Niben101Scf141790g02010.1, but with no coverage in available RNAseq datasets. The Niben101Scf02239Ctg025 and Niben101Scf22800Ctg001 sequences were confirmed in a more recently assembled *N. benthamiana* genome (Schiavinato et al., 2019). In this assembly, the pseudogene Niben101Scf14179Ctg028 was annotated as part of Niben101Scf02239Ctg025, and there were no additional annotated ASAT candidates.

To infer ASAT evolution and function, we constructed a protein phylogenetic tree of previously annotated Solanaceae ASATs (Figures 2, S1 and S2; Tables S1 and S2). In this tree, Niben101Scf02239Ctg025 formed a monophyletic group with other ASATs, including the biochemically characterized *Ss*ASAT1, *Pa*ASAT1 and *Hn*ASAT1. Therefore, we named Niben101Scf02239Ctg025 as *N. benthamiana* ASAT1 (*Nb*ASAT1). Niben101Scf22800Ctg001 formed a monophyletic group with other ASATs including the biochemically characterized *Na*ASAT2, *Hn*ASAT2, and *Pa*ASAT2. Therefore, we named Niben101Scf22800Ctg001 as *N. benthamiana* ASAT2 (*Nb*ASAT2). Notably, the ASAT2 monophyletic group also included the biochemically characterized *Sp*ASAT1, *Sl*ASAT1, and *Sn*ASAT1 (Figure 2).

**Figure 2.**
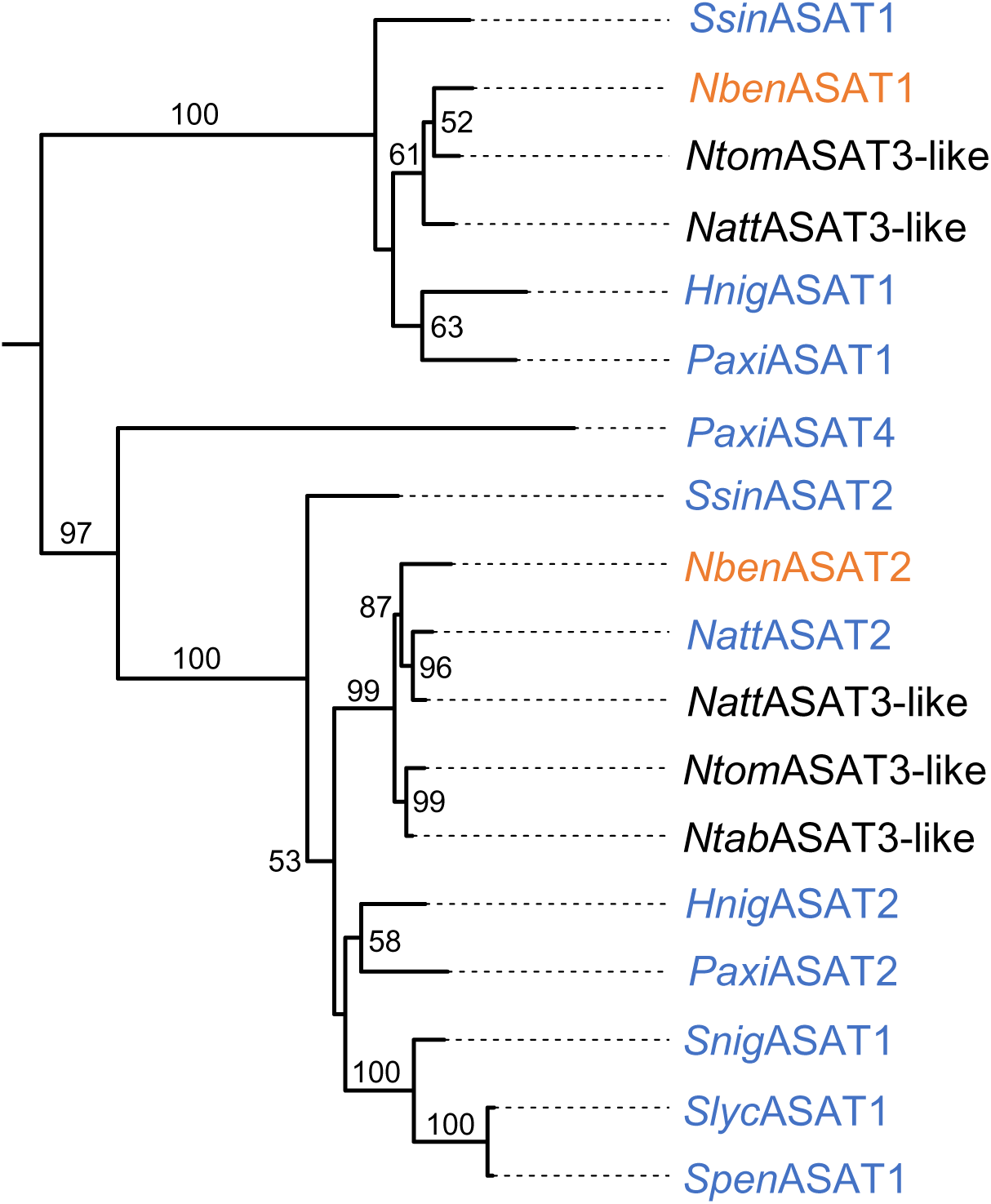
**ASAT phylogenetic tree** The evolutionary history of ASATs in the Solanaceae (Tables S1 and S2) was inferred using the Maximum Likelihood method in RAxML. Presented is a subtree of a larger tree that includes all annotated ASATs (Figure S1). The branch labels indicate the percentage of trees in which the associated taxa clustered together (bootstrap of 1000). Only values greater than 50 are presented. The two predicted *N. benthamiana* ASATs are highlighted in orange and ASATs that were previously chemically characterized are highlighted in blue.

### Generation of *ASAT* mutants

Using CRISPR/Cas9 coupled with tissue culture, we obtained two independent homozygous mutants for both *NbASAT1* and *NbASAT2*. *asat1-1* has a five-nucleotide deletion at the gRNA3 cutting site and a single nucleotide insertion at the gRNA2 cutting site, leading to a frameshift between gRNA3 and gRNA2. *asat1-2* has a 318-nucleotide deletion between the gRNA3 and gRNA2 cutting sites (Figure 3A). *asat2-1* has a single-nucleotide deletion at the gRNA3 cutting site and single-nucleotide insertion at the gRNA2 cutting site, leading to a translation frame shift between the two sites. *asat2-2* has a 115-nucleotide deletion at the gRNA3 cutting site and a single-nucleotide insertion at the gRNA2 cutting site (Figure 3B). Homozygous mutants were confirmed by DNA sequencing in the T2 generation, and this generation was used for all experiments.

**Figure 3.**
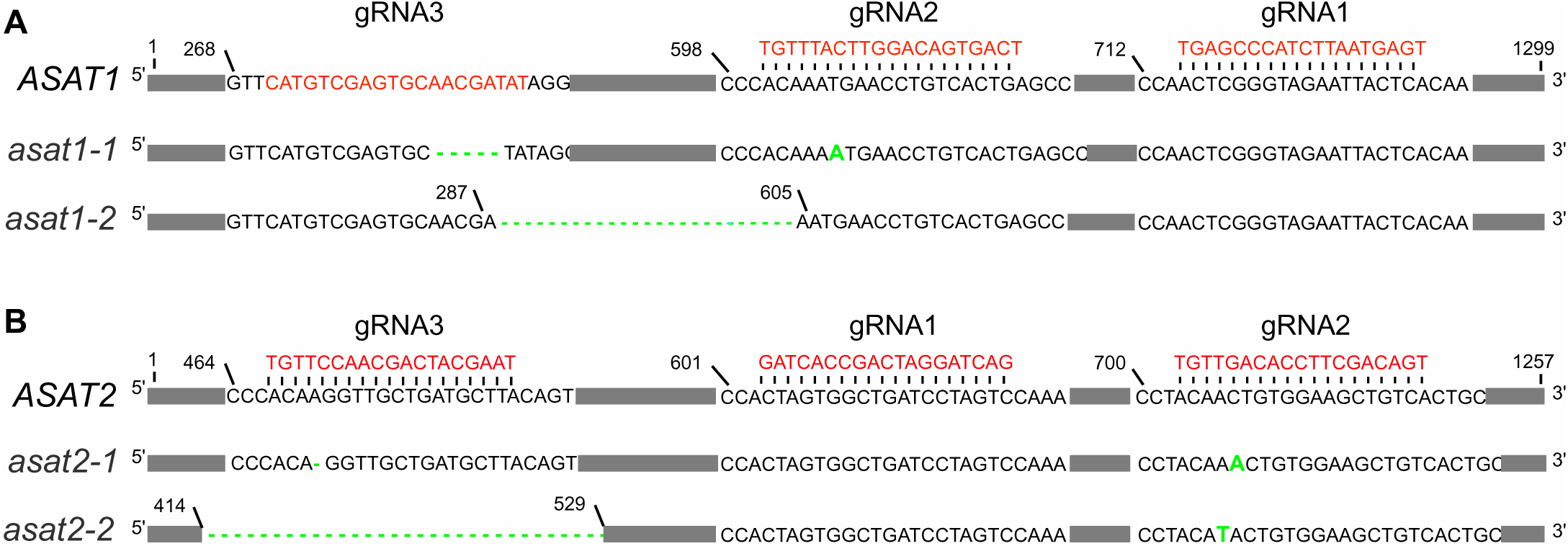
***N. benthamiana ASAT* mutations produced with CRISPR/Cas9** Three gRNAs (sequences shown in red) were designed to edit either *ASAT1* or *ASAT2*. Whereas gRNA3 for ASAT1 is on the sense strand, the other gRNAs are on the antisense strand. For both *ASAT1* and *ASAT2*, we obtained two independent mutations resulting from the corresponding gRNA2 and gRNA3. Single-base mutations and deletions are shown in green. (**A**) *asat1-1* has a five-nucleotide deletion at gRNA3 and a single-nucleotide insertion at gRNA2. *asat1-2* has a 318-nucleotide deletion between the gRNA3 and gRNA2 cutting sites. (**B**) *asat2-1* has a single-nucleotide deletion at gRNA3 and single-nucleotide insertion at gRNA2 leading to a translation frame shift between the two mutations. *asat2-2* has a 115-nucleotide deletion at gRNA3 and a single-nucleotide insertion leading to a translation frame shift at gRNA2.

### *ASAT2* knockout depletes acylsugar biosynthesis

We quantified the acylsugar content in the *ASAT* mutants by LC/MS, comparing to that of wildtype *N. benthamiana* plants. In the LC/MS profile of *N. benthamiana* leaf surface washes, we characterized twelve mass features as acylsucroses based on their characteristic peaks and neutral losses (Figure S3A). In negative electron spray ionization mode, the characteristic peak features included the mass of 341.11 for sucrose, 509.22 for sucrose + C2+C8, 467.21 for sucrose + C8, 495.21 for sucrose + C7, and 383.12 for sucrose + C2; the neutral loss peaks included mass for 126.10 for C8 (acyl-chain with 8 carbons), 129.09 for C7 + H_2_O, and 59.01 for C2 + H_2_O (Figure S3B). Those twelve *m/z* ratios included 383.12, 467. 21, 509.22, 555.23, 593.32, 621.31, M625.31, 635.32, 639.32, 667.32, 671.30, 681.34 (Figure S3A). Based on their MS/MS peak features, retention times and relative abundances, we predicted that the identified mass features are mainly derived from two acylsucroses as formate or chloride adducts, pathway intermediates, and/or resulted from in-source fragmentation. We named the two acylsucroses S3:17(2,7,8) and S3:18(2,8,8) (in the nomenclature, “S” refers to the sucrose backbone, “3:18” indicates three acyl chains with total eighteen carbons, and the length of each acyl chain is shown in parentheses) (Figure 1; Figure S3A).

In wildtype plants, S3:18(2,8,8) is the dominant acylsucrose, whereas S3:17(2,7,8) has relatively low abundance (Figure 4). S2:16(8,8) and S2:15(7,8), which may be biosynthetic pathway intermediates for S3:18(2,8,8) and S3:17(2,7,8), respectively, are present at lower levels (Figure 4). Compared to wildtype *N. benthamiana*, both *asat2-1* and *asat2-2* were almost completely depleted in both S3:17(2,7,8) and S3:18(2,8,8), as well as in the two predicted biosynthetic intermediates S2:15(7,8) and S2:16(8,8) (Figure 4). For *asat1-1* and *asat1-2*, the detected acylsucroses were less abundant, and significantly reduced in *asat1-1* (Figure 4). Although acylsugar content was significantly reduced in the *ASAT* mutants, the structure and abundance of trichomes on the leaf surface were not visibly changed (Figure S4).

**Figure 4.**
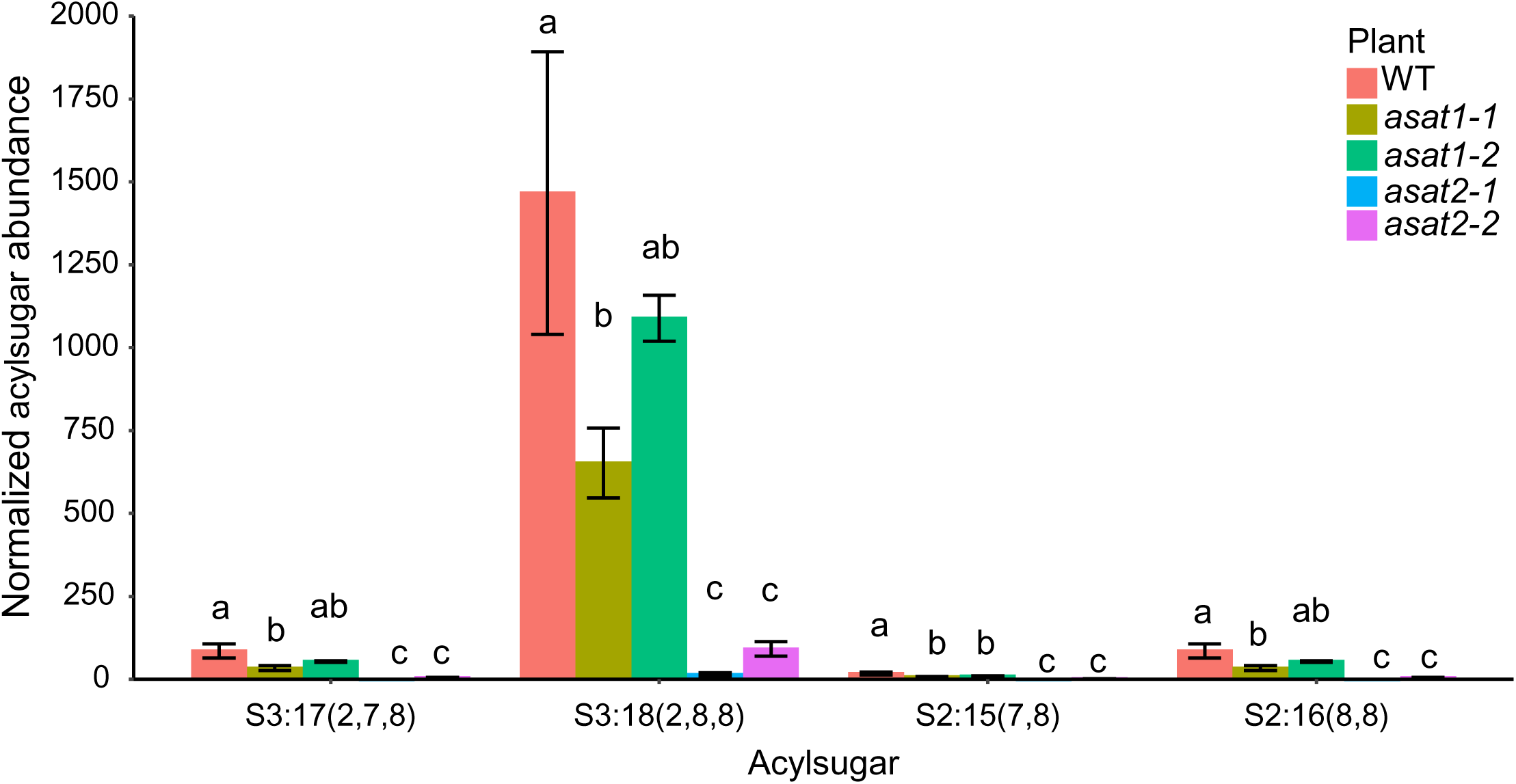
**The abundances of two *Nicotiana benthamiana* acylsugars (S2:17(2,7,8) and S2:18(2,8,8)), and two likely pathway intermediates/fragmentations (S2:15(7,8) and S2:16(8,8))** Acyl sugar LC/MS peak areas were normalized relative to the peak area of Telmisartan, which was added as an internal control, and then to the leaf dry weight (per gram). Error bars represent standard errors from measurements of three plants of each genotype. Significant differences for each acylsugar between different genotypes were tested using one-way ANOVA followed by a Duncan’s post hoc test (*p* < 0.05). Differences between groups are denoted with letters.

### Insect performance is improved on *ASAT2* mutant lines

We used *asat1* and *asat2* mutant lines to test the role of acylsugars in protecting *N. benthamiana* against insect pests. After placing synchronized first-instar aphids on mutant and wildtype *N. benthamiana* leaves, we monitored survival and growth over time. Significant improvements in aphid survivorship were observed as early as at 2 days post-feeding (*p* < 0.001) on the *asat2-1* and *asat2-2* mutants and increased until the end of the 5-day monitoring period (*p* < 0.001) (Figure 5A and Table S3). After 5 days of feeding, surviving aphids on both *asat1* and *asat2* plants were larger than those on wildtype plants (Figures 5B). When we measured progeny production by five adult aphids over a period of seven days, an average of more than 200 nymphs were produced on the *asat2* mutants, significantly more than the number of nymphs produced on either wildtype or *asat1* mutants (*p* < 0.05, Figure 5C). In aphid choice assays with detached leaves, they preferentially chose *asat2-1* mutant leaves relative to wildtype and *asat1-1* leaves (*p* < 0.05, Figures 5D-F). A preference for *asat2-1* and *asat2-2* leaves was consistently observed in choice assays involving any pairwise combination with wildtype, *asat1-1*, and *asat1-2* leaves (*p* < 0.001, *Chi-square* test, Figures 5D-F and S5). No *M. persicae* preference was observed when comparing wildtype *N. benthamiana* and *asat1* mutants (*p* > 0.05, Figures 5D-F and S5).

**Figure 5.**
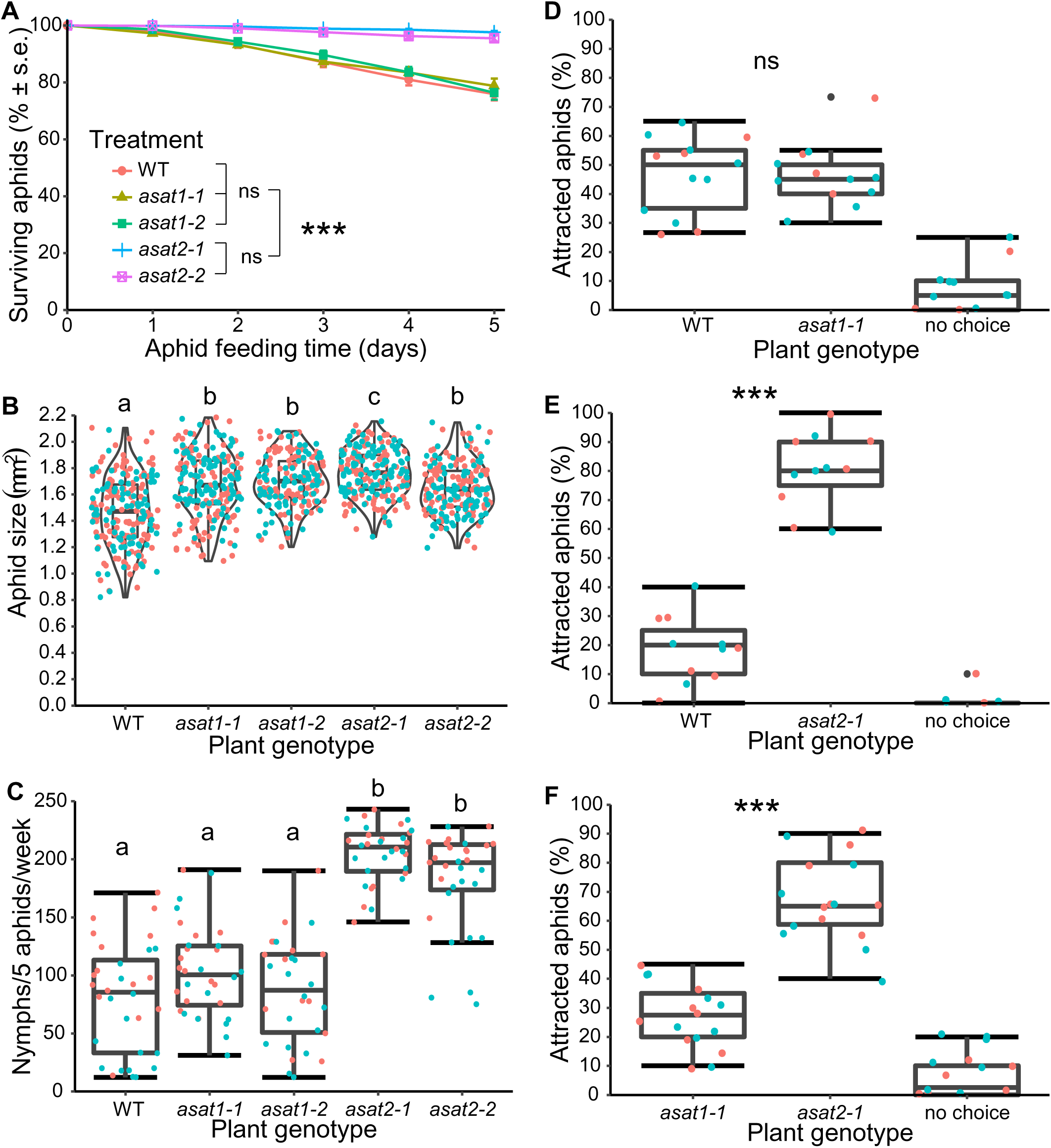
***Myzus persicae* bioassays with wildtype (WT), *asat1,* and *asat2 Nicotiana benthamiana*** Data shown in all panels were combined from two independent experiments, which are shown in color orange and cyan in panels B-F. **(A)** Survival over 5 days of nymphs placed onto mutant and wildtype *N. benthamiana. S*ignificant differences are indicated for the 5-day time point: ns: not significant, *** *p* < 0.001, mean +/- s.e. of n = 15-16. Significant differences were tested using one-way ANOVA with a fixed factor of genotypes and a block effect of experiment followed by a Bonferroni post hoc test for multiple comparisons. Full statistical data for all time points are in Table S3. **(B)** Aphid growth, as measured by body size after 5 days feeding on *N. benthamiana*. **(C)** Aphid reproduction, as measured by the number of nymphs that were produced by five aphids in one week. Significant differences between different groups (*p* < 0.05) were determined using ANOVA with a fixed factor of genotypes and a block effect of experiment followed by a Duncan’s post hoc test and are indicated by lowercase letters above each group in panels B and C. **(D-F)** Aphid choice among detached leaves of each plant genotype, significant differences between genotypes were tested using Chi-square test, ns: not significant, *** *p* < 0.001. no choice: aphids were elsewhere in the Petri dish and not on a leaf. The box plots show the median, interquartile range, maximum and minimum after removal of outliers, and the individual data points.

When aphid colonies were allowed to grow long-term on *asat2-1* mutant and wildtype *N. benthamiana* in the same growth chamber, there were many more aphids on the mutant plants (Figure S6A,B), likely resulting from a combination of host plant choice and increase growth on the *asat2-1* mutant. It is noteworthy that, on the *asat2-1* mutant plants, aphids were feeding on the more nutritious younger leaves, which tend to be better-defended in plants. By contrast, on wildtype *N. benthamiana* aphids were only were able to feed on older, senescing leaves and were primarily on the abaxial surface. Consistent with the increased aphid presence, growth of the *asat2-1* mutant plants was visibly reduced relative to wildtype *N. benthamiana* (Figure S6B). Given the almost identical phenotypes of *asat2-1* and *asat2-2* mutants, subsequent insect assays were conducted with T2 progeny of the *asat2-1* line, which also were confirmed by PCR to contain the Cas9 transgene.

As we observed with *M. persicae*, the *asat2-1* mutation improved the *M. euphorbiae* performance on *N. benthamiana* (Figure 6A-C). Significantly increased *M. euphorbiae* survival was observed after 24 hours on *asat2-1* compared to wildtype (*p* < 0.001, Figure 6A). Additionally, significantly more nymphs were produced by adult *M. euphorbiae* in the course of 24 hours on *asat2-1* than on wildtype (*p* < 0.05, Figure 6B). In choice assays, potato aphids preferentially chose the *asat2-1* leaves over wildtype leaves (*p* < 0.001, Figure 6C). Whereas we were not able to establish an *M. euphorbiae* colony on wildtype *N. benthamiana*, the aphids readily formed colonies on the *asat2-1* mutant plants (Figure 6D).

**Figure 6.**
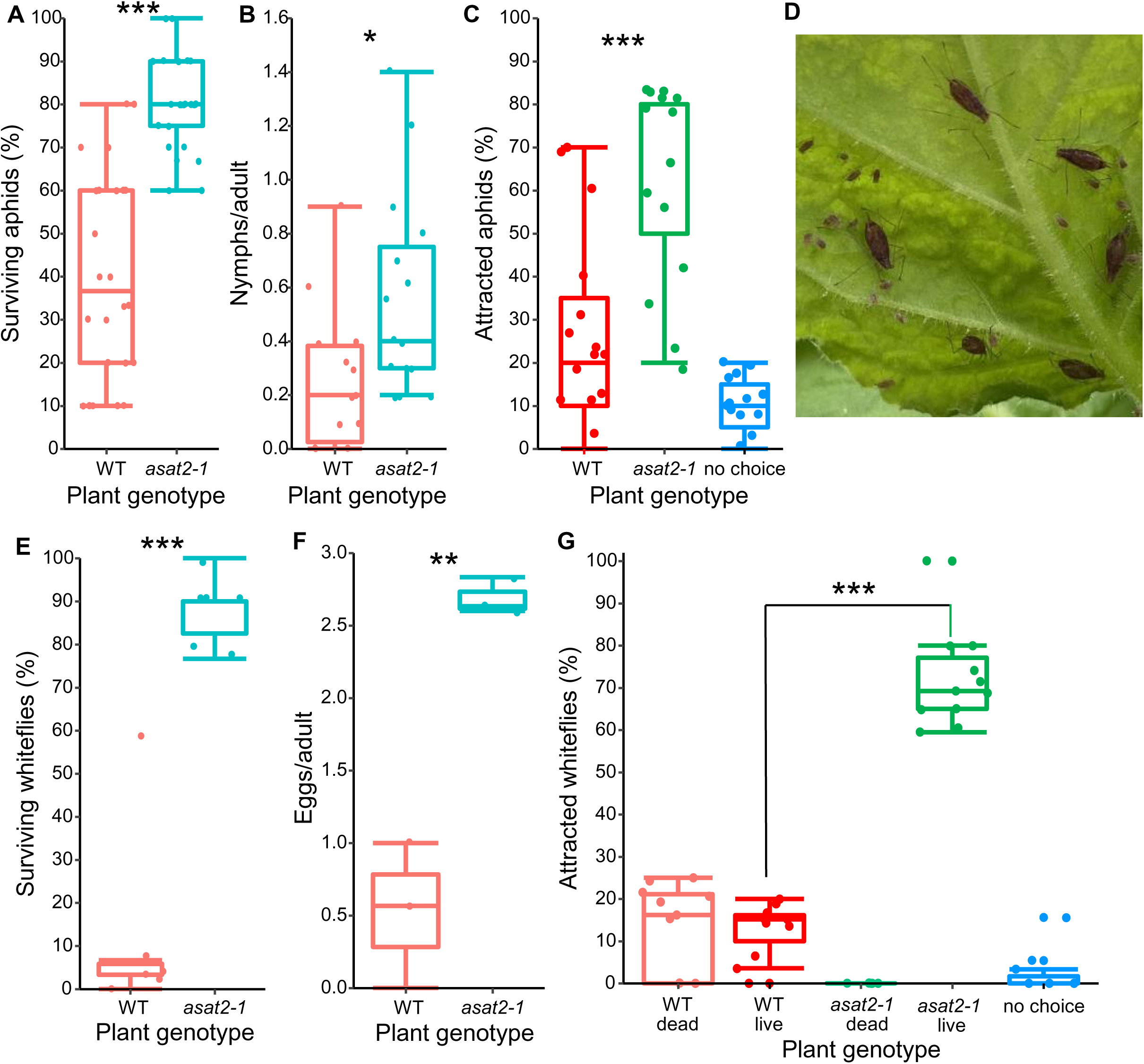
**Potato aphid and whitefly bioassays on *asat2-1* and wildtype (WT) *N. benthamiana.*** (**A-D**) potato aphid (*M. euphorbiae*) bioassays. (**A)** aphid survival in 24 hours (n=15). (**B)** aphid reproduction in 24 hours (n=15). (**C**) aphid choices between detached leaves of each plant genotype (n=15). (**D**) An established *M. euphorbiae* colony on an *N. benthamiana asat2-1* leaf. (**E-G**) whitefly bioassays. (**E)** whitefly survival in 3 days (n=6). (**F)** whitefly reproduction measured as number of eggs produced per adults in 3 days (n=3). (**G)** whitefly choices between plants of each genotype (n=3 for 4 experiments). Significantly differences were tested using independent t-test for aphid and whitefly survival and reproduction data, and Chi-square test was used for aphid and whitefly choice assays. * *p* < 0.05, ** *p* < 0.01, *** *p* < 0.001. no choice: insects were not on a leaf at the end of the experiment. The box plots show the median, interquartile range, maximum and minimum after removal of outliers, and the individual data points.

Survival of *B. tabaci* was greatly increased on the *asat2-1* mutant line relative to wildtype (Figure 6E). Moreover, whiteflies laid significantly fewer eggs on wildtype *N. benthamiana* than on *asat2-1* mutant plants over three days (Figure 6F). Dead adult whiteflies were observed on wildtype plants (Figure S7A), and it was not possible to establish a reproducing colony. By contrast, after 23 days of feeding on *N. benthamiana*, whiteflies of different life stages were observed on *asat2-1* mutant plants (Figure S7B-D). In choice assays with mutant and wildtype plants in the same cage, *B. tabaci* preferentially settled on *asat2-1* plants (72%) over wildtype (26%) in a 24-hour experiment (Figure 6G). Notably, all whiteflies on *asat2-1* were alive after 24 hours, whereas about half of the whiteflies found on the wildtype plants were dead at the same time point (Figure 6F).

To determine whether depletion of acylsugars in *ASAT2* mutants improves the performance of lepidopteran herbivores on *N. benthamiana*, we conducted experiments with *Helicoverpa zea* (corn earworm), *Heliothis virescens* (tobacco budworm) and *Trichoplusia ni* (cabbage looper). When neonates were placed on the leaves of wildtype or *asat2-1* mutant *N. benthamiana,* no *H. zea* caterpillars were recovered (Figure 7A). Survivorship of H*. virescens* and *T. ni* larvae on *N. benthamiana* was low, and the mass of the surviving larvae after ten days was not significantly increased on the mutant line relative to wildtype (Figure 7B,C). Due to the low survival of neonates, we repeated the caterpillar bioassay using five-day-old larvae that had been reared on artificial diet. Almost all *H. zea* and *H. virescens* larvae survived for seven days on wildtype and *asat2-1* mutant plants, and survival of *T. ni* caterpillars was higher on *asat2-1* than on wildtype plants (Figure 7D-F). The relative growth rates of surviving *H. zea*, *H. virescens*, and *T. ni* larvae were higher on the *asat2-1* mutant by 35%, 47%, and 99%, respectively, than on wildtype plants (Figure 7D-F).

**Figure 7.**
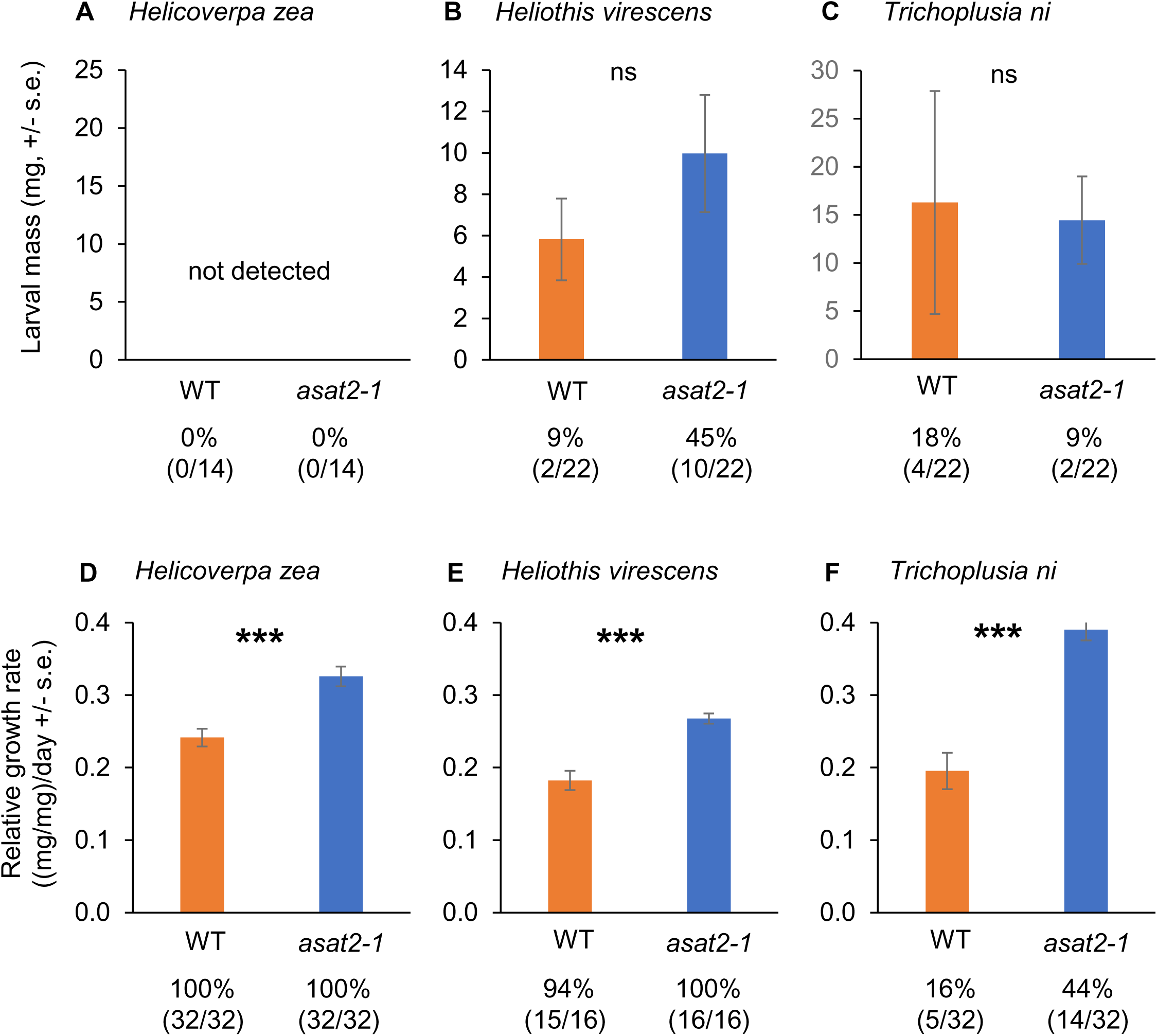
**Caterpillar bioassays on wildtype (WT) and *asat2-1* mutant *Nicotiana benthamiana.*** (**A, B, C**) Larval mass of surviving *Helicoverpa zea*, *Heliothis virescens*, and *Trichoplusia ni* after 10 days after being placed on plants as neonates. (**D, E, F**) Relative growth rate of surviving *H. zea*, *virescens*, and *T. ni* on wildtype and *nata2-1* plants. Insects were raised for five days on artificial diet, prior to 7 days of feeding on *N. benthamiana*. Percent survival (number of surviving insects/number of total insects) is shown below each figure. Mean +/- s.e. ,\*\*\**p* < 0.001, ns: no significant difference (P > 0.05, *t-*test).

### Virus transmission by aphids is increased on *asat2-1* mutant plants

To demonstrate the use of *asat2-1* mutants for aphid experiments, we measured *M. persicae* transmission of a GFP-expressing turnip mosaic virus (TuMV-GFP; Lellis et al., 2002; Casteel et al., 2015) using all combinations of wildtype and *asat2-1* plants as virus donors and recipients, respectively (Figure 8). The earliest GFP signals in the recipient plants were visible under UV light on day 4 after TuMV-GFP transmission. Successful transmission and virus replication was monitored until day 9, when no plants developed additional visible TuMV-GFP infections relative to the previous day (Figure 8). The *asat2-1* → *asat2-1* transmission group showed the highest final TuMV-GFP infection rate (73%), significantly higher than that observed in the other three groups (*p* < 0.05, ANOVA). Relative to the wildtype → wildtype TuMV-GFP transmission, transmission was not significantly increased in the *asat2*-*1* → wildtype and wildtype → *asat2-1* groups. The significantly higher virus transmission rate by aphids in the *asat2*-*1* → *asat2-1* group is consistent with the feeding preference of *M. persicae* for *asat2-1* mutant plants (Figure 5).

**Figure 8.**
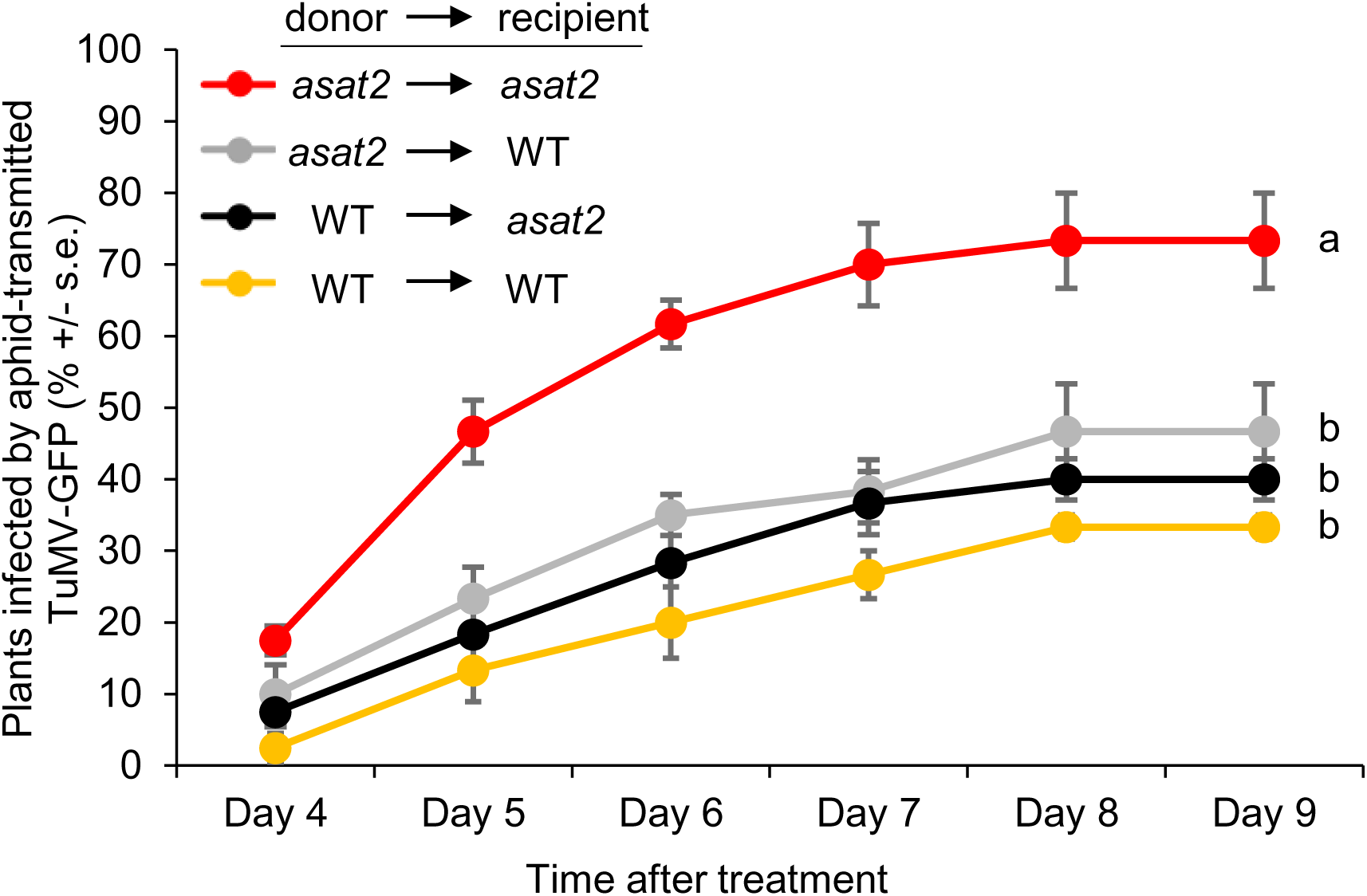
**Aphid transmission of GFP-expressing turnip mosaic virus (TuMV-GFP) on wildtype (WT) and *asat2-1* plants.** Aphids were transferred from *N. benthamiana* plants infected with TuMV-GFP to uninfected plants and the percentage of GFP-expressing recipient plants was recorded over time. Data are mean +/- s.e. of N = 60, with data from three separate experiments combined. Statistical analyses were conducted using one-way ANOVA followed by a Tukey HSD post hoc test. Significant differences were observed from day 5 post treatment and thereafter. Lowercase letters denoted the differences between each group at day 9 post treatment (*p* < 0.05). WT: wildtype.

### The *asat2-1* mutant is suitable for plant-mediated VIGS in whiteflies

To demonstrate the utility of the *asat2-1* mutant for whitefly experiments, we performed plant-mediated virus-induced gene silencing (VIGS) using tobacco rattle virus (TRV; Hayward et al., 2011). We selected two previously validated *B. tabaci* RNAi targets, *AchE* (*acetylcholinesterase*; Malik et al., 2016) and *TLR7* (*toll-like receptor 7*; Chen et al., 2015), as well as two predicted horizontally transferred genes (Chen et al., 2016), *SQS* (*squalene synthase*) and *TSCD* (*tubulin-specific chaperone D*), that had not been previously investigated in whiteflies. When whiteflies were place on *N. benthamiana* infected with TRV VIGS constructs, expression of all four target genes was significantly reduced after one day of feeding (Figure 9A). After seven days of feeding, the expression of *AchE* and *SQS* was significantly reduced relative to whiteflies on empty vector control plants, but expression of *LTR7* and *TCSD* was not (Figure 9B). After seven days, survivorship of the whiteflies on VIGS plants was significantly reduced relative to control plants with empty vector or GFP control TRV infections (Figure 9C). Consistent with the low survival observed in previous experiments (Figure 6E), whitefly survival on TRV-infected wildtype *N. benthamiana* was very low (Figure 9D), never exceeding 10%. None of the VIGS constructs decreased survival to a lower level than on the empty vector and TRV-GFP control plants.

**Figure 9.**
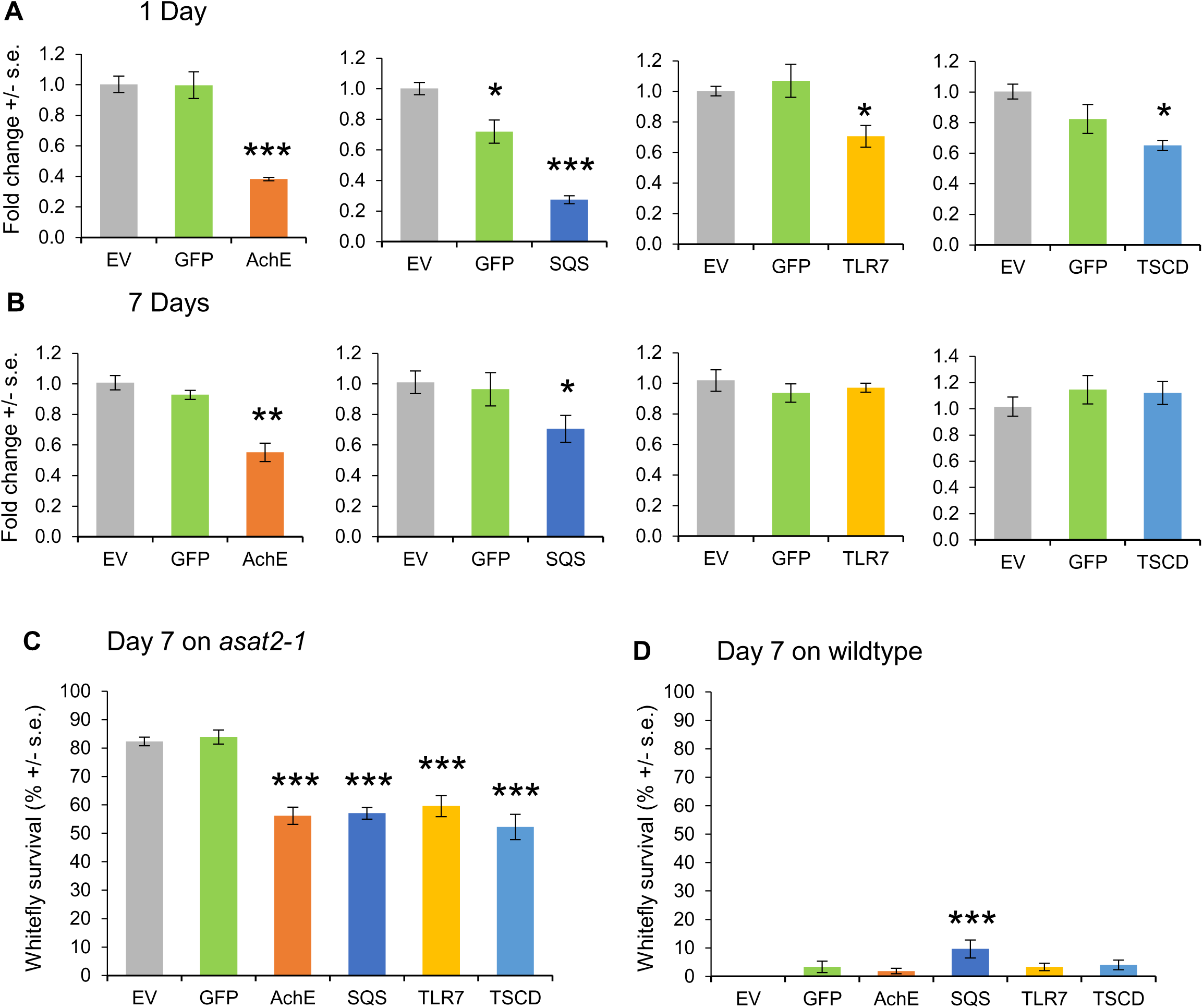
**Virus induced gene silencing (VIGS) of whitefly genes using tobacco rattle virus.** Whiteflies were placed on plants infected with TRV expressing VIGS constructs and gene expression was measured after (**A**) one and (**B**) seven days. In each experiment, the empty virus vector (EV) and GFP-carrying virus were used as controls. Gene expression was normalized to 1 for EV control plants. Whitefly survival was assessed on TRV-infected plants after (**C**) seven days of feeding on *asat2-1* plants, and (**D**) seven days of feeding on wildtype plants. GFP/AchE/SQS/TLR7/TSCD: VIGS plants with TRV expressing RNA constructs targeting GFP, AchE, SQS, TLR7, and TSCD, respectively; AchE: acetylcholinesterase, SQS: squalene synthase, TLR7: toll-like receptor 7, TSCD: tubulin-specific chaperon D. The survival and the 7-day qPCR experiments were done for three times. The qPCR experiments for 1-day were done twice. Significant differences were determined using one-way ANOVA with a fixed factor of treatments and a block effect of experiment, followed by a Dunnett’s post hoc test for comparing VIGS constructs to the empty vector (EV) control. Mean +/- s.e. of n = 3 (A), n = 9 (B), and n = 27 (C,D). \**p* < 0.05; \*\**p* < 0.01, \*\*\**p* < 0.001.

### Water loss is greater in *asat2* mutant plants that in wildtype

While conducting aphid choice assays with detached leaves (Figures 5D-F and 6C), we noticed that the mutant leaves dried out faster than wildtype leaves. This effect was quantified using detached-leaf assays, in which *asat2* leaves lost significantly more water over 24 hours than leaves from either wildtype or *asat1* leaves (Figures 10A and S8A). When plants are subjected to drought, high/low temperature, salinity, herbivores, or other stresses, they absorb and reflect specific wavelengths of light, which have been used to determine vegetation indices. For example, the water band index (WBI) (Penuelas et al., 1993) has been used to monitor changes in the plant canopy/leaf water content. Using hyperspectral imaging, we determined that the leaf water content of intact plants, as measured by the WBI, was significantly lower in *asat2* mutants than in wildtype (Figures 10B and S8B). Although the *asat1* mutants did not lose water faster than wildtype in detached leaf assays (Figure 10A), the leaf water content in *asat1* mutants was significantly lower than wildtype (Figures 10B and S10B). Measurement of leaf temperature by thermal imaging showed that, consistent with the reduced leaf water content, the leaf temperature of the acylsugar mutants was significantly higher than that of wildtype plants (Figures 10C and S10C).

**Figure 10.**
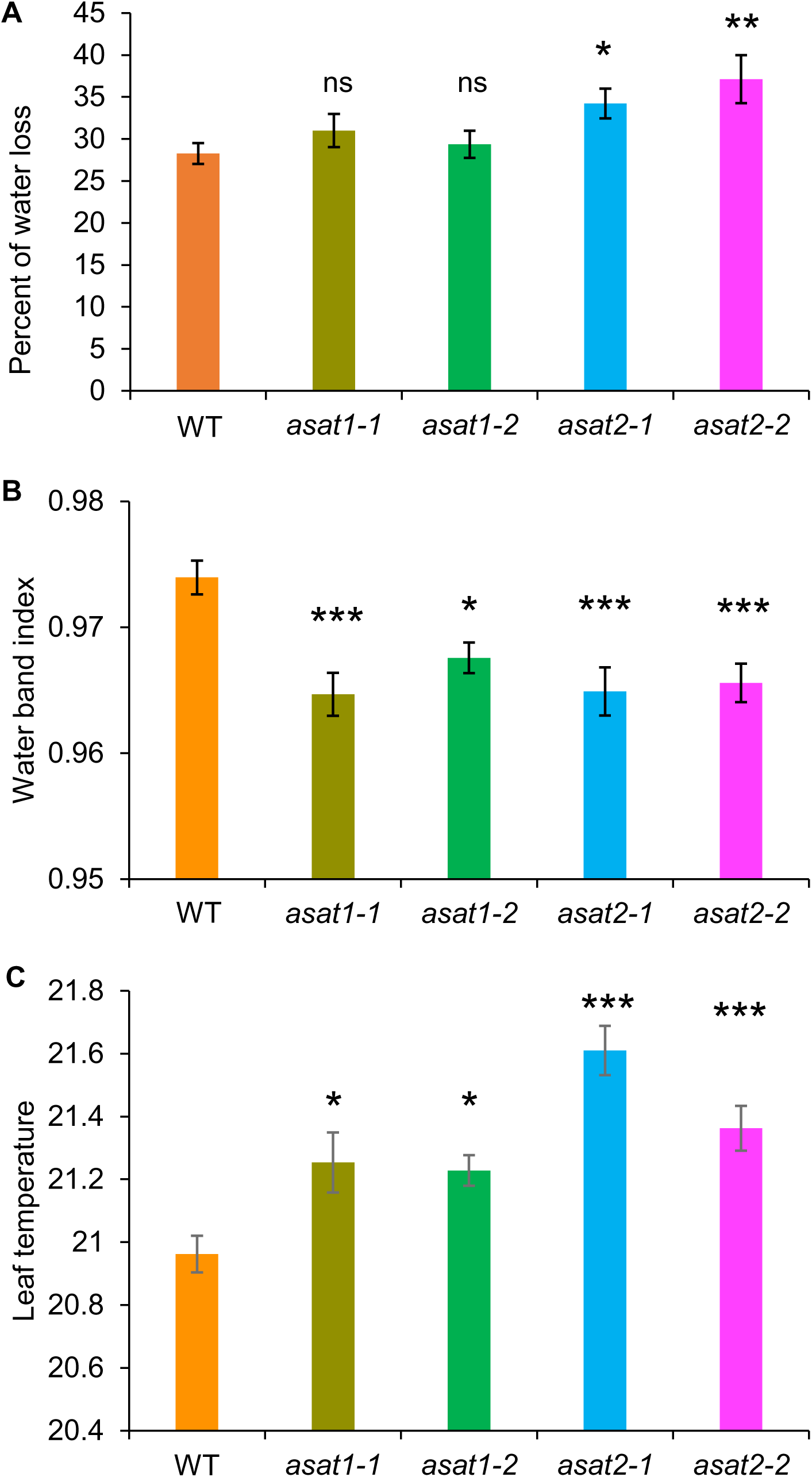
**Water loss and leaf temperature of wildtype (WT), *asat1*, and *asat2 Nicotiana benthamiana.* (A)** Percent of water loss from detached leaves in 24 hours, mean +/- s.e. of n = 15. (**B**) Leaf water content measured by the water band index from hyperspectral imaging, mean +/- s.e. of n = 20. (**C**) Leaf temperatures from leaves of different plant genotypes, mean +/- s.e. of n = 20. ns = not significant, \**p* < 0.05, \*\**p* < 0.01, \*\*\**p* < 0.001, Dunnett’s test relative to wildtype control.

## Discussion

We were able to identify only two *ASAT* genes, *NbASAT1* and *NbASAT2*, as well as a fragmented pseudogene, in the *N. benthamiana* genome (Table S2). By contrast, in other *Nicotiana* species, there are larger numbers of predicted *ASATs*, *e.g.* one *ASAT1*, one *ASAT2*, and 20 *ASAT3*-like genes in *N. attenuata* (Gaquerel et al., 2013; Van et al., 2017), 35 *ASAT3*-like genes in *N. tabacum*, and 19 *ASAT3*-like genes in *N. tomentosiformis* (Egan et al., 2019). Given the relatively small number of predicted *ASAT* genes in *N. benthamiana*, other enzymes may also be involved in acylsugar biosynthesis. In the Solanaceae, two common pathways are known for aliphatic acid elongation via acetate: the fatty acid synthase (FAS) and the alpha-ketoacid elongation (αKAE) pathways (Kroumova and Wagner, 2003). In the FAS pathway, two carbons from an acetyl-acyl carrier are retained per elongation cycle, whereas in the αKAE pathway, one carbon is retained per elongation cycle (Kroumova and Wagner, 2003). Knockdown of *E1-*β *branched-chain* α *keto acid dehydrogenase* (*BCKD*) significantly reduces acylsugars in *N. benthamiana* (Slocombe et al., 2008). Additionally, *Isopropylmalate Synthase 3* (*IPMS3*) in cultivated and wild tomatoes (Ning et al., 2015) and *Acyl-Sucrose Fructo-Furanosidase 1* (*ASFF1*) in wild tomato (Leong et al., 2019) are involved in determining acylsugar composition. Further studies will be needed to characterize other genes involved in the *N. benthamiana* acylsugar biosynthesis pathway.

Acylsugars can be categorized as sucrose or glucose esters based on the sugar cores, which are decorated with varying numbers or lengths of acyl chains (Kim et al., 2012). Whereas some wild tomatoes produce a mixture of acylsucroses and acylglucoses, we observed only acylsucroses (Figures 1 and 4), consistent with previous identification of these compounds in *N. benthamiana* (Matsuzaki et al., 1989; Matsuzaki et al., 1992; Hagimori et al., 1993; Slocombe et al., 2008). Nevertheless, it has been reported that *N. benthamiana* produces acylglucoses, although in lower abundance than acylsucroses (Hagimori et al., 1993), and one glucose ester structure has been proposed (Matsuzaki et al., 1992). Our failure to detect acylglucoses may be explained by the use of different isolates of *N. benthamiana*, growth conditions, growth stage of plants, and/or the detection methods. Whereas we used ∼1-month-old plants and LC/MS for our assays, Matsuzaki et al. (1992) used ∼3-month-old plants and GC/MS to detect acylglucoses in *N. benthamiana*.

Abundance of the characterized acylsugars was reduced to a greater extent in *N. benthamiana asat2* than in *asat1* mutants (Figure 4). The smaller reduction of acylsugars in *asat1* mutants suggests that either *Nb*ASAT2 functions upstream of *Nb*ASAT1 in the acylsugar biosynthesis pathway, but partially complements *Nb*ASAT1 activity; or *Nb*ASAT1 and *Nb*ASAT2 have similar functions in the biochemical pathway, but *NbASAT2* had higher abundance or enzymatic activity. It is not known whether *N. benthamiana* ASATs are monomeric or multimeric, but BAHD acyltransferases generally are monomeric enzymes (D’Auria, 2006), suggesting that heterodimers between ASAT1 and ASAT2 are unlikely to affect the observed phenotypes.

In the ASAT phylogenetic tree (Figure 2), *Nb*ASAT2 is closely related to some biochemically characterized ASAT1 proteins in other Solanaceae species, including the *Sp*ASAT1, *Sl*ASAT1, and *Sn*ASAT1. Those ASAT1s have some substrate overlap with the ASAT2s found in the corresponding species, indicating that ASAT2 has moved toward utilizing the ASAT1 substrate in these species over time (Moghe et al., 2017). The final activity shift that has become fixed in the *Solanum* genus, most likely occurred after the divergence of the *Solanum* and *Capsicum* clades (Moghe et al., 2017). However, if our hypothesis of partial complementation of *Nb*ASAT1 by *Nb*ASAT2 is correct, it may flag a transition stage or suggest independent *Nicotiana*-specific evolution of the ASAT1 and ASAT2 functions. Based on previous knowledge of BAHD activities (Moghe et al., 2017), we postulate that S2:15 (7,8) and S2:16 (8,8) are produced by *Nb*ASAT1 and *Nb*ASAT2, whereas the acetylation is carried out by another unrelated BAHD enzyme – not unlike the distantly related *Sl*ASAT4 and *Salpiglossis sinuata* ASAT5 enzymes (Schilmiller et al., 2015; Moghe et al., 2017). Further characterization will be required to identify specific acyltransferase enzyme activities in *N. benthamiana*.

The observed role of acylsugars in protecting against desiccation (Figures 10 and S8) is consistent with previous reports from other Solanaceae (Fobes et al., 1985; O’ Connell et al., 2007; Kroumova et al., 2016) and is likely an adaptation to the seasonally arid native habitat of *N. benthamiana* in northwestern Australia (Goodin et al., 2008; The Australasian Virtual Herbarium, https://avh.ala.org.au). Relative to *asat1-1* and *asat1-2* mutants, the lower acylsugar content of *asat2-1* and *asat2-2* mutants (Figure 4), resulted in more rapid water loss in detached leaves (Figures 10A and S8A). However, despite the only partial decrease in the acylsugar content of *asat1* mutants, the decreases in water content and increases in leaf temperature of intact plants were similar to those of *asat2* mutants (Figures 10B,C and S8B,C).

Acylsugars with C_7-12_ chains have been shown to be the most toxic sugar esters for small phloem-feeding Hemiptera such as aphids, Asian citrus psyllids, and whiteflies (Chortyk et al., 1996; McKenzie and Puterka, 2004; Song et al., 2006). Synthetic acylsucroses with di-heptanoic acid (C7), di-octanoic acid (C8), and di-nonanoic acid (C9) acyl groups showed the highest mortality in bioassays with *M. persicae* and *B. tabaci* (Chortyk et al., 1996; McKenzie and Puterka, 2004; Song et al., 2006). *Nicotiana gossei*, a tobacco species that produces mainly C7-C8 acyl group acylsugars, has a high level of insect resistance relative to close relatives with acylsugar profiles that are not dominated by those with C7-C12 acyl groups (Thurston, 1961; Kroumova and Wagner, 2003). In *N. benthamiana*, the two most abundant acylsugars that we found contain C7 and predominantly C8 acyl groups, which is consistent with previous findings of mainly 5- and 6-methyl heptanoate (C8) in *N. benthamiana* (Kroumova and Wagner, 2003; Slocombe et al., 2008) and *N. alata* (Moghe et al., 2017). The almost complete depletion of acylsugars in our *asat2* mutants improved both hemipteran and lepidopteran performance, suggesting that the identified C8-chain acyl group acylsugars are providing insect resistance for *N. benthamiana*.

We cannot rule out the possibility secondary effects that might also influence insect performance on acylsugar-depleted *N. benthamiana*. Specialized metabolites in other plants, for instance glucosinolates in *Arabidopsis thaliana* (Clay et al., 2009) and benzoxazinoids in *Zea mays* (Meihls et al., 2013), regulate callose deposition as a secondary defense response. It is not known whether acylsugars contribute to the regulation of other defense responses in *N. benthamiana.* The observation of numerous dead whiteflies on wildtype *N. benthamiana* plants in choice assays (Figure 6G), despite the option of moving to presumably more desirable *asat2-1* mutant plants in the same cage, suggests that the acylsugars stickiness also plays a role in plant defense by immobilizing the insects. Both altered leaf turgor and leaf temperature (Figure 10B,C) could affect insect feeding behavior and growth rate, though the specific effects on the six tested insect species cannot be determined without further research.

*Nicotiana benthamiana* has been used for virus transmission assays, RNA interference and VIGS of *M. persicae* genes, and transient expression of genes from other species to determine their effects on aphid growth and reproduction (Ramsey et al., 2007; Bos et al., 2010; Pitino and Hogenhout, 2013; Casteel et al., 2014; Elzinga et al., 2014; Rodriguez et al., 2014; Krenz et al., 2015; Tzin et al., 2015; Mulot et al., 2016; Mathers et al., 2017; Del Toro et al, 2018; Cui et al., 2019; Worrall et al., 2019). However, the poor growth of many *M. persicae* isolates on *N. benthamiana* (Thurston, 1961; Hagimori et al., 1993; Figure S6) hampers experiments of this kind Similarly, due to the very poor survival of *B. tabaci* on wildtype plants (Simon et al., 2003; Figure 6E), *N. benthamiana* has not been useful as a host plant for whitefly research.

To demonstrate the utility of *asat2* mutants as a tool for aphid research, we performed TuMV-GFP transmission experiments using *M. persicae*. Our results show that *asat2-1* plants significantly increased virus transmission (Figure 8). Interestingly, efficient virus transmission only occurred when both the virus donor and the recipient plants had the *asat2-1* genotype. This suggests that aphid behavior on *asat2-1* plants, perhaps faster or more continuous probing, promotes both uptake and delivery of TuMV. The higher virus transmission on *asat2-1* mutant *N. benthamiana* will facilitate use of this model system for future research on plant-aphid-virus interactions and factors that promote virus transmission.

The low survival of whiteflies on wildtype *N. benthamiana* (Figures 6E and 9D) makes it difficult to assess the negative effects of gene expression silencing by TRV VIGS. By contrast, we were able to demonstrate reduced survival with all four tested whitefly genes with TRV VIGS using *asat2-1* mutant plants (Figure 9C). *TLR7* and *TSCD* gene expression was reduced on day 1 but not on day 7 in the VIGS experiment (Figure 10A,B), but nevertheless there was a negative effect on whitefly survival over 7 days (Figure 10C). Two possible explanations for this finding are: (i) There was a survivor bias in that we could only measure gene expression levels in surviving aphids, and perhaps all aphids with efficient expression silencing of *TLR7* and *TSCD* were dead after 7 days. (ii) There may be gene expression compensation at the whole-insect level over time, but not in specific tissues that affect insect survival. Quantitative PCR of fractionated whiteflies would be necessary to determine the time course of *TLR7* and *TSCD* expression silencing and whether VIGS primarily affects gene expression in specific tissue types.

In addition to confirming the importance of two *B. tabaci* genes, *AchE* and *TLR7*, that have been targeted by RNAi (Malik et al., 2016; Chen et al., 2015), we chose two previously uncharacterized horizontally transferred genes *SQS* and *TSCD,* as VIGS targets. *Bemisia tabaci* MEAM1 has at least 142 horizontally transferred genes from bacteria and fungi (Chen et al., 2016). Given that horizontal transfer of functionally expressed microbial genes into insect germlines is rare on an evolutionary timescale, there is likely a selective advantage to having these genes expressed in whiteflies. This was confirmed by the observation that VIGS of both *SQS* and *TSCD* reduced whitefly survival relative to control plants (Figure 9C,D). Transient expression knockdown of *SQS*, *TSCD*, and other horizontally transferred genes will enable future research to study the functions of these genes in whitefly metabolism. Due to their importance for whitefly survival, as well their absence in beneficial insects such as ladybugs and lacewings, horizontally transferred genes also are attractive targets for controlling whiteflies by RNA interference. The established *N. benthamiana* VIGS system, which allows rapid cloning of targets by Gateway recombination (Liu et al., 2002), will allow rapid screening of other horizontally transferred genes to identify ones that would be most suitable for whitefly control on crop plants by RNA interference.

Although *H. zea*, *H. virescens*, and *T. ni* larvae grow well on cultivated tobacco, neonate larvae had a low survival rate on both wildtype *asat2-1 N. benthamiana* (Figure 7A-C). There was a higher survival rate with five-day-old larvae of the three tested species, which all grew significantly better on *asat2-1* mutants than on wildtype *N. benthamiana* (Figure 7D-F). Thus, *N. benthamiana* acylsugars likely provide at least some protection against lepidopteran pests. However, the high mortality of neonate larvae on *asat2-1* plants suggests that either residual acylsugars or as yet unknown resistance mechanisms in *N. benthamiana* can provide protection. Additional mutations that decrease insect resistance, perhaps regulatory genes such as *COI1* or genes affecting the production of other specialized metabolites, will be necessary to facilitate *N. benthamiana* experiments with *H. zea*, *H. virescens*, *T. ni*, and other commonly studied lepidopteran species.

The *Cas9* transgene, which is still present in the *asat2-1* mutant line that we used for most of our experiments, may facilitate further mutagenesis. In a recently described method (Ellison et al., 2020) *Cas9*-transgenic *N. benthamiana* plants were infected with TRV carrying gRNAs linked to phloem movement sequences. Seeds harvested from these plants had a high frequency of homozygous knockout mutations in the CRISPR/Cas9-targeted genes. The ease of generating germline mutations using this approach will make it possible to test the function of other predicted insect defense genes by knocking out their expression in the *N. benthamiana asat2-1* mutant background using a TRV-expressed gRNA.

Our knockout of acylsugar biosynthesis is an important first step toward improving the already excellent *N. benthamiana* model system (Goodin et al., 2008; Bally et al., 2018), making it more suitable for studying plant interactions with *M. persicae*, *B. tabaci*, and other agriculturally relevant insect pests. Such experiments can include transient expression assays to test the function of insect elicitors and insect-defensive genes from other plant species in *N. benthamiana*, as well as VIGS to down-regulate insect gene expression in a targeted manner. Furthermore, the almost complete absence of acylsugars in the *asat2* mutant lines, in combination with the facile *Agrobacterium-* and virus-mediated transient gene expression systems available for *N. benthamiana*, will make these mutants a suitable platform for the functional analysis of ASATs from other Solanaceae.

## Materials and Methods

### Insect and plant cultures

A red strain of *M. persicae* (Ramsey et al., 2007; Ramsey et al., 2014), originally collected from *N. tabacum* by Stewart Gray (Robert W. Holley Center for Agriculture & Health, Ithaca, NY), was maintained on *N. tabacum* plants in a growth room at 23°C with a 16:8 h light:dark photoperiod. Insect bioassays were conducted in the same growth room. Colonies of *B. tabaci* MEAM1 were provided by Danielle Preston and Angela Douglas (Cornell University) and Jane Polston (University of Florida). *Macrosiphum euphorbiae* was obtained from Isgouhi Kaloshian (UC Riverside) and was maintained on tomato cv. Moneymaker. Eggs of *H. zea*, *H. virescens*, and *T. ni* were purchased from Benzon Research (www.benzonresearch.com). *Nicotiana benthamiana* wild type and mutant plants for aphid experiments, caterpillar experiments, and whitefly VIGS assays were maintained at 23°C and a 16:8 h light:dark photoperiod in a Conviron (Winnipeg, Canada) growth chamber and, for seed production, in a greenhouse at 27/24°C (day/night) with ambient light conditions. *Nicotiana benthamiana* wild type and mutant plants for whitefly choice and no-choice assays were maintained at 26°C and a 16:8 h light:dark photoperiod in a growth room and, for seed production, in a growth chamber (Percival Scientific, Perry, IA) at 27/24°C (day/night) with a 16:8 h light:dark photoperiod.

### Identification of ASAT1 and ASAT2 orthologs in N. benthamiana

To identify ASAT1 and ASAT2 orthologs in *N. benthamiana*, protein sequences of *Salpiglossis sinuate* and *Solanum lycopersicum* ASAT1 and ASAT2 (Moghe et al., 2017) were compared to predicted proteins encoded by the *N. benthamiana* genome. Sequences with >67% identity were selected as potential ASAT1 and ASAT2 candidates and nucleotide sequences were obtained from the Solanaceae Genomics Network (www.solgenomics.net). The candidate ASAT sequences also were confirmed by comparing them to the most recent published *N. benthamiana* genome assembly (Schiavinato et al., 2019). To confirm the nucleotide sequences of *N. benthamiana ASAT1* and *ASAT2*, genes were amplified with ASAT1F/ASAT1R and ASAT2F/ASAT2R primers (Table S4) using genomic DNA as the template. Amplified fragments were cloned in pDONOR™207 (ThermoFisher Scientific, US) and were sequenced in their entirety using Sanger sequencing. This confirmatory sequencing showed no differences relative to the published *N. benthamiana* genome.

### Phylogenetic analysis of *N. benthamiana* ASATs

A protein phylogenetic tree of previously annotated Solanaceae ASATs (Figures 2, S1, S2; Tables S1, S2) was constructed using maximum likelihood method. Briefly, the ASATs protein sequences were aligned in program ClustalW (Thompson et al., 1994). Then the alignment was improved by removing the spurious sequences and poorly aligned regions (gap threshold at 0.25) using the program TrimAL v 1.4.rev22 (Capella-Gutierrez et al., 2009). Finally, an unrooted maximum likelihood tree was generated using the improved alignment with a bootstrap of 1000 in RAxML v8.2.12 (Stamatakis, 2014). The tree was visualized and presented using FigTree v1.4.4 (http://tree.bio.ed.ac.uk).

### sgRNA design and plasmid cloning

Single-guide RNAs (sgRNA) targeting *ASAT1* and *ASAT2* were designed based on the coding regions using online software, CRISPR-P v2.0 (Liu et al., 2017) and CRISPRdirect (https://crispr.dbcls.jp/), based on two parameters, cleavage efficiency and potential off-targets. Additionally, only sgRNAs with >40% GC content were selected. Three Cas9/gRNA constructs each were constructed for *ASAT1* and *ASAT2* following a previously developed CRISPR/Cas9 system (Jacobs et al., 2015). Four segments of DNA were prepared with 20 bp overlaps on their ends: ssDNA gRNA oligo, linearized p201N:Cas9 plasmid (Addgene 59175-59178), the *Medicago truncatula* (Mt) U6 promoter (377 bp), and a scaffold DNA (106 bp). For ssDNA gRNA, oligonucleotides targeting either the sense or antisense sequence of target genes were designed as: sense oligo TCAAGCGAACCAGTAGGCTT--GN19--GTTTTAGAGCTAGAAATAGC, and antisense oligo GCTATTTCTAGCTCTAAAAC--N19C—AAGCCTACTGGTTCGCTTGA. The gRNA oligonucleotide sequences are shown in Figure 3 and Table S4.

Oligonucleotide sequences were synthesized by Integrated DNA Technologies (www.idtdna.com). One μl of each 100 μM oligo was added to 500 μl 1x NEB buffer 2 (New England Biolabs, www.neb.com). The p201N:Cas9 plasmid was linearized by digestion with *Spe*1 (www.neb.com) in 1x buffer 4 at 37°C for 2 h, followed by column purification and a second digestion with *Swa*l in 1 x buffer 3.1 at 25°C for 2 h. Complete plasmid digestion was confirmed on a 0.8% agarose gel. The MtU6 promoter and Scaffold DNAs were PCR-amplified from the pUC gRNA Shuttle plasmid (Jacobs et al., 2015) using the primers *Swa*l_MtU6F/MtU6R and ScaffoldF/Spe_ScaffoldR, respectively (Table S4). The PCR reactions were performed with a high-fidelity polymerase (2x Kapa master mix; www.sigmaaldrich.com) using the program: 95°C for 3 min followed by 31 cycles of 98°C for 20 sec, 60°C for 30 sec, 72°C for 30 sec, and a final extension of 72°C for 5 min. Finally, cloning was done using the NEBuilder® HiFi DNA Assembly Cloning Kit. For each reaction, the four pieces of DNA were mixed in a 20 μl reaction with the NEBbuilder assembly mix with a final concentration of 0.011 pmol (∼100 ng) of p201N:Cas9 plasmid, 0.2 pmol of MtU6 amplicon (∼ 50 ng), scaffold amplicon (∼ 12 ng) and ssDNA gRNA oligo (60-mer, 1 μl). The reactions were placed in thermal cycler at 50°C for 1 h.

Two μl of the cloning reaction were transformed into 50 μl of the One Shot™ Top10 chemically competent cells (Invitrogen, www.thermofisher.com) and plated on LB (Bertani, 1951) agar medium with 50 μl/ml kanamycin for selection of transformants. Colonies with the correct inserts were screened using the Ubi3p218R and IScelR primers (Table S4). Plasmids carrying the designed gRNA constructs were then transformed into *Agrobacterium tumefaciens* strain GV3101 for generating transgenic plants. All constructs were confirmed by Sanger sequencing.

To avoid off-target effects, gRNAs were further checked by comparison against the reference *N. benthamiana* genome v1.0.1 (www.solgenomics.net). Only two sites in the *N. benthamiana* genome were found to have non-target matches >17 nt (both with 1 internal mismatch), and with the NGG PAM sequence on the correct strand. These two sites were checked by PCR amplification and Sanger sequencing and showed no unexpected editing in our *asat1-1*, *asat1-2*, *asat2-1*, or *asat2-2* mutant plants. Primers used for off-target Sanger sequencing are listed in Table S4.

### Confirmation of gRNA functionality by transient expression

For both *ASAT1* and *ASAT2*, we *Agrobacterium-*infiltrated *N. benthamiana* plants at the four-leaf stage. For *ASAT1*, we used *Agrobacterium* carrying the Cas9/gRNA constructs at OD of 1 and 3, whereas for *ASAT2*, we used an OD 1.5. Each leaf was saturated with *Agrobacterium* solution. After infiltration, the plants were cultured in a growth chamber for 2 days and then the infiltrated whole leaves were collected for genomic DNA extraction and tested by PCR amplification for detection of insertion/deletion polymorphisms in the target region (Figure S9). During the method optimization, a positive control construct targeting the *N. benthamiana Drm3* gene (gRNA: GCCACTATCTGGCCGGGGAC, provided by the Greg Martin lab, Boyce Thompson Institute) was infiltrated in parallel.

### Stable mutagenesis of ASATs using tissue culture

Stable *ASAT* mutant *N. benthamiana* plants were created in the Boyce Thompson Institute plant transformation facility using CRISPR/Cas9 with gRNAs that had been confirmed to be functional as described above and a previously described protocol (Van Eck et al., 2019), with minor modifications. To prepare plants for transformation, we disinfected *N. benthamiana* seeds with 1.5 ml 1.25% sodium hypochlorite (1:5 dilution of 5.25% sodium hypochlorite Clorox bleach), with 100 μl Tween-20 added, for 20 min on a shaker platform. After sodium hypochlorite treatment, seeds were rinsed three times with sterile water. Subsequently, we placed seeds on Tobacco Seed Germination Medium (composition per liter: 1.08 g Murashige and Skoog salts (Murashige and Skoog, 1962), 30 g sucrose, and 8 g agar, pH = 5.7±0.1) and incubated the seeds under yellow filtered light with a 16-h photoperiod at 27°C. After ∼2 weeks, the seedlings were transferred to the Rooting Medium (composition per liter: 2.15 g Murashige and Skoog salts, 30 g sucrose, 1 ml B5 vitamin/amino acid stock, and 8 g agar, pH = 5.7±0.1). After ∼6 weeks, the plants were ready for *Agrobacterium* infection. First, *Agrobacterium* carrying the gRNA plasmids was amplified in YEP medium (composition per liter: 10 g yeast extract, 10 g peptone, and 5 g NaCl with antibiotics, 50 µg/µl gentamycin, 50 µg/µl kanamycin, and 50 µg/µl rifampicin), and *Agrobacterium* culture then was pelleted and combined to a certain OD (OD 1.0 for *ASAT1* gRNAs, and OD 1.0 or 1.5 for *ASAT2* gRNAs) with filtered cell suspension buffer (10 mM MES, 10 mM MgCl_2_ and 200 µM acetosyringone). Then, fully expanded but immature *N. benthamiana* leaves were dissected into 5 mm segments by removing the leaf margins. The leaf segments were incubated in the *Agrobacterium* culture for 30 min with shaking. The incubated leaf segments were transferred to co-cultivation medium plates (composition per liter: 4.3 g Murashige and Skoog salts, 30 g sucrose, 1 ml B5 vitamin/amino acid stock, and 5 g agar, pH = 5.7±0.1) and kept in the dark at room temperature. Leaf segments treated with YEP medium were used as a negative control. After three days, the leaf segments were transferred from co-cultivation medium to shoot bud initiation medium with selections (composition per liter: 4.3 g Murashige and Skoog salts, 30 g sucrose, 1 ml B5 vitamin/amino acid stock, 0.1 mg NAA, 1.0 mg BA, and 6 g agar, pH=5.7±0.1; with 200 μg/ml kanamycin and 250 μg/ml Timentin). Medium with Timentin but without kanamycin was used as a positive control. Leaf segments on selection medium were incubated under a 16:8 h light-dark photoperiod at 24-25°C under yellow filtered light. The leaf segments were sub-cultured every 2-weeks onto the same medium until shoot buds were visible. Finally, to generate roots, shoot buds were transferred to rooting medium (composition per liter: 2.15 g Murashige and Skoog salts, 30 g sucrose, 1 ml B5 vitamin/amino acid stock, 8 g agar with 200 μg/ml kanamycin and 250 μg/ml Timentin, pH=5.7±0.1).

### Confirmation of homozygous mutant plants in the T2 generation

Rooted *N. benthamiana* plants from tissue culture were transferred to soil (T0 generation). CRISPR/Cas9-induced mutations were identified by PCR amplification of genomic regions of the gRNA target sites in *ASAT1* and *ASAT2* (Figure 3), followed by Sanger sequencing. Lines with mutations were used to generate T1 plants, which were subjected to PCR amplification and sequencing to confirm homozygous mutations. T2 seeds from confirmed homozygous mutant *asat1-1*, *asat1-2*, *asat2-1*, and *asat2-2* T1 plants were used for all experiments. Homozygous mutations were confirmed in randomly selected T2 plants by PCR amplification and Sanger sequencing. The presence of Cas9 in transgenic plants in the T0, T1, and T2 generations was confirmed by PCR amplification and agarose gel electrophoresis. Primers that were used to identify mutations and confirm the presence of Cas9 are listed in Table S4.

### Acylsugar measurements by LC/MS

Liquid chromatography/mass spectrometry (LC/MS) was used to confirm the effect of the *ASAT* mutations by measuring acylsugar content in leaf extracts from wildtype and *ASAT* mutant plants. New leaflets were rinsed in acylsugar extraction solution (3:3:2 acetonitrile:isopropanol:water, 0.1% formic acid, and 1 μM Telmisartan as internal standard) and gently agitated for 2 min. Then, the extraction solutions were transferred to LC/MS glass vials, and the leaves were air dried for leaf weight measurements.

Chromatography of leaf surface washes was performed on a ThermoScientific Ultimate 3000 HPLC with a glass vial autosampler and coupled with a Thermo Scientific Q Exactive™ Hybrid Quadrupole-Orbitrap™ Mass Spectrometer (Mass Spectrometry Facility at Boyce Thompson Institute). Acylsugar extracts were separated on an Ascentis Express C18 HPLC column (10 cm × 2.1 mm × 2.7 μm) (SigmauAldrich, St. Louis, MO) with a flow rate of 0.3 ml/min, using a gradient flow of 0.1% formic acid (Solvent A) and 100% acetonitrile (Solvent B). We used a 7-min LC method for metabolite profiling, which involved a linear gradient from 95:5 A:B to 0:98 A:B. Fulluscan mass spectra were collected (mass range: m/z 50–1000) in both positive and negative electron spray ionization (ESI) modes. Mass spectral parameters were set as follows: capillary spray voltage 2.00 kV for negative ion-mode and 3.00 kV for positive ion-mode, source temperature: 100°C, desolvation temperature 350°C, desolvation nitrogen gas flow rate: 600 liters/h, cone voltage 35 V. Acylsugars were identified and annotated using Thermo Xcalibur Qual Browser (Thermo Fisher) and MS-DIAL v4.20 based on the MS/MS peak features and neutral losses. The acylsugar abundances were estimated using peak areas at the respective *m/z* channel under negative ESI mode. Acylsugar quantification was first normalized to the internal control Telmisartan to account for technical variation between samples, and then normalized to the leaf dry weight to allow comparisons between samples.

### Insect choice and no-choice bioassays

To measure *M. persicae* and *M. euphorbiae* growth, we caged aphids on individual leaves of mutant and wildtype 4 to 5-week-old *N. benthamiana* (Figure S10A, B). Twenty adult *M. persicae* from *N. tabacum* (naïve to *N. benthamiana*) were placed in each cage and allowed to generate nymphs for ∼12 hrs. Twenty-five nymphs were left in each cage and were monitored for 5 d to assess nymph survival. At the end of the *M. persicae* survival monitoring period, five *M. persicae* were left in each cage and reproduction was monitored for one week. Finally, the remaining *M. persicae* were collected to measure aphid size by taking a picture and assessing the area of each aphid using ImageJ (Schneider et al., 2012). Ten adult *M. euphorbiae* from a colony on tomato cv. Moneymaker were placed in each individual cage on *N. benthamiana* leaves. Surviving aphids and progeny were counted after 24 hours.

*Myzus persicae* and *M. euphorbiae* choice assays were performed with detached leaves from 4 to 5-week-old *N. benthamiana*. Two similarly-sized leaves from individual *ASAT* mutant and wildtype plants were cut and placed in 15-cm Petri dishes, with their petioles inserted in moistened cotton swabs (pairwise comparison is shown in Figure S10C). Ten naïve adult aphids, which had not previously encountered *N. benthamiana*, were released at the midpoint between pairs of leaves (wildtype, *asat1*, or *asat2*), and the Petri dishes were placed under 16:8 h light:dark photoperiod. The aphids on each leaf were counted at 24 h after their release in the Petri dishes.

To measure whitefly survival and fecundity on wildtype and *asat2-1 N. benthamiana* plants, cages were set up with *N. benthamiana* plants at the seven-leaf stage (approximately 3 weeks old). Each cage contained three plants, either wildtype or *asat2-1*. Ninety adult whiteflies reared on *Brassica oleracea* (variety Earliana; www.burpee.com, catalog number 62729A) were introduced into each cage (60 x 60 x 60 cm) with *N. benthamiana* (30 whiteflies/plant) and were allowed to feed for three days at 26°C with a 16:8 h light:dark photoperiod. The numbers of whiteflies surviving on each host plant were counted, after which the remaining insects were killed with insecticidal soap. The following day, the number of whitefly eggs on each plant was counted. This experiment was conducted twice with similar results.

For whitefly choice assays, wildtype and *asat2-1* plants at the seven-leaf stage were placed together in the same cage. Approximately 150 whiteflies from cabbage plants were moved into each cage. After 24 h at 26°C with a 16:8 h light:dark photoperiod, live and dead whiteflies were counted on the plants and elsewhere in the cage. This experiment was repeated three times.

Eggs of *H. zea, H. virescens* and *T. ni* were hatched on artificial diet (Southland Products, Lake Village, Arkansas). Neonate larvae were confined onto individual *N. benthamiana* leaves, one larva per plant, using 10 x 15 cm organza mesh bags (www.amazon.com, item B073J4RS9C). After ten days, the surviving larvae were counted and weighed. In a separate experiment, *H. zea, H. virescens*, and *T. ni* were reared on artificial diet (beet armyworm diet, www.southlandproducts.net) for five days. Individual five-day-old caterpillars were weighed and then confined on 4 to 4.5-week-old *N benthamiana* plants using 30 cm x 60 cm micro-perforated bread bags (www.amazon.com). After seven days, the surviving larvae were weighed again. Relative growth rate was calculated as: ln(((day-12 mass)/(mean day-5 mass))/7).

### Leaf water loss and temperature assays

To measure the leaf water loss, two leaves from each of eight plants were detached. The fresh weight of each leaf was determined on a Sartorius Ultra Micro Balance. All leaves were placed at 23°C and a 16:8 h light:dark photoperiod. Each leaf was weighted again after 24 h and the percentage of water loss was calculated as [(fresh weight - final weight)/fresh weight]*100%.

Thermal images were acquired in the growth chamber environment using a thermal camera (A655sc, FLIR Systems Inc., Boston, MA, USA) with a spectral range of 7.5–14.0 mm and a resolution of 640 x 480 pixels. The camera was placed approximately 1 m away from each plant and a white background was used when the plant images were acquired. One region of interest, corresponding to the perimeter of each leaf, was specified per leaf for each of 20 leaves per genotype. Using the FLIR ResearchIR Max software (version 4.40.9.30) thermal images files were exported as CSV files. Images were segmented from the background using Gaussian mixture models in MATLAB to determine the temperature of each leaf. After segmentation, the temperature was averaged across the segmented leaf.

Hyperspectral images were acquired in a dark room using a hyperspectral imager (SOC710, Series 70-V, Surface Optics Corporation, San Diego, CA, USA) that covered a spectral range from 400 to 1000 nm for 128 wavebands. Image acquisition and recording were performed using a Dell DELL XPS 15 9570 laptop computer that controls the camera. The camera was fixed using a stand with the lens facing the plants and capturing top view images approximately 1 m away. A Spectralon tile (Labsphere, Inc, North Sutton NH, USA) was placed next to the plant trays, covering one corner of the image to facilitate subsequent image processing and calibration. The nominal reflectance value for the Spectralon tile was 99% and it had a 30.5 cm x 30.5 cm reflective area. Lighting consisted of two halogen lamps placed at ∼ 45° angles on either side of the camera to create an even light distribution. All image analysis was performed in HSIviewer, a MATLAB package (Stone et al., 2020). White reflectance calibration was performed using the Spectralon tile. One region of interest ROI was specified for each of 20 leaves per *N. benthamiana* genotype. This ROI corresponded to the perimeter of each leaf. From each hyperspectral cube image, the vegetation pixels (green portion of the plant) were extracted using the Normalized Difference Vegetation Index (NDVI). Mean reflectance (R) was calculated per band per 10 leaves in order to obtained the WBI results

To calculate NDVI and WBI we used the following formulas, where *R* corresponds to the reflectance at a specific wavelength (nm): WBI = (R970/R900) (Penuelas et al., 1993) and NVDI = (R750 - R705)/(R750 + R705) (Gitelson and Merzlyak, 1994).

### Virus transmission assays

For virus transmission assays, young adult *M. persicae* were starved for 3 h and then allowed to feed for 30 min on 6-week-old wildtype or *asat2-1 N. benthamiana* plants infected with TuMV-GFP (Lellis et al., 2002; Casteel et al., 2015). Aphids were transferred in groups of ten to individual 4-week-old wildtype or *asat2-1* recipient plants. The aphids were allowed to feed on recipient plants for 24 h and were then removed. Donor and recipient plants were kept in the Conviron growth chamber at 23°C and a 16:8 h light:dark photoperiod. The development of GFP fluorescence was monitored every 24 h using a 395 nm UV flashlight (www.walmart.com) in the dark, until no new GFP signals developed from one day to the next. In each experiment, the virus donor plant group consisted of eight TuMV-GFP infected plants (2 weeks after TuMV-GFP infection by rub-inoculation) of each genotype, with aphids randomly spread on the plants. Each recipient group consisted of 20 plants of each genotype, with the aphids confined on individual plants. The experiment was repeated three times with similar results, and the results were combined for statistical analysis of differences between groups using one-way ANOVA with a block effect followed by a Tukey HSD post hoc test.

### Plant-mediated virus induced gene silencing in whitefly with WT and *asat2-1*

For each target gene, a 200-550 bp RNA sequence was designed using the E-RNA*i* webserver (https://www.dkfz.de/signaling/e-rnai3/). The designed fragments were checked against the *B. tabaci* MEAM1 (Chen et al., 2016) and *N. benthamiana* (Bombarely et al., 2012) reference genomes using local blast (Mount, 2007) to avoid potential off-target sequences with full matches ≥ 19 nt. The identified gene fragments were then PCR-amplified from whitefly cDNA and cloned into a binary tobacco rattle virus VIGS vector pTRV2 (Liu et al., 2002) using the Invitrogen Gateway recombination cloning technology (Invitrogen, USA). As a positive control, we used a pTRV2-PDS construct with an RNA fragment targeting *N. benthamiana* phytoene desaturase (Velásquez et al., 2009). As negative controls, we used pTRV2-EV (empty vector) and pTRV2-GFP, with an RNA fragment targeting *Aequorea victoria GFP*, a sequence that is not present in either *N. benthamiana* or *B. tabaci*. The PCR primers used to generate the VIGS vectors are in Supplemental Table S4.

To infiltrate plants, plasmids (pTRV1 and pTRV2 containing constructs targeting the genes of interest) were transformed into *Agrobacterium tumefaciens* GV3101. Equal amounts of *Agrobacterium* with TRV1 and TRV2 were resuspended in infiltration medium (10 mM MES, 10 mM MgCl_2_, 200 μM acetosyringone) to an OD 0.3 and incubated at room temperature for 3 h. The *Agrobacterium* mixture was infiltrated using a 1-ml needleless syringe to saturate three leaves of ∼4-week-old *N. benthamiana*. Inoculated plants were kept in a Conviron growth chamber for 2∼3 weeks until control plants infiltrated with pTRV2-PDS showed photobleaching symptoms. Successful growth of TRV targeting the genes of interest in whiteflies was confirmed by PCR and the confirmed plants were then used for insect bioassays.

For whitefly bioassays, three infiltrated plants were used for each gene of interest, and three cages were attached to each plant. Ten young adult whiteflies (emerged within the past week) were placed in each cage. After seven days, the survival rates of whiteflies were assessed and compared across different treatments. The surviving whiteflies were collected for qPCR analyses of target gene expression using quantitative PCR using the PowerUp^TM^ SYBR^TM^ green Master Mix (AppliedBiosystem, Thermo Fisher Scientific) on a QuantStudio^TM^ 6 Flex Real-time PCR system (AppliedBiosystem, Thermo Fisher Scientific), and results were analyzed using the 2^-ΔΔCT^ method (Schmittgen and Livak, 2008). The whitefly VIGS bioassays, comparing *asat2-1* and wildtype *N. benthamiana*, were repeated three times with similar results. Two replicates of a separate experiment was set up on *asat2-1* plants to collect whiteflies after one day for gene expression analyses.

### Statistical analysis

All statistical comparisons were conducted using SPSS v25, R and MATLAB R2019a (MathWorks, Inc., Natick, MA, USA). ANOVA followed by a Dunnett’s post hoc test was used to determine differences in leaf water loss, leaf temperature, and WBI across genotypes in each data set. ANOVA followed by a Duncan post hoc test was used for aphid bioassay and LC/MS results. A Chi-square test was used to test for differences in pairwise aphid choice assays, whitefly, and lepidopteran assays.

## Acknowledgements

We want to thank Patricia Keen and Joyce Van Eck for their help with *N. benthamiana* stable transformation and access to the tissue culture facility, Danielle Preston, Angela Douglas, and Jane Polston for providing whitefly colonies, Isgouhi Kaloshian for the potato aphid colony, Ning Zhang and Greg Martin for sharing the p201N-Cas9 construct and the *Drm3* sgRNA control construct, and William Stone and Thomas Lawton for providing custom image processing software. This research was supported by Cornell startup funds to G.D.M., Deutsche Forschungsgemeinschaft award #411255989 to L.H.K., and United States Department of Agriculture Biotechnology Risk Assessment Grant 2017-33522-27006, US National Science Foundation award IOS-1645256, and Defense Advanced Research Projects Agency (DARPA) agreement HR0011-17-2-0053 to G.J, and US National Science Foundation award #1723926 to C.L.C. G.S. is part of a team supporting DARPA’s Insect Allies program under agreement HR0011-17-2-0055. M.A.G. is part of a team supporting DARPA’s Advanced Plant Technologies program under agreement HR0011-18-C-0146. The views and conclusions contained in this document are those of the authors and should not be interpreted as representing the official policies of the U.S. Government.

## Author Contributions

G. J. and H.F. conceived the original research plans; H.F., S.S., L.A., H.X., L.K., J.D.T, S.H.C., and A.N.F. performed the experiments; H.F., L.A., L.K., and G.D.M. analyzed the data; C.L.C., M.A.G., G.D.M., G.S., and G.J. supervised the experiments; H.F. and G.J. wrote the article with contributions from all of the authors; G.J. agrees to serve as the contact author responsible for communication and distribution of samples.

---

**Figure S1.**
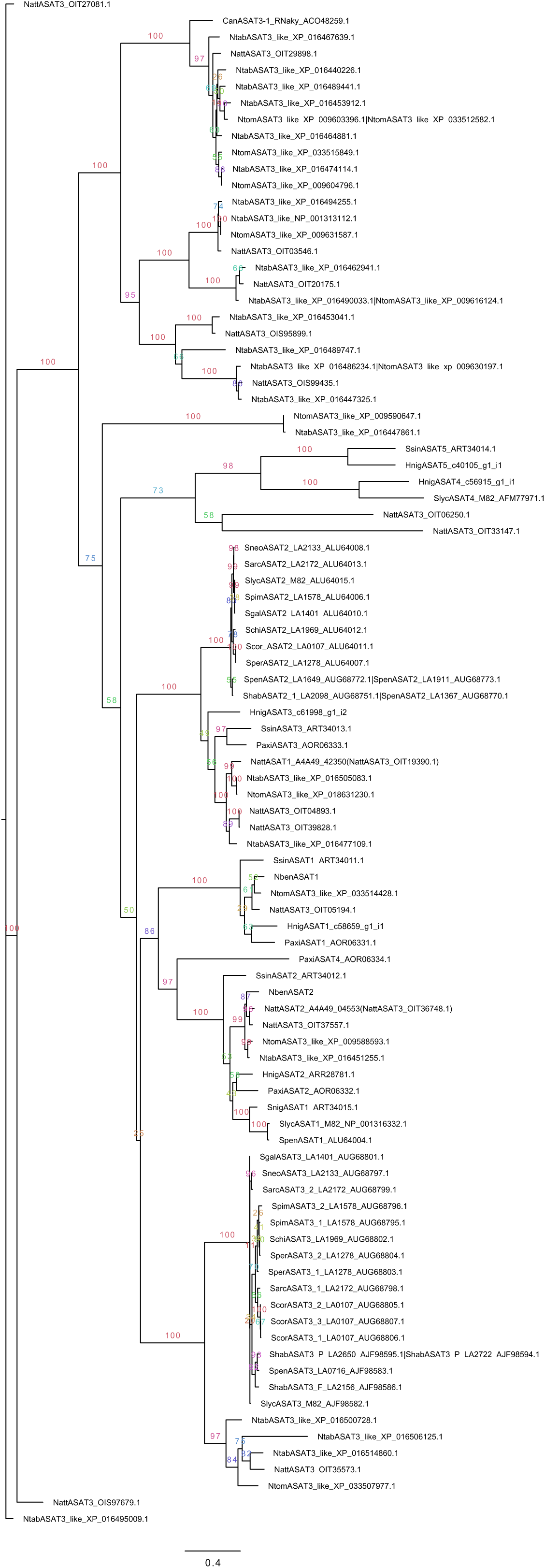
**Phylogenetic analyses of annotated ASATs from Solanaceae species** The ASAT proteins used in constructing the tree are listed in Tables S1 and S2. The evolutionary history of ASATs from Solanaceae species was inferred by using the Maximum likelihood method in RAxML. The branch labels indicate the percentage of trees in which the associated taxa clustered together (bootstrap of 1000).

**Figure S2.**
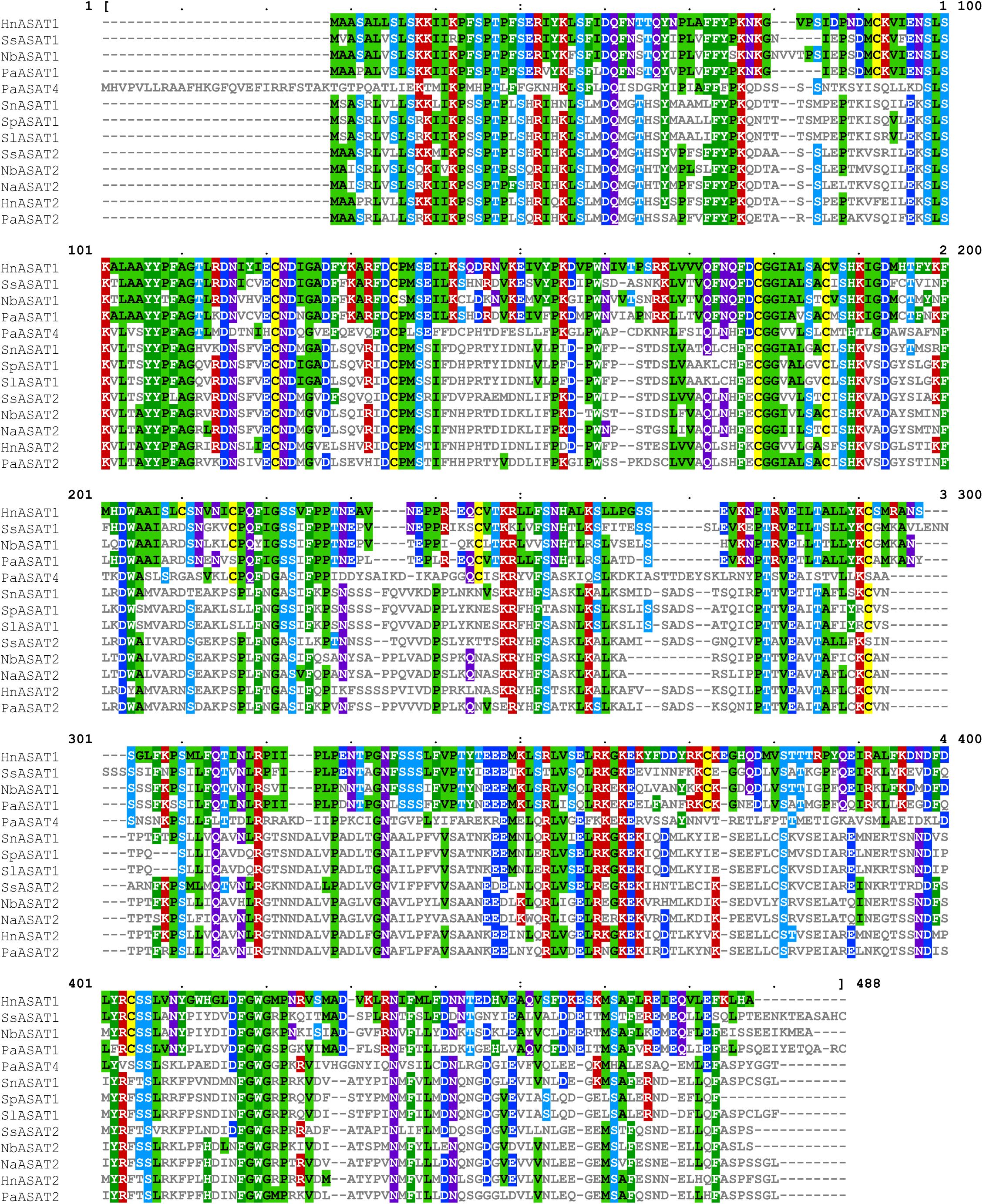
**Protein sequence alignment of *Nb*ASAT1 and *Nb*ASAT2 with other functionally characterized ASATs in the Solanaceae species**. The alignment was generated using Clustal Omega and visualized using Mview. Colors are coded based on the level of sequence identity.

**Figure S3.**
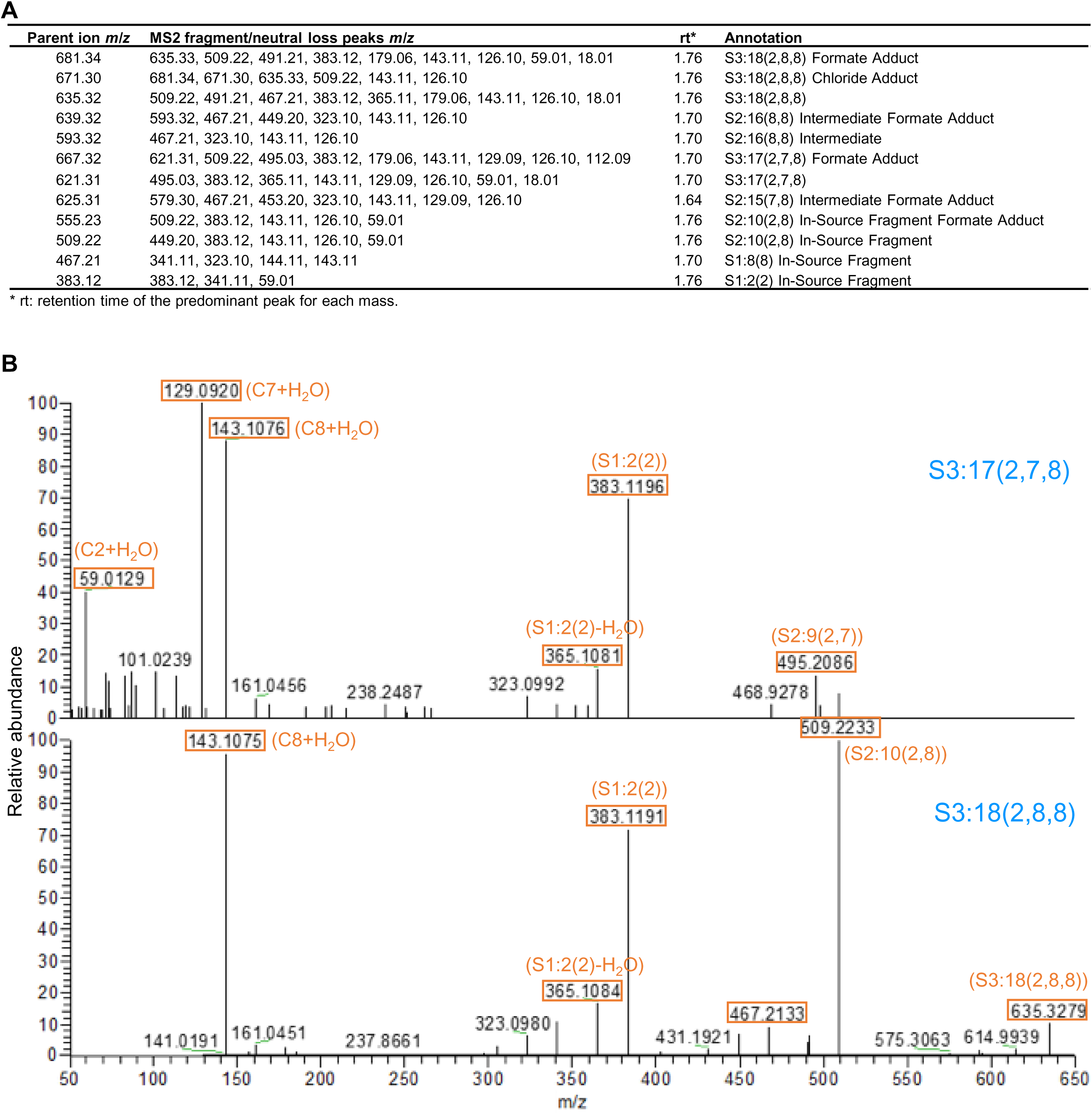
**MS/MS chromatography of *N. benthamiana* acylsugars (A)** *N. benthamiana* **a**cylsugar LC/MS peak identification and annotation **(B)** MS^2^ chromatography of the two predominant acyl sugars, S2:17(2,7,8) and S2:18(2,8,8). The names of the acylsugars are highlighted in blue and the chromatography is aligned on the x axis based on *m/z*. Highlighted in orange boxes are some of the abundant characteristic mass features, with their compositions in parenthesis.

**Figure S4.**
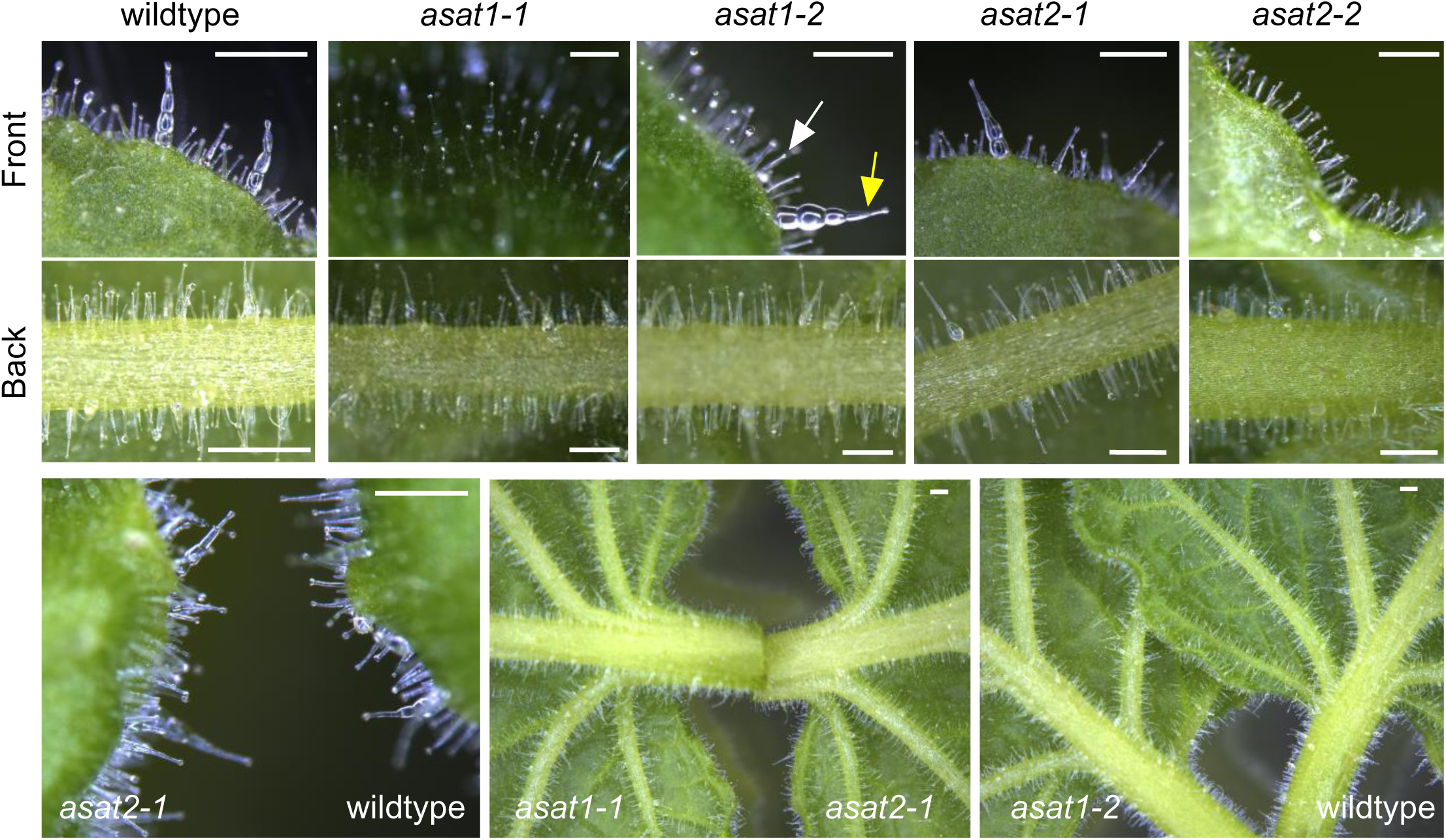
***ASAT* mutations do not affect trichome morphology and abundance** The white arrow indicates the small trichomes and yellow arrow indicates the large swollen-stalk trichomes. Trichome pictures were taken from both the front and back sides of the leaf. Examples of leaves are put side-by-side from different genotypes to facilitate comparisons. Scale bars = 200 μm.

**Figure S5.**
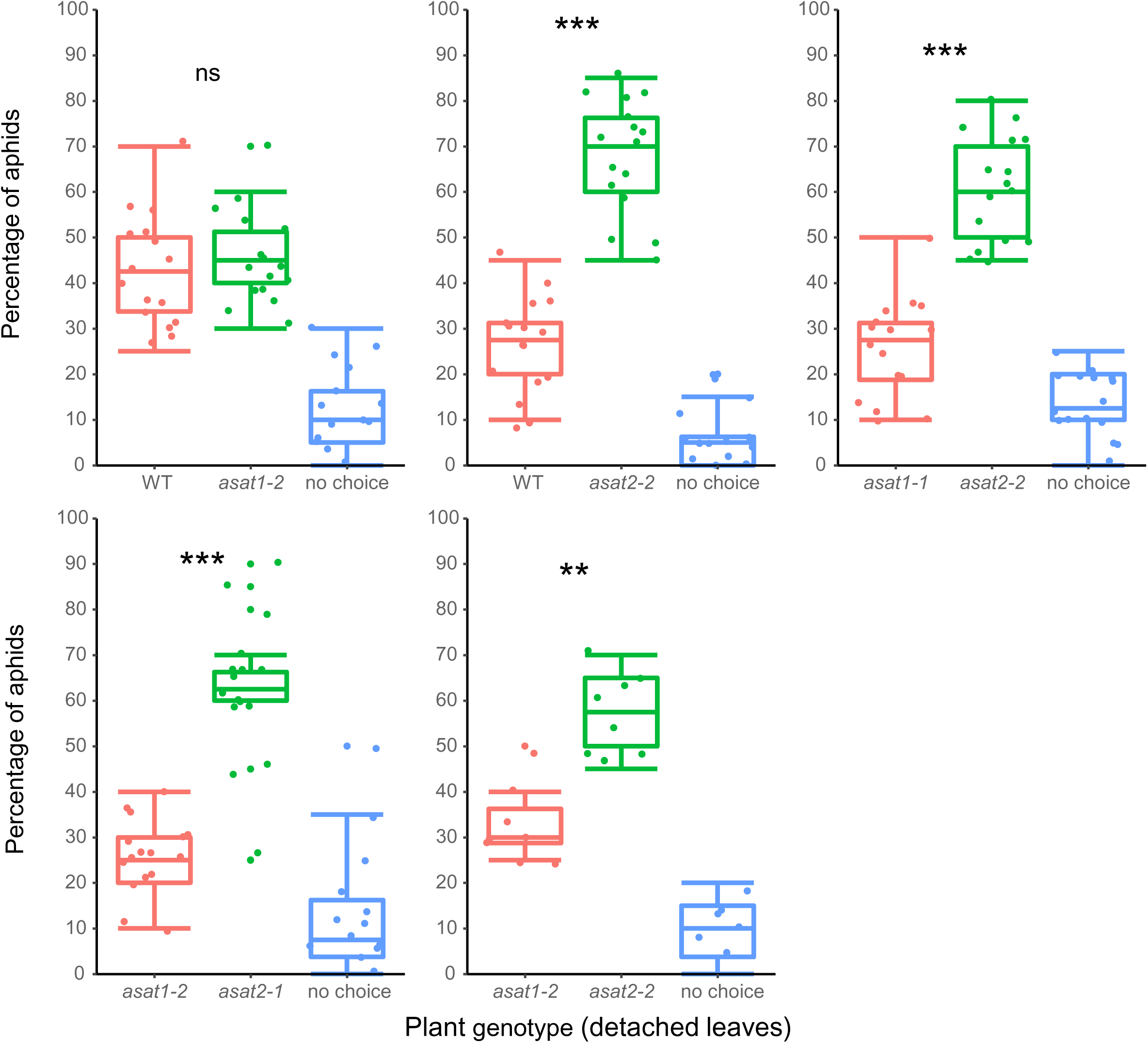
**Pairwise aphid choice assay with wildtype and *ASAT* mutant leaves** Each experiment included detached leaves from 5-8 plants of each genotype and was repeated twice with similar results. Chi-square test was used for difference for all choice assays between plant genotypes. \*\**p* < 0.01, \*\*\**p* < 0.001, ns: not significant. no choice: aphids were elsewhere in the Petri dish and not on leaves at the end of the experiment. The box plots show the median, interquartile range, maximum and minimum after removal of outliers, and the individual data points.

**Figure S6.**
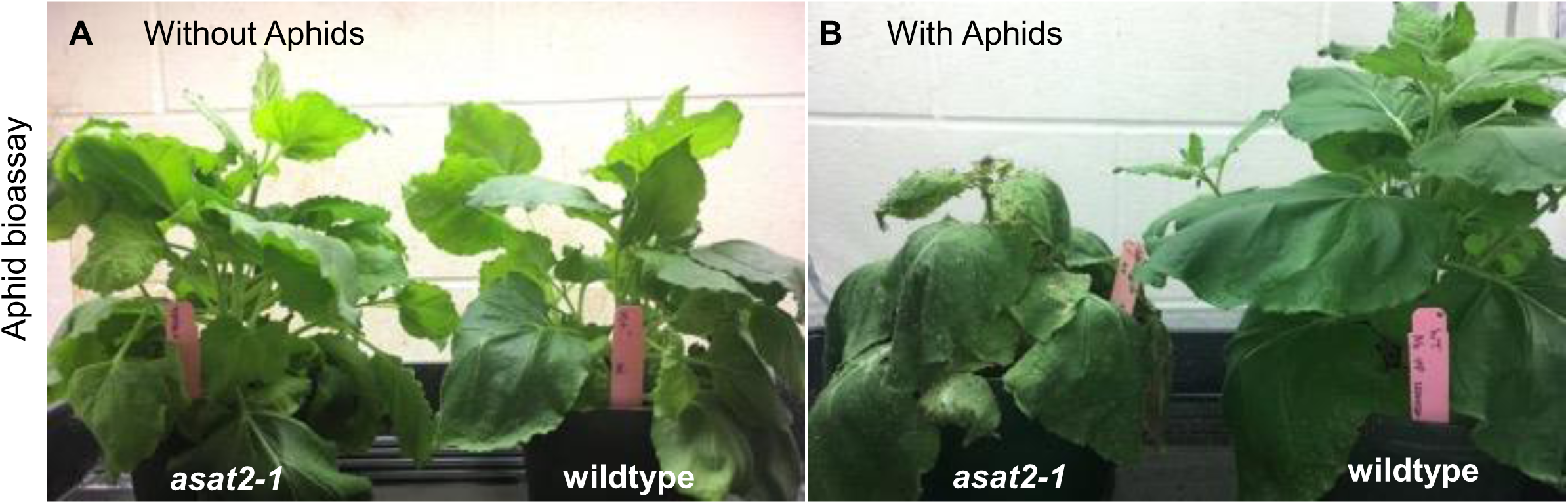
**Aphid bioassays on wildtype and *asat2-1* mutant *N. benthamiana*** (**A**) without *M. persicae* feeding. (**B**) with *M. persicae* feeding for one month. One wildtype (right) and one *asat2-1* (left) plant were placed in the same tray in an insect cage. For the aphid challenge, ten adult *M. persicae* (from an aphid colony on *N. tabacum*) were released on each of the wildtype and the *asat2-1* plants. The pictures were taken about one month after the aphid release. In the case of control plants without aphids (**A**), wildtype and *asat2-1* look similar. (**B**) After one month of aphid feeding, the asat2-1 and wildtype plants look very different, with the wildtype plant looking vigorous and the *asat2-1* line being smaller and wilted. On wildtype *N. benthamiana*, *M. persicae* reproduce slowly and sometimes not at all. If they do survive, they tend to feed on the undersides of senescing leaves. By contrast, *M. persicae* are able to establish a dense colony on the *asat2-1* mutant and feed from the youngest leaves, which tend to be the most nutritious but also the best-defended. Plants in three replications of this experiment looked similar.

**Figure S7.**
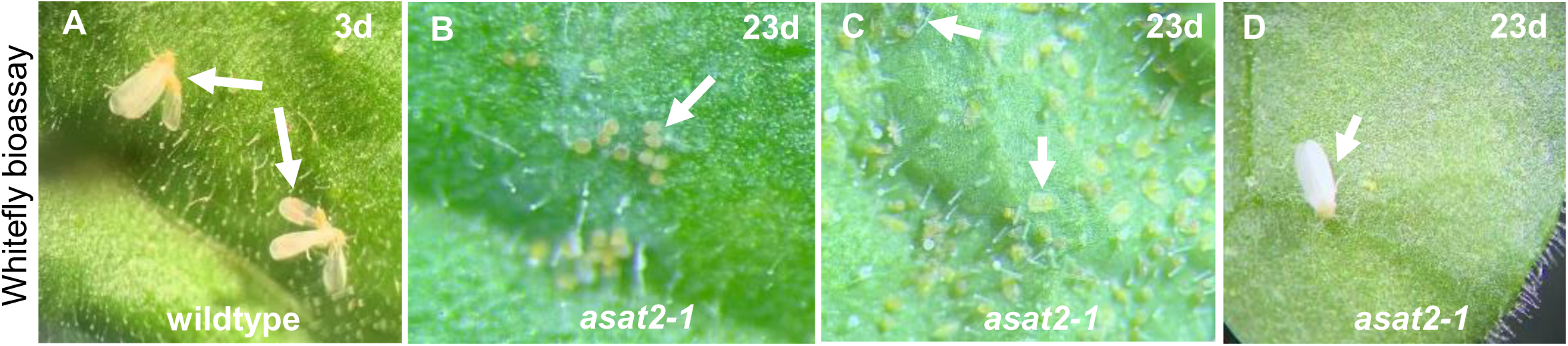
**Whitefly bioassays on wildtype and *asat2-1* mutant *N. benthamiana*** (**A**) Most whiteflies died within three days after release on wildtype *N. benthamiana*. Arrow indicates dead whiteflies. (**B-D**) Whiteflies survived and completed their life cycle on *asat2-1* mutant *N. benthamiana*. At 23 days after release, whiteflies of different life stages were present on *asat2-1* mutant plants. Arrows indicate eggs (**B**), eggshells and nymphs (**C**), and adults (**D**) on *asat2-1* mutant plants.

**Figure S8.**
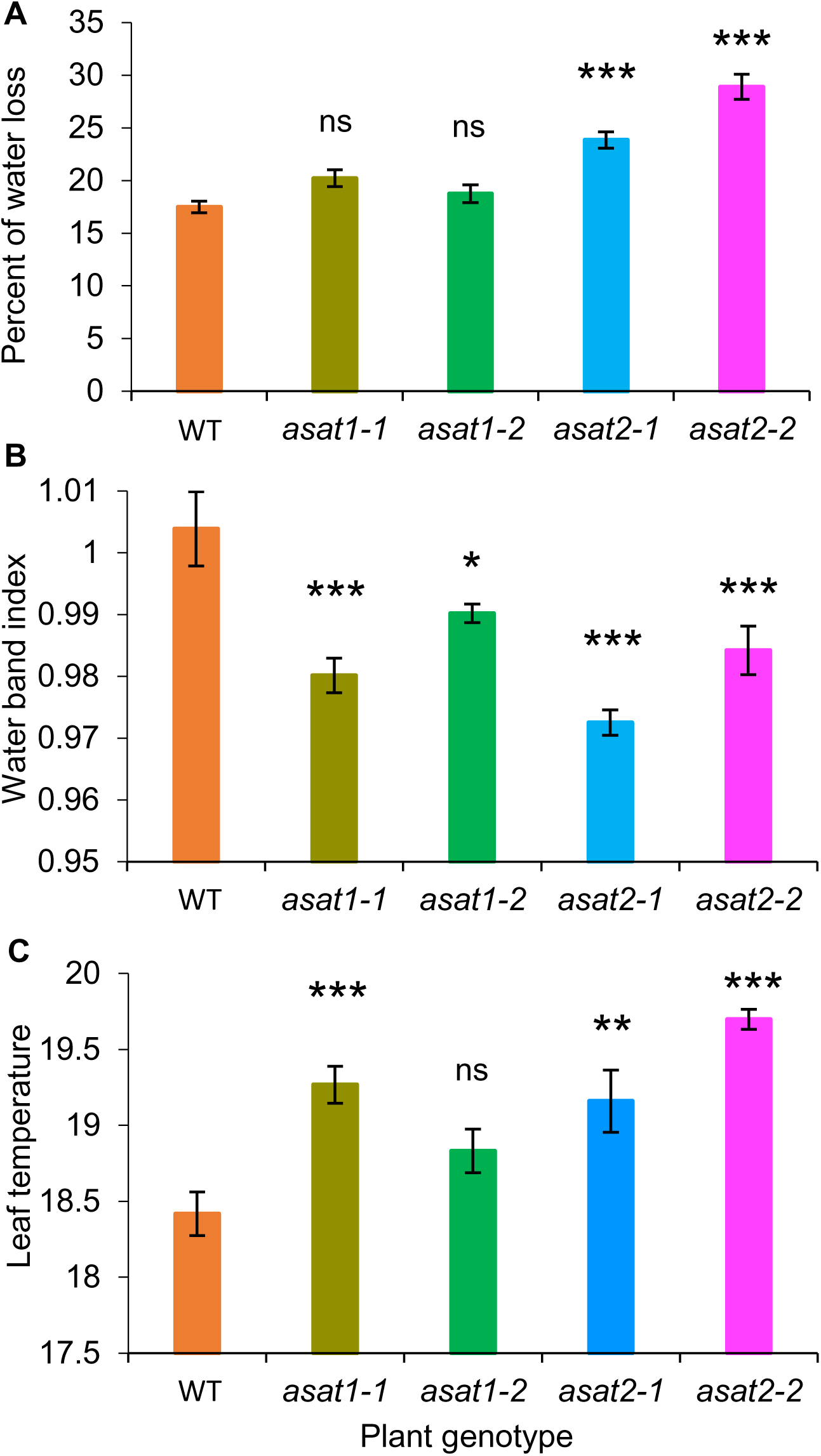
**Independent repeat of water loss and leaf temperature shown in Figure 8. (A)** Percent of water loss from detached leaves after 24 hours, mean +/- s.e. of n =16. (**B**) Leaf water content measured by the water band index from hyperspectral imaging, mean +/- s.e. of n = 10. (**C**) Leaf temperatures from leaves of different plant genotypes, mean +/- s.e. of n=10 for wildtype and n=5 for mutants. ns, not significant, *p < 0.05, **p < 0.01, ***p < 0.001, Dunnett’s test relative to wildtype control. Error bars = standard error. NS = not significant.

**Figure S9.**
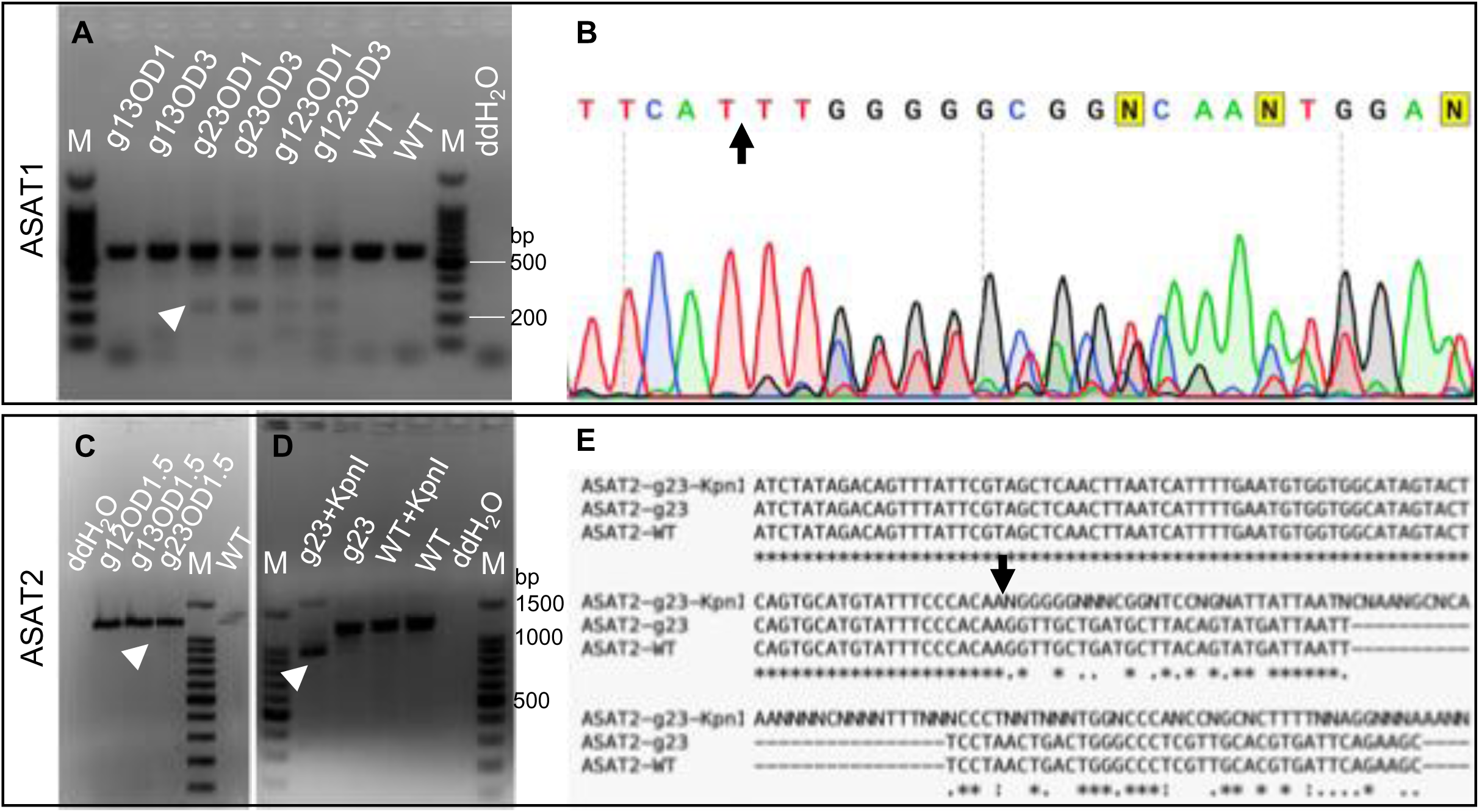
***ASAT* mutagenesis by transiently expressing Cas9 and gRNAs in *N. benthamiana* leaves via *Agrobacterium* infiltration** Three gRNAs each were used for *ASAT1* and *ASAT2* mutagenesis to test efficacy. (**A**) PCR of plants that were infiltrated with combinations of *ASAT1* gRNA constructs. (**B**) DNA sequence confirmation of a representative sample (*i.e.* g23OD3) that displayed a shorter band in panel A. (**C**) PCR of plants that were infiltrated with combinations of *ASAT2* gRNA constructs. (**D**) Restriction enzyme digestion site loss in PCR of g23OD1.5 (from panel C), where, prior to the PCR reaction, genomic DNA was digested with *Kpn*I to remove wild type *ASAT2*. The *Kpn*I restriction site is only present in the expected deletion region. (**E**) Sequence confirmation of sample g23+*Kpn*I from digestion site loss PCR in panel D. Arrow heads indicate the short versions of target genes after a deletion by the two working gRNA constructs. Arrows indicate the positions expected to be cut in gene sequences. The label for each lane in the gel: M, 100 bp DNA marker ladder; WT, samples from wild type plants; ddH2O, no genomic DNA PCR control.

**Figure S10.**
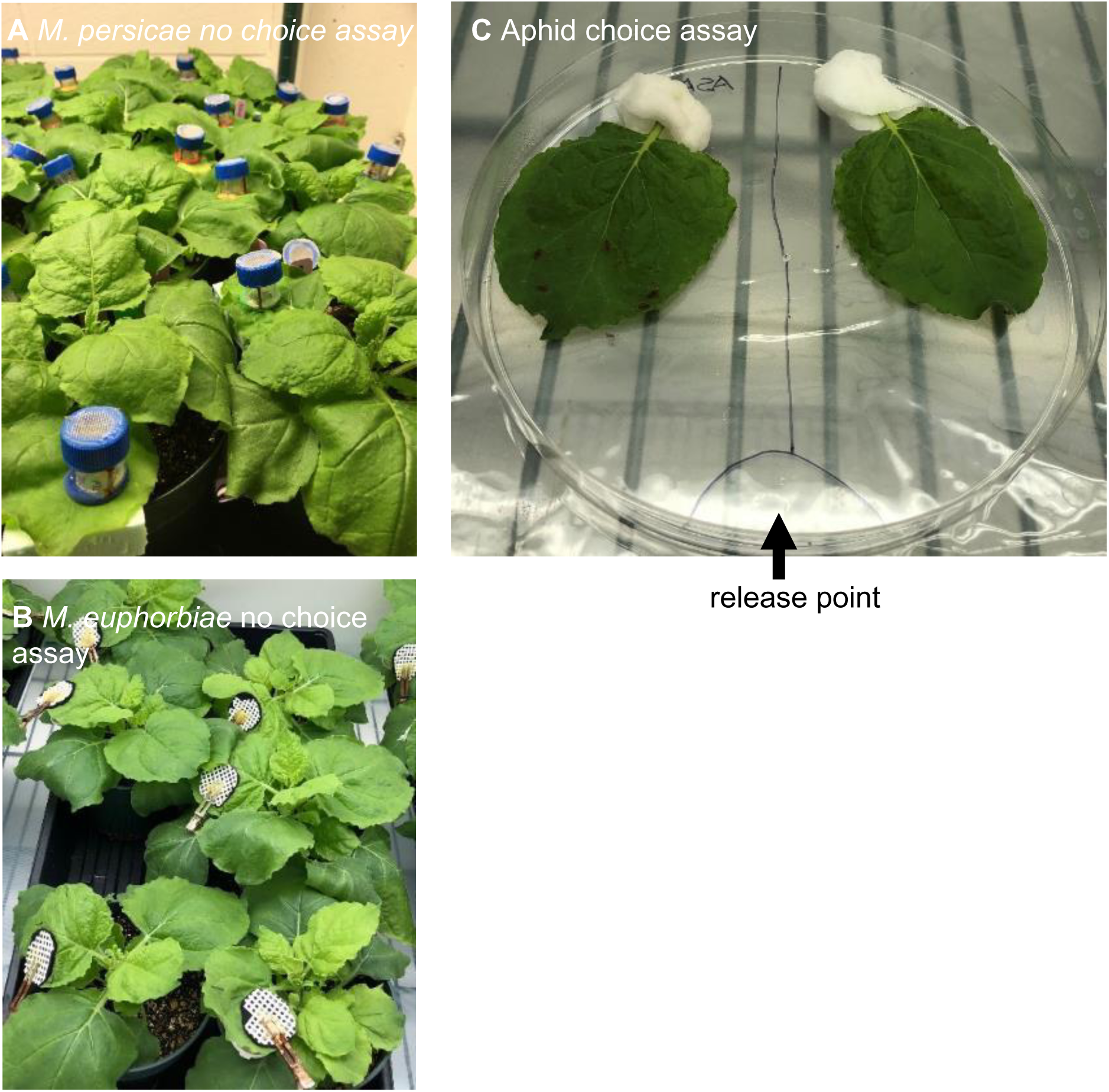
**Experimental setups for *Myzus persicae* bioassays** (**A**) *Myzus persicae* and (**B**) *Macrosiphum euphorbiae* no-choice assays. Aphids were caged on individual *Nicotiana benthamiana* leaves. Survival, reproduction, and size were assessed at time points described in the methods section. (**C**) Aphid choice assays. Two *N. benthamiana* leaves were placed in a 15 cm diameter Petri dish with their petioles inserted in moistened cotton swabs, ten aphids were released into the Petri dish at the indicated position, the dish was covered and placed under a 16:8 light:dark photoperiod, and, after 24 hours, the aphids residing on each leaf were counted.

