## Supplemental Table S2 for "An acylsugar-deficient *Nicotiana benthamiana* strain for aphid and whitefly research"

**Table S2.** Proteins sequence used for the phylogenetic trees in Figures 1 and S1

>SpenASAT1_ALU64004.1

MSASRLVSLSRKIIKPSSPTPLSHRIHKLSLMDQMGTHSYMAALLFYPKQNTTTSMPEPT

KISQVLEKSLSKVLTSYYPFAGQVRDNSFVECNDIGADLSQVRIDCPMSSIFDHPRTYID

NLVLPFDPWFPSTDSLVAAKLCHFECGGVALGVCLSHKVSDGYSLGKFLKDWSMVARDSE

AKLSLLFNGSSIFKPSNSSSFQVVADPPLYKNESKRFHFTASNLKSLKSLISSSADSATQ

ICPTTVEAITAFIYRCVSTPQSLLIQAVDQRGTSNDALVPADLTGNAILPFVVSATNKEE

MNLERLVSELRKGKEKIQDMLKYIESEEFLCSMVSDIARELNERTSNNDIPMYRFSSLRR

FPSNDINFGWGRPRQVDFSTYPMNMFILMDNQNGDGVEVIASLQDGELSALERNDEFLQF

ASPCLGF

>NattASAT1_A4A49_42350(NattASAT3_OIT19390.1)

MYASNLVSSVCKKIIKPYFPTPTSLRCHKLSYLDQMVGGIYEPFAFLYPKMSDTWSDKPS

NVISEHLENSLSKVLSHYYPFAGKLNENISVDCNDRGVEFFTTKIDCPMAKILKDPYSDK

EDLVYPKGIPCTYSYEGSLAVVQLSYFNCGGIAVSVCLTHKIGDGYTIANFIRDWATIAR

NPSSKIQSPQFNAASIYPPTKDMANLPEVVPKGEECSSKSFTFSSSKLAALRDMVISNSE

VQNPTTTEIVSAFIYLRAMATKKKTSGSICPSALVHAVSLRPPLPKTLMGNIGSFFSILT

TEEKEMDLPKVVGKLRAAKEELRQKYKNAKTDEFLPITIELYKEANNSFFNNSCYDIYRF

SSISKFPIYDIDFGWGKPEKVSFVSNGPVKNMFLLIDNKSRDGMEVITYMKEQDMSAFER

DEELLEFTSPSN

>SsinASAT1_ART34011.1

MVASALVSLSKKIIRPFSPTPFSERIYKLSFIDQFNSTQYIPLVFFYPKNKGNIEPSDMC

KVFENSLSKTLAAYYPFAGTLRDNICVECNDIGADFFKARFDCPMSEILKSHNRDVKESV

YPKDIPWSDASNKKLVTVQFNQFDCGGIALSACISHKIGDFCTVINFFHDWAAIARDSNG

KVCPQFIGSSIFPPTNEPVNEPPRIKSCVTKKLVFSNHTLKSFITESSSLEVKEPTRVEI

LTSLLYKCGMKAVLENNSSSSSIFNPSILFQTVNLRPFIPLPENTAGNFSSSLFVPTYIE

EETKLSTLVSQLRKGKEEVINNFKKCEGGQDLVSATKGPFQEIRKLYKEVDFQLYRCSSL

ANYPIYDVDFGWGRPKQITMADSPLRNTFSLFDDNTGNYIEALVALDDEITMSTFEREME

QLLESQLPTEENKTEASAHC

>SnigASAT1_ART34015.1

MSASRLVLLSKKLIKPSSPTPLSHRIHNLSLMDQMGTHSYMAAMLFYPKQDTTTSMPEPT

KISQILEKSLSKVLTSYYPFAGHVKDNSFVECNDMGADLSQVRIDCPMSSIFDQPRTYID

NLVLPIDPWFPSTDSLVATQLCHFECGGIALGACLSHKVSDGYTMSRFLRDWAMVARDTE

AKPSPLFNGASIFKPSNSSSFQVVKDPPLNKNVSKRYHFSASKLKALKSMIDSADSTSQI

RPTTVETITAFLSKCVNTPTFTPSLLVQAVNLRGTSNDALVPADLTGNAALPFVVSATNK

EEMNLQRLVIELRKGKEKIQDMLKYIESEELLCSKVSEIAREMNERTSNNDVSIYRFTSL

RKFPVNDMNFGWGRPRKVDVATYPINMFVLMDNQNGDGLEVIVNLDEGKMSAFERNDELL

QFASPCSGL

>PaxiASAT1_AOR06331.1

MAAPALVSLSKKIIKPFSPTPFSERVYKFSFLDQFNSTQYVPLVFFYPKNKGIEPSDMCK

VIENSLSKALAAYYPFAGTLKDNVCVECNDNGADFFKARFDCPMSEILKSHDRDVKEIVF

PKDMPWNVIAPNRKLLTVQFNQFDCGGIAVSACMSHKIGDMCTFNKFLHDWAAIARDSNE

NVSPQFIGSSIFPPTNEPLTEPLREQCVTKRLLFSNHTLRSLATDSEVKNPTRVETLTAL

LYKCAMKANYSSSFKSSILFQTINLRPIIPLPDNTPGNLSSSFFVPTYNEEEMKLSRLIS

QLRKEKEELFANFRKCKGNEDLVSATMGPFQQIRKLLKEGDFQLFRCSSLVNYPLYDVDF

GWGSPGKVIMADFLSRNFFTLLEDKTGEHLVAQVCFDNEITMSAFVREMEQLIEFELPSQ

EIYETQARC

>SlycASAT1_M82_NP_001316332.1

MSASRLVSLSRKIIKPSSPTPLSHRIHKLSLMDQMGTHSYMAALIFYPKQNTTTSMPEPT

KISRVLEKSLSKVLTSYYPFAGQVRDNSFVECNDIGADLSQVRIDCPMSSIFDHPRTYID

NLVLPFDPWFPSTDSLVAAKLCHFECGGVALGVCLSHKVSDGYSLGKFLKDWSMVARDSE

AKLSLLFNGSSIFKPSNSSSFQVVADPPLYKNESKRFHFSASNLKSLKSLISSADSATQI

CPTTVEAITAFIYRCVSTPQSLLIQAVDQRGTSNDALVPADLTGNAILPFVVSATNKEEM

NLERLVSELRKGKEKIQDMLKYIESEEFLCSKVSDIARELNKRTSNNDIPMYRFSSLRRF

PSNDINFGWGRPRQVDISTFPINMFILMDNQNGDGVEVIASLQDGELSALERNDDFLQFA

SPCLGF

>NbenASAT1

MAASALVSLSKKIIKPFSPTPFSERIYKFSFIDQFNSTQYIPLVFFYSKNKGNVVTPSIE

PSDMCKVIENSLSKTLAAYYTFAGTLRDNVHVECNDIGADFFKARFDCSMSEILKCLDKN

VKEMVYPKGIPWNVVTSNRKLVTVQFNQFDCGGIALSTCVSHKIGDMCTMYNFLQDWAAI

ARDSNLKLCPQYIGSSIFPPTNEPVTEPPIQKCLTKRLVVSNHTLRSLVSELSHVKNPTR

VELLTTLLYKCGMKANSSSFKPSILFQTVNLRSVIPLPNNTAGNFSSSIFVPTYNEEEMK

LSRLVSQLRKEKEQLVANYKKCKGDQDLVSTTIGPFQEIRKLFKDMDFDMYRCSSLANYP

IYDIDFGWGKPNKISIADGVFRNVFLLYDNKTSDKLEAYVCLDEERTMSAFLKEMEQFLE

FEISSEEIKMEA

>SperASAT2_LA1278_ALU64007.1

MSSVSRLVSSVCKKIIKPYSPTPISLRCHKLSYLDQMVGGIYMPLALFYPKLSNTWSNKV

VSQHLEKSLSKVLTNYYPFAGKLNDNISIDCNDNGVEFFITKIDCPMSEIFNHHYFEKHN

LVYPSQVINNQYTYEGSLAVFQLTHFSCGGIAISMCLSHKVGDGYTFGNFMNHWATIARN

PLSSEIYPISPKFDGSFYFPPTKDEDSSNVSNNNIVPQREECVSKSFSISSSKLTALKAR

VINDSEVQNPSDTEVVSAFLYQRAMATKKLVNSSSICPSLLHQAVNLHPPLPKHTMGNIC

SLFSILTKEEEEMDLARVVSKLRKEKEEVKQKYKNAKIEELLPITLEQHRKANDLLVNNS

CCDLYRFSSLITFPSYDVDFGWGKPEKVISPISTSNPPIKNMFVLMSDKNRDGVNLTCSM

KKQDMLAFERDEELLQFASPTS

>SlycASAT2_M82_ALU64015.1

MSSSVSRLVSSVCKKIIKPYSPTPISLRCPKLSYLDQMVGGIYIPLALFYPKLSNTWSNK

PNNVVSQHLEKSLSKVLTNYYPFAGKLNDNISIDCNDNGVEFFVTEINCPMSEIFNHPYF

EKHNLVYPTEVINNQYTYEGSLAVFQLTHFNCGGIAISMCLSHKVGDGYTFGNFMNHWAT

IARNPLSSEIYPISPKFDGSFYFPPAKDEDSSNVSNNNIVPQREECVSKGFSISSSKLTA

LKARVINDSEVQNPSDTEVVSAFLYQRAMATKKLVNSDSIRPSLLHQAVNLRPPLPKHTM

GNICSLFSILTKEEKEMDLARVVSKLRKEKEEVKQKYKNAKIEELLPITLEQHRKANDLL

VNNSCYDLYRFSSLITFPSYEVDFGWGKPEKVISPISTSNPPIKNMFFLMADKNRDGVNL

TCSMKKQDMLAFERDEELLRFASPTS

>SsinASAT2_ART34012.1

MAASRLVLLSKKMIKPSSPTPISHRIHKLSLMDQMGTHSYVPFSFFYPKQDAASSLEPTK

VSRILEKSLSKVLTSYYPLAGRVRDNSFVECNDMGVDFSQVQIDCPMSRIFDVPRAEMDN

LIFPKDPWIPSTDSLVVAQLNHFECGGVVLSTCISHKVADGYSIAKFLRDWAIVARDSGE

KPSPLFNGASILKPTNNSSTQVVDPSLYKTTSKRYHFSASKLKALKAMISADSGNQIVPT

AVEAVTALLFKSINARNFKPSMLMQTVNLRGKNNDALLPADLVGNVIFPFVVSAANEDEL

NLQRLVSELREGKEKIHNTLECIKSEELLCSKVCEIAREINKRTTRDDFSMYRFTSVRKF

PLNDIDFGWGRPRRADFATAPINLIFLMDDQSGDGVEVLLNLGEEEMSTFQSNDELLQFA

SPS

>SarcASAT2_LA2172_ALU64013.1

MSSSVSRLVSSVCKKIIKPYSPTPISLGCHKLSYLDQMVGGIYIPLALFYPKLSNTWSNK

PNNVVSQHLEKSLSKVLTNYYPFAGKLNDNISIDCNDNGVEFFITKINCPMSEIFNHPYF

EKHNLVYPTEVINNQYTYEGSLAVFQLTHFNCGGIAISMCLSHKVGDGYTFGNFMNHWAT

IARNPLSSEIYPISPKFDGSFYFPPTKDEDSSNVSNNNIVPQREECVSKGFSISSSKLTA

LKARVINDSEVQNPSDTEVVSAFLYQRAMATKKLANSDSIRPSLLHQAVNLRPPLPKHTM

GNICSLFSILTKEEKEMDLARVVSKLRKEKEEVKQKYKNAKIEELLPITLEQHRKANDLL

VNNSSYDLYRFSSLITFPSYEVDFGWGKPEKVISPISTSNPPIKNMFVLMADKNRDGVNL

TCSMKKQDMLAFERDEELLRFASPTS

>ShabASAT2_1_LA2098_AUG68751.1|SpenASAT2_LA1367_AUG68770.1

MSSVSRLVSSVCKKIIKPYSPTPISLRCHKLSYLDQMVGGIYLPLALFYPKLSNTWSNKP

NNVVSQHLEKSLSKVLTNYYPFAGKLNDNISIDCNDSGVEFFITKINCPMSEIFNHPYFE

KHNLVYPTEVINNQYTYEGSLAVFQLTHFSCGGIAISMCLSHKVGDGYTFGNFMNHWATI

ARNPLSSEIYPISPKFDGSFYFPPTKDEDSSNVSNNNIVPQREECVSKGFSISSSKLTAL

KARVINESKVQNPSDTEVVSAFLYQRAMATKKLVNSSSIRPSLLHQAVNLRPPLPKHTMG

NIGCLFSILTKEEKEMDLAQVVSKLRKEKEEVKQKYKNAKIEELLPITLEQYRKANDLLV

NNSCYDLYRFSSLITFPSYDVDFGWGKPEKVISPLSTSNPPIKNMFVLMDDKNRDGVNLT

CSMKKQDMLAFERDEELLRFASPTS

>SneoASAT2_LA2133_ALU64008.1

MSSSVSRLVSSVCKKIIKPYSPTPISLGCHKLSYLDQMVGGIYIPLALFYPKLSNTWSNK

PNNVVSQHLEKSLSKVLTNYYPFAGKLNDNISIDCNDNGVEFFITKINCPMSEIFNHPYF

EKHNLVYPTEVINNQYTYEGSLAVFQLTHFNCGGIAISMCLSHKVGDGYTFGNFMNHWAT

IARNPLSSEIYPISPKFDGSFYFPPTKDEDSSNVSNNNIVPQREECVSKGFSISSSKLTA

LKARVINDSEVKNPSDTEVVSAFLYQRAMATKKLVNSDSIRPSLLHQAVNLRPPLPKHTM

GNICSLFSILTKEEKEMDLARVVSKLRKEKEEVKQKYKNAKIEELLPITLEQHRKANDLL

VNNSSYDLYRFSSLITFPSYEVDFGWGKPEKVISPISTSNPPIKNMFVLMADKNRDGVNL

TCSMKKQDMLAFERDEELLRFASPTS

>SgalASAT2_LA1401_ALU64010.1

MSSSVSRLVSSVCKKIIKPYSPTPISLRCPKLSYLDQMVGGIYIPLAHFYPKLSNTWSNK

PNNVVSQHLEKSLSKVLTNYYPFAGKLNDNISIDCNDNGVEFFVTEINCPMSEIFNHPYF

EKHNLVYPTEVINNQYTYEGSLAVFQLTHFNCGGIAISMCLSHKVGDGYTFGNFMNHWAT

IARNPLSSEIYPISPKFDGSFYFPPTKDEDSSNVSNNNIVPQREECVSKGFSISSSKLTA

LKARVINDSEVQNPSDTEVVSAFLYQRAMATKKLVNSDSIRPSLLHQAVNLRPPLPKHTM

GNICSLFSILTKEEKEMDLARVVSKLRKEKEEVKQKYKNAKIEELLPITLEQHRKANDLL

VNNSCYDLYRFSSLITFPSYEVDFGWGKPEKVISPISTSNPPIKNMFFLMADKNRDGVNL

TCSMKKQDMLAFERDEELLRFASPTS

>SchiASAT2_LA1969_ALU64012.1

MSSVSRLVSSVCKKIIKPYSPTPISLRCHKLSYLDQMVGGIYMPLALFYPNLSNTWSNKP

NNVVSQHLEKSLSKVLTNYYPFAGKLNDNISIDCNDNGVEFFITKINCPMSEIFNHPYFE

KHNLVYPSQVINNQYTYKGSLAVFQLTHFNCGGIAISMCLSHKVGDGYTFGNFMNHWATI

ARNPLSSEIYPISPKFDGSFYFPPTSSSNVSNNNIVPQREECVSKGFSISSSKLNALKAR

VINESKVQNPSDTEVVSAFLYQRAMATKKLVNSSSIRPSLLHQAVNLRPPLPKHTMGNIC

SLFSILTKEEEEMDLARVVSKLRKEKEEVKQKYKNAKIEELLPITLEQHRKANDLLVNNS

CCDLYRFSSLITFPSYDVDFGWGKPEKVISPISTSNPPVKNMFVLMADKNRDGVNLTCSM

KKQDMLAFERDEELLQFASPTS

>PaxiASAT2_AOR06332.1

MAASRLALLSRKIIKPSSPTPLSQRIHKLSLMDQMGTHSSAPFVFFYPKQETARSLEPAK

VSQIFEKSLSKVLTAYYPFAGRVKDNSIVECNDMGVDLSEVHIDCPMSTIFHHPRTYVDD

LIFPKGIPWSSPKDSCLVVAQLSHFECGGIALSACISHKVSDGYSTINFLRDWAMVARNS

DAKPSPLFNGASIFKPVNFSSPPVVVDPPLKQNVSERYHFSATKLKSLKALISADSKSQN

IPTTVEAITAFLCKCVNTPTFRPSLLIQAVNIRGTNNDALVPADLVGNAFLPFAVSAANK

EELNYQRLVDELRNGKEKIRDTLKYNKSEELLCSRVPEIARELNKQTSSNDISIYRFTSL

RKFPFHDINFGWGMPRKVDLATVPVNMFILLDNQSGGGLDVLVNLEQGEMSAFQSNDELL

HFASPSSGL

>SpimASAT2_LA1578_ALU64006.1

MSSSVSRLVSSVCKKIIKPYSPTPISLRCPKLSYLDQMVGGIYIPLALFYPKLSNRWSNK

PNNVVSQHLEKSLSKVLTNYYPFAGKLNDNISIDCNDNGVEFFVTEINCPMSEIFNHPYF

EKHNLVYPTEVINNQYTYEGSLAVFQLTHFNCGGIAISMCLSHKVGDGYTFGNFMNHWAT

IARNPLSSEIYPISPKFDGSFYFPPTKDEDSSNVSNNNIVPQREECVSKGFSISSSKLTA

LKARVINDSEVQNPSDTEVVSAFLYQRAMATKKLVNSDSIRPSLLHQAVNLRPPLPKHTM

GNICSLFSILTKEEKEMDLARVVSKLRKEKEEVKQKYKNAKIEELLPITLEQHRKANDLL

VNNSCYDLYRFSSLITFPSYEVDFGWGKPEKVISPISTSNPPIKNMFFLMADKNRDGVNL

TCSMKKQDMLAFERDEELLRFASPTS

>NattASAT2_A4A49_04553(NattASAT3_OIT36748.1)

MAISRLVSLSRKIIKPSSPTPFSHRIHKLSLMDQMGTHTYMPFSFFYPKQDTASSLELTK

VSQILEKSLSKVLTAYYPFAGRLRDNSFVECNDMGVDLSQVRIDCPMSSIFNHPRTDIDK

LIFPKDPWNPSTGSLIVAQLNHFECGGIILSTCISHKVADGYSMTNFLTDWALVARDSEA

KPSPLFNGASVFQPANYSAPPQVADPSLKQNASKRYHFSASKLKALKARSLIPPTTVEAV

TAFLCKCANTPTSKPSLFIQAVNLRGTNNDALVPAGLVGNAILPYVASAANEEDLKWQRL

IGELRERKEKVRDMLKDIKPEEVLSSRVSELATQINEGTSSNDFSIYRFSSLRKFPFHDI

NFGWGRPTRVDVATFPVNMFLLLDNQNGDGVEVVVNLEEGEMSVFESNEELLQFASPSSG

L

>SpenASAT2_LA1649_AUG68772.1|SpenASAT2_LA1911_AUG68773.1

MSSVSRLVSSVCKKIIKPYSPTPISLRCHKLSYLDQMVGGIYLPLALFYPKLSNTWSNKP

NNVVSQHLEKSLSKVLTNYYPFAGKLNDNISIDCNDSGVEFFITKIDCPMSEIFNHPYFE

KHNLVYPTEVINNPCTYEGSLAVFQLTHFSCGGIAISMCLSHKVGDGYTFGNFMNHWATI

ARNPLSSEIYPISPKFDGSFYFPPTKDEDSSNVSNNNIVPQREECVSKGFSISSSKLTAL

KARVINESKVQNPSDTEVVSAFLYQRAMATKKLVNSSSIRPSLLHQAVNLRPPLPKHTMG

NIGCLFSILTKEEKEMDLAQVVSKLRKEKEEVKQKYKNAKIEELLPITLEQYRKANDLLV

NNSCYDLYRFSSLITFPSYDVDFGWGKPEKVISPLSTSNPPIKNMFILMADKNRDGVDLT

CSMKKQDMLAFERDEELLQFASPTS

>NbenASAT2

MAISRLVSLSQKIVKPSSPTPSSQRIHKLSIMDQMGTHTYMPLSLFYPKQDTASSLEPTM

VSQILEKSLSKVLTAYYPFAGRVRDNSFVECNDMGVDLSQIRIDCPMSSIFNHPRTDIDK

LIFPKDTWSTSIDSLFVAQLNHFECGGIVLSACISHKVADAYSMINFLTDWALVARDSEA

KPSPLFNGASIFQSANYSAPPLVADPSPKQNASKRYHFSASKLKALKARSQIPPTTVEAV

TAFICKCANTPTFKPSLLIQAVHLRGTNNDALVPAGLVGNAVLPYLVSAANEEDLKLQRL

IGELREGKEKVRHMLKDIKSEDVLYSRVSELATQINERTSSNDFSIYRFSSLRKLPFHDL

NFGWGRPKIVDIATSPMNMFYLLENQNGDGVDVLVNLEEGEMSLFESNEELLQFASLS

>Scor_ASAT2_LA0107_ALU64011.1

MSSVSRLVSSVCKKIIKPYSPTPISLRCHKLSYLDQMVGGIYMPLALFYPKLSNTWSNKV

VSQHLEKSLSKVLTNYYPFAGKLNDNISIDCNDNGVEFFITKIDCPMSEIFNHHYFEKHN

LVYPSQVINNQYTYEGSLAVFQLTHFSCGGIAISMCLSHKVGDGYTFGNFMNHWATIARN

PLSSEIYPISPKFDGSFYFPPTKDEDSSNVANNNIVPQREECVSKSFSISSSKLTALKAR

VINDSEVQNPSDTEVVSAFLYQRAMATKKLVNSSSIRPSLLHQAVNLHPPLPKHTMGNIC

SLFSILTKEEEEMDLARVVSKLRKEKEEVKQKYKNAKIEELLPITLEQHRKANDLLVNNS

CCDLYRFSSLITFPSYDVDFGWGKPEKVISPISTSNPPIKNMFVLMSDKNRDGVNLTCSM

KKQDMLAFERDEELLQFASPTS

>HnigASAT2_ARR28781.1

MAAPRLVLLSKKIIKPSSPTPLSHRIQKLSLMDQMGTHSYSPFSFFYPKQDTASSPEPTKVFEILEKSLS

KVLTAYYPFAGRIRDNSLIECNDMGVELSHVRIDCPMSTIFNHPHTDIDNLIFPDDPWFPSTESLVVAQL

SHFKCGGVVLGASFSHKVSDGLSTIKFLRDYAMVARNSEAKPSPLFTGASIFQPIKFSSSSPVIVDPPRK

LNASKRYHFSTSKLKALKAFVSADSKSQILPTTVEAVTAFLCKCVNTPTFKPSLLVQAVNLRGTNNDALV

PADLFGNAVLPFAVSAANKEEINLQRLVGELRKGKEKIQDTLKYVKSEELLCSTVSEIAREMNEQTSSND

MPMYRFTSLRKFPLHDINFGWGRPRRVDMATYPVNMFVLMDNLSGDGVEVLVNLEEGEMSAFESNNELHQ

FASPFSGL

>SarcASAT3_1_LA2172_AUG68798.1

MASSTIISRKMIKLLSPTPSSLRCHKLSFMDHINFPLHSPYAFFYPKIPQNYSNKISQIL

ENSLSKVLSFYYPLAGKINNNYTCVDCNDTGAEYLNVRIDCPMSQILNHPYNDVVDVVFP

QDLPWSSSGLTRTPLVVQLSHFDCGGVAVSACTSHTIFDGYCLSKFINDWASTARNMDFK

PSPKFNASTFFPLPSETNLSGALPTTRPSKHHVSRMYNFSSSNLTRLKDIVTKESHVKNP

TRIEVASALVHKCGVSMSMEKSGMFKPTLMNHTISLRPPIPLNTMGNATSIILTTTMTED

EAKLPNFAAKLRKNKQQLRDKLKDMKEDMMPLYTIELGKNAMNIIDKDTHDVYLCSSMTN

TGLHKIDFGWGEPVRVTLATHPNKNTFIFMDEQSGDGLNVLITMTKDDMLKFQSNKELLE

FASPVVESTK

>SarcASAT3_2_LA2172_AUG68799.1

MASSTIISRKMIKLLSPTPSSLRCHKLSFMDHINFPLHSPYAFFYPKIPQNYSNKISQIL

ENSLSKVLSFYYPLAGKINNNYTCVDCNDTGAEYLNVRIDCPMSQILNHPYNDVVDVVFP

QDLPWSSSGLTRTPLVVQLSHFDCGGVAVSACTSHTIFDGYCLSKFINDWASTARNMDFK

PSPQFNASTFFPLPSETNLSSTLPATRPSQRHVSRLYSFSSSNLTRLKDIVTEESHVKNP

TRVEVASALVHKCGVTMSMESSGMFKPTLMSHAMNLRPPIPLNTMGNATCIILTTAMTED

EVKLPNFVAKLQKDKQQVRDKLKDMKEDRMPLYTLELGKNAMNIIEKDTHDVYLCSGMTN

TGLHKIDFGWGEPVRVTLATHPNKNTFIFMDEQSGDGLNVLITMTKDDMLKFQSNKELLE

FASPVVESTK

>SchiASAT3_LA1969_AUG68802.1

MASSKMISRKMIKLLSPTPSSLRQHKLSFMDHINFPIHSPFAFFYPKIPQNYSNKISQIL

ENSLSKVLSSYYPLAGKINNNYTYVDCNDTGAEYLNVRIDFPMSQILNHPYNDAVDVVFP

QDLPWSSSLHRCPLVVQLSHFDCGGVAVSACTSHTIFDGYSLSKFINDWASTARNMDFKP

SPKFNASTFFPLPSETNLSSTLPTTRPSQRHVSRLYNFSSSNLTRLKDIVTKESHVKNPT

RIEVASALVHKCGVSMSMEKSGMFKPTLMNHTMNLRPPIPLNTMGNATSIILTTTMTEDE

VKLPNFVAKLHKNKQQLRDKLKDMKEDMMPLYTIELGKNAMKIIDKDTHDVYLCSSMTNT

GLHKIDFGWGEPVRVTLATHPNKNTFIFMDEQSGDGLNVLITMTKDDMLKFQSNKELLEF

ASPVVESTK

>ScorASAT3_3_LA0107_AUG68807.1

MASSTIISRKMIKLLSPTPSSLRQHKLSFMDHINFPIHTPFAFFYPKIPQNYSNKISQIL

ENSLSKVLSSYYPLAGKINNNYTYVDCNDTGAEYLNVRIDCPMSQILNHPYNDAVDVVFP

QDLPWSSSLHRCPLVVQLSHFDCGGVAVSACTSHTIFDGYCISKFINHWASTARNMDFKP

SPQFNASTFFPLPSETNLSGALPTTRPSKHHVSRMYNFSSSNLTRLKDIVTKESHVKNPT

RIEVASALVHKCGVSVSMEKSGMFKPTLMNHTMSLRPPIPLNTMGNATSIILTTTMTEDE

AKLPNFAAKLQKNKQQLRDKLKDMKEDMMPLYTIELGKNAMNIIEKDTHDVYLCSSMTNT

GLHKIDFGWGEPVRVTLATHPNKNNFIFMDEQSGDGLNVLITMTKDDMLKFQSNKELLEF

ASPVVESTK

>ScorASAT3_1_LA0107_AUG68806.1

MASSTIISRKMIKLLSPTPSSLRQHKLSFMDHINFPIHTPFAFFYPKIPQNYSNKISQIL

ENSLSKVLSSYYPLAGKINNNYTYVDCNDTGAEYLNVRIDCPMSQILNHPYNDAVDVVFP

QDLPWSSSLHRCPLVVQLSHFDCGGVAVSACTSHTIFDGYCISKFINHWASTARNMDFKP

SPQFNASTFFPLPSETNLSGALPTTRPSKHHVSRMYNFSSSNLTRLKDIVTKESHVKNPT

RIEVASALVHKCGVSVSMEKSGMFKPTLMNHTMSLRPPIPLNTMGNATSIILTTTMTEDE

AKLPNFAAKLQKNKQQLRDKLKDMKEDMMPLYTIELGKNAMNIIEKDTHDVYLCSSMTNT

GLHKIDFGWGEPVRVTLATHPNKNNFIFMDEQSGDGLNVLITMTKDEMLKFQSNKELLEF

ASPVVESTK

>ScorASAT3_2_LA0107_AUG68805.1

MASSTIISRKMIKLLSPTPSSLRQHKLSFMDHINFPIHSPFAFFYPKIPQNYSNKISQIL

ENSLSKVLSSYYPLAGKINNNYTYVDCNDTGAEYLNVRIDCPMSQILNHPYNDAVDVVFP

QDLPWSSSLHRCPLVVQLSHFDCGGVAVSACTSHTIFDGYCISKFINHWASTARNMDFKP

SPQFNASTFFPLPSETNLSGALPTTRPSKHHVSRMYNFSSSNLTRLKDIVTKESHVKNPT

RIEVASALVHKCGVSVSMEKSGMFKPTLMNHTMSLRPPIPLNTMGNATSIILTTTMTEDE

AKLPNFAAKLQKNKQQLRDKLKDMKEDMMPLYTIELGKNAMNIIEKDTHDVYLCSSMTNT

GLHKIDFGWGEPVRVTLATHPNKNNFIFMDEQSGDGLNVLITMTKDDMLKFQSNKELLEF

ASPVVESTK

>SgalASAT3_LA1401_AUG68801.1

MASSTIISRKMIKLLSPTPSSLRCHKLSFMDHINFPLHSPYAFFYPKIPQNYSNKISQIL

ENSLSKVLSFYYPLAGKINNNYTYVDCNDTGAEYLNVRIDCPMSQILNHPYNDVVDVVFP

QDLPWSSSSLTRSPLVVQLSHFDCGGVAVSACTSHTIFDGYCLSKFMNDWASTARNMDFK

PSPQFNASTFFPLPSETNLSSTLPATRPSQRHVSRMYNFSSSNLTRLKDIVTKESHVKNP

TRVEVASALVHKCGVTMSMESSGMFKPTLMSHAMNLRPPIPLNTMGNATCIILTTAMTED

EVKLPNFVAKLQKDKQQLRDKLKDMKEDRMPLYTLELGKNAMNIIEKDTHDVYLCSGMTN

TGLHKIDFGWGEPVRVTLATHPNKNNFIFMDEQSGDGLNVLITLTKDDMLKFQSNKELLE

FASPVVESTK

>SlycASAT3_M82_AJF98582.1

MASSTIISRKMIKLLSPTPSSLRCHKLSFMDHINFPLHSPYAFFYPKIPQNYSNKISQVL

ENSLSKVLSFYYPLAGKINNNYTYVDCNDTGAEYLNVRIDCPMSQILNHPYNDVVDVVFP

QDLPWSSSSLTRSPLVVQLSHFDCGGVAVSACTSHTIFDGYCLSKFINDWASTARNMEFK

PSPQFNASTFFPLPSETNLSSTLPATRPSQRHVSRMYNFSSSNLTRLKDIVTKESHVKNP

TRVEVASALVHKCGVTMSMESSGMFKPTLMSHAMNLRPPIPLNTMGNATCIILTTAMTED

EVKLPNFVAKLQKDKQQLRDKLKDMKEDRMPLYTLELGKNAMNIIEKDTHDVYLCSGMTN

TGLHKIDFGWGEPVRVTLATHPNKNNFIFMDEQSGDGLNVLITLTKDDMLKFQSNKELLE

FASPVVESTK

>SneoASAT3_LA2133_AUG68797.1

MASSKMISRKMIKLLSPTPSSLRCHKLSFMDHINFPLHSPYAFFYPKIPQNYSNKISQIL

ENSLSKVLSFYYPLAGKINNNYTYVDCNDTGAEYLNVRIDCPMSQILNHPYNDVVDVVFP

QDLPWSSSSLTRTPLVVQLSHFDCGGVAVSACTSHTIFDGYCLSKFINDWASTARNMDFK

PSPQFNASTFFPLPSETNLSSTLPATRPSQRHVSRLYSFSSSNLTRLKDIVTEESHVKNP

TRVEVASALVHKCGVTMSMESSGMFKPTLMSHAMNLRPPIPLNTMGNATCIILTTAMTED

EVKLPNFVAKLQKDKQQVRDKLKDMKEDRMPLYTLELGKNAMNIIEKDTHDVYLCSGMTN

TGLHKIDFGWGEPVRVTLATHPNKNNFIFMDEQSGDGLNVLITLTKDDMLKFQSNKELLE

FASPVVESTK

>SperASAT3_1_LA1278_AUG68803.1

MASSTIISRKMIKLLSPTPPSLRCHKLSFMDHINFHLHSPYAFFYPKIPQNYSSKISQIL

ENSLSKVLSFYYPLAGKINNNYTYVDCNDTGAEYLNVRINCPMSQILNHPYNDAVDVVFP

QDLPWSSSLHRCPLVVQLSHFDCGGVAVSACTSHTIFDGYSLSKFINDWASTARNMDFKP

SPKFNASTFFPLPSETNLSSTLPTTRPSQRHVSRLYNFSSSNLTRLKDIVTKESHVKNPT

RIEVASALVHKCGVSMSMEKSGMFKPTLMNHTMNLRPPIPLNTMGNATSIILTTTMTEDE

VKLPNFVAKLHKNKQQLRDKLKDMKEDMMPLYTIELGKNAMKIIDKDTHDVYLCSSMTNT

GLHKIDFGWGEPVRVTLATHPNKNTFIFMDEQSGDGLNVLITMTKDDMLKFQSNKELLEF

ASPVVESTK

>SperASAT3_2_LA1278_AUG68804.1

MASSTIISRKMIKLLSPTPSSLRQHKLSFMDHINFPIHSPFAFFYPKIPQNYSNKISQIL

ENSLSKVLSSYYPLAGKINNNYTYVDCNDTGAEYLNVRIDFPMSQILNHPYNDAVDVVFP

QDLPWSSSLHRCPLVVQLSHFDCGGVAVSACTSHTIFDGYSLSKFINDWASTARNMDFKP

SPKFNASTFFPLPSETNLSSTLPTTRPSQRHVSRLYNFSSSNLTRLKDIVTKESHVKNPT

RIEVASALVHKCGVSMSMEKSGMFKPTLMNHTMNLRPPIPLNTMGNATSIILTTTMTEDE

VKLPNFVAKLHKNKQQLRDKLKDMKEDMMPLYTIELGKNAMKIIDKDTHDVYLCSSMTNT

GLHKIDFGWGEPVRVTLATHPNKNTFIFMDEQSGDGLNVLITMTKDDMLKFQSNKELLEF

ASPVVESTK

>SpimASAT3_1_LA1578_AUG68795.1

MASSKMISRKMIKLLSPTPSSLRQHKLSFMDHINFPIHSPFAFFYPKIPQNYSNKISQIL

ENSLSKVLSSYYPLAGKINNNYTYVDCNDTGAEYLNVRINCPMSQIINHPYNDAVDVVFP

QDLPWSSSSLHRCPLVVQLSHFDCGGVAVSACTSHTIFDGYCLSKFINDWASTARNMDFK

PSPKFNASTFFPLPSETNLSGTLPTTRPSQRHVSRLYNFSSSNLTRLKDIVTKESHVKNP

TRIEVASALVHKCGVSMSMEKSGMFKPTLMNHTMNLRPPIPLNTMGNATSIILTTTMTED

EVKLPNFVAKLHKNKQQLRDKLKDMKEDMMPLYTIELGKNAMKIIDKDTHDVYLCSSMTN

TGLHKIDFGWGEPVRVTLATHPNKNTFIFMDEQSGDGLNVLITMTKDDMLKFQSNKELLE

FASPVVESTK

>SpimASAT3_2_LA1578_AUG68796.1

MASSKMISRKMIKLLSPTPSSLRQHKLSFMDHINFPIHSPFAFFYPKIPQNYSNKISQIL

ENSLSKVLSFYYPLAGKINNNYTYVDCNDTGAEYLNVRINCPMSQILNHPYNDAVDVVFP

QDLPWSRSSLHRSPLVVQLSHFDCGGVAVSACTSHTIFDGYSLSKFINDWASTARNMDFK

PSPKFNASTFFPLPSETNLSSTLPTTRPSQRHVSRLYNFSSSNLTRLKDIVTKESHVKNP

TRIEVASALVHKCGVSMSMEKSGMFKPTLMNHTISLRPPIPLNTMGNATSIILTTTMTED

EAKLPNFAAKLRKNKQQLRDKLKDMKEDMMPLYTIELGKNAMNIIEKDTHDVYLCSSMTN

TGLHKIDFGWGEPVRVTLATHPNKNNFIFMDEQSGDGLNVLITMTKDDMLKFQSNKELLE

FASPVVESTK

>PaxiASAT3_AOR06333.1

MGCVSNLVSSVTKKIIKPYSPTPTSLRCHKLSYLDQIIGSIFMPVALFYPKYSDRKPSID

ISEHLEKSLSKVLTHYYPFAGKLNNNVSVDCNDSGVEFYTTKIDCPMSQIINNPYADKED

VVFPKGVPCSSSYEGSPAVIQLSHFDCGGIAISTCLTHKIGDAYTMGNFLKDWATITRDP

SSKILPSPQFNGASFFPPSKDLSNLPIVVPESEECSSRNFVFSASKLAALKARVISQSQV

QNPTNTEIVSSFIYQRVMAINKRTLGVTRPSLLVQGASLRPPLPVNTMGNIGSFFSVLTT

EEKEMDLPRVVGKLRKAKEEYRQKYKHAKTNEYLPITTGLYRDADEYFTNNCYDIYRFTS

VTNFPFYDVDFGWGKPERYSPSSYGPVRNMFCLLGNKGRDGVEVISYMKEQDMLALERDE

ELLEFSSPST

>SsinASAT3_ART34013.1

MGGVSNLVSSSCKKIIKPYFPTPTSLRCCKLSYLDQIVGRIYAPMAFFYPKRSDIIKPSV

NISQHLQNSLSKVLTHYYPYAGKLNDNVSVDCNDSGVEFFTTKINCPMSEIVNNPYADKQ

DLVFPKGIACTPSYEGSLAVIQLSHFDCGGFAISACVTHKIGDAYTIGNFINDWATITRD

PSSKILPGPQFNGASFFPPSNDLPNESNFIVPKSKECLSRSFDFSQSKLASLKSRVINSL

PEVKNLTDTEIVSAFIYQRAMATKNAVVGSIQPSLLIQAANLRPPVPKSTMGNIGSFFSV

QTTEENEIDLPSLVGKLRKAKEEYRQKYKHAKTNELLPITIGLYRDADEFLVNNCYDIYR

FSSIKKFPLYDVDFGWGKPERVSLPSNCPVRNFFALVDNKARDGMEVIMYMKEQDILALD

KDEELLEFASPST

>SlycASAT4_M82_AFM77971.1

MNCYIEIQSRKMVKPSAPTPDNLRRLKLSLFDQMDIGAYVPIVFNYLPNSTSSYDHDDKL

EKSLSETLTKFYPFAGRFRKGIDPFSIDCNDEGIEYVRTKVNADDLAQYLRGQAHNDIES

SLIDLLPVMHRLPSSPLFGVQVNVFNNGGVTIGIQILHMVSDAFTLVKFVNEWAHTTLTG

TMPLDNPGFGQLPWLFPARALPFPLPDFNTTTAPNYKNVTKRFLFDALAIENLRNTIKAN

DMMMKQPSRVVVVMSLIWKVLTHISSAKNNGNSRDSSLVFVVNLRGKLSCTAPSLEHVVG

NCVIPATANKEGDEARRKDDELNDFVKLVRNTIRDTCEAIGKAESVDDISSLAFNNLTKC

IEKILHGDEMDFYSCSSWCGFPWYEADFGWGKPFWVSSVSFGHHGVTNLMDTKDGDGIQV

TICLKENDMIEFERDPHILSSTSKLAFHSLG

>PaxiASAT4_AOR06334.1

MHVPVLLRAAFHKGFQVEFIRRFSTAKTGTPQATLIEKTMIKPMHPTLFFGKNHKLSFLD

QISDGRYIPIAFFFPKQDSSSSNTKSYISQLLKDSLSKVLVSYYPFAGTLMDDTNIHCND

QGVEFQEVQFDCPLSEFFDCPHTDFESLLFPKGLPWAPCDKNRLFSIQLNHFDCGGVVLS

LCMTHTLGDAWSAFNFTKDWASLSRGASVKLCPQFDGASIFPPIDDYSAIKDIKAPGGQC

ISKRYVFSASKIQSLKDKIASTTDEYSKLRNYPTSVEAISTVLLKSAASNSNKPSLLFLT

TDLRRRAKDIIPPKCIGNTGVPLYIFAREKREMELQRLVGEFKKEKERVSSAYNNVTRET

LFPTTMETIGKAVSMLAEIDKLDLYVSSSLSKLPAEDIDFGWGGPKRVIVHGGNYIQNVS

ILCDNLRGDGIEVFVQLEEQKMHALESAQEMLEFASPYGGT

>SsinASAT5_ART34014.1

MELISQEFIKPVSPTPDHLKFYEYSILDQLIGPSLYTPVLLYYPSPPDNNNNKKPEAINT

SRLKHLKKSLSQILAHYYPLAGRLRNDNTGVDCNDEGVPFLEAFVHNHRLQDILDGKRVV

TESLVPLTNYESVFPSNTLLLVQVTLFECGGMAVGISATHKILDARSLITFLTDWAALTR

QDDPNFTLTQLVPLSKIIPPANGLPPSIPVEGFISTEPCVRNIFVFRPCSIADLKIKAAS

ENILRPSRVEVVTSTIWMCLMENSKRPCLITHMVNLRKRVDPPMHDHHLGNFIGMVVAHN

DENHIANMVASLRKGISEFEKKCLKGEREALGASIVNHATESINYLVSRKDADLYKFNSW

CGFPFYDVDFGWGKPVLSSTAEGKSKNSIKLIDTKDGGVEALVCLSEEDMKVFEKNTELL

TYATLKPNSHV

>ShabASAT3_P_LA2650_AJF98595.1|ShabASAT3_P_LA2722_AJF98594.1

MASSTIISRKMIKLLSPTPPSLRCHKLSFMDHINFHLHSPYAFFYPKIPQNYSSKISQIL

ENSLSKVLSFYYPLAGKINNNYTYVDCNDTGAEYLNVRIDCPMSQIINDPYNDAVGVVFP

QDLPWSSSLNRSPLVVQLSHFDCGGIAVSACTSHTIFDGHSLSKFINDWASTARNMDFKP

SPQFNASTFFPLPSETNLSSTLPTTHPSKRHVSRMYNFSSSNLTRLKDIVTKESHVKNPT

RIEVASALVHKCGVTMSMEKSGMFKPTLMIHAMNLRPPIPLNTMGNAVCIILTTTVTEDE

VKLPNFVAKLQKDKQQLQDKLKDMKKDMMPLYTLELAKNAMNIIEKGTHDVYGCSGMTNT

GLHKIDFGWGEPVRVTLATSKNNNFIFMDEQSGDGLNVLITLTKDDMLKFQSNKELLEFA

SPVVESTK

>ShabASAT3_F_LA2156_AJF98586.1

MASSKMISRKMIKLLSPTPPSLRCHKLSFMDHINLPLHSPCAFFYPKIPQNYSNKISQIL

ENSLSKILSFYYPLAGKINNNYTYVDCNDTGAEYLNVRINCPMSQIINNPYNDAVGVVFP

QDLPWSSSLNRSPLVVQLSHFDCGGIAVSACTSHTIFDGYCVSKFINDWASTARNMDFKP

SPQFSASTFFPLPSETNLSSTLPTTHPSKRHVSRMYNFSSSNLTRLKDIVTKESHVKNPT

RIEVASALVHKCGVAMSMEKSGIFKPTLMSHAMNLRPPIPLNTMGNATCIILTTTVTEDE

VKLPHFVAKLQKDKQQLRDKLKDMKKDMMPLYTLELGKNAMNIIEKDTHDVYVCSGMTNT

GLHKIDFGWGEPVRVTLATSNKNNFIFMDEQSGDGLNVLITLTKDDMLKFQSNKELLEFA

SPVVESTK

>SpenASAT3_LA0716_AJF98583.1

MASSTIISRKMIKLLSPTPPSLRCHKLSFMDHINFHLHSPYAFFYPKIPQNYSSKISQIL

ENSLSKVLSFYYPLAGKINNNYTYVDCNDTGAEYLNVRIDCPMSQIINDPYNDAVGVVFP

QDLPWSSSWNRSPLVVQLSHFDCGGIAVSACTSHTIFDGHSLSKFINDWASTARNMDFRP

SPQFNASTFFPLPSETNLSSTLPTKHPSKRHVSRMYNFSSSNLTRLKDMVTKESHVKNPT

RIEVASALVHKCGVTMSMEKSGMFKPTLMTHAMNLRPPIPLNTMGNAVCIILTTTVTEDE

VKLPNFVAKLQKDKQQLRDKLKDMKKDMMPLYTLELGKNAMNIIEKDTHDVYVCSGMTNT

GLHKIDFGWGEPVRVTLATSNKNNFIFMDEQSGDGLNVLITLTKDYMLKFQSNKELLEFA

SPVVESTK

>CanASAT3-1_RNaky_ACO48259.1

MAFALPSSLVSVCDKSFIKPSSLTPSTHRFHKLSFIDQSLSNMYIPCAFFYPKVQQRLED

SKNSDELSHIAHLLQTSLSQTLVSYYPYAGKLKDNATVDCNDMGAEFLSVRIKCSMSEIL

DHPHASLAESIVLPKDLPWANNCEGGNLLVVQVSKFDCGGIAISVCFSHKIGDGCSLLNF

LNDWSSVTRDHTTTTLVPSPRFVGDSVFSTKKYGSLITPQILSDLNECVQKRLIFPTDKL

DALRAKVAEESGVKNPTRAEVVSALLFKCATKASSSMLPSKLVHFLNIRTMIKPRLPRNA

IGNLSSIFSIEATNMQDMELPTLVRNLRKEVEVAYKKDQVEQNELILEVVESMREGKLPF

ENMDGYENVYTCSNLCKYPYYTVDFGWGRPERVCLGNGPSKNAFFLKDYKAGQGV

>NattASAT3_OIS95899.1

MGVASRLLSTISKKTIKPFSPTPSTQRLHKLCLPGQAFTPNHIPLVFFYPKQQVDTIFPNGHKQQLSKFL

EISLSKTLTCYYPWAGRLKDNATVECNDVGAEYFEVQINSSMDEVLHHSDSRIKDLVTFPQGAPWGSCTD

RALAIAQLSHFECGGIAISICLSHKVGDGCSCYWFLRDWASLTRDPNTKIASSPYFVEDSIIPSPLPDGP

LVSPVSKKKQECIEKTFDFSALKLSALKAFVEAESGVQNPTRNEVVNALVYKCAANAASIHQSHQLVQYS

NIRQSVSPPLPPTSIGNLLTVFSTPIYSSSREGDFNLSKVVSDIRNSKQKLLSIDIIKENEWALEILNAY

KTGKGTFQQRNCDVYVCTSLCQFPFQDLDFGWGKPSRARLGGHFISKYFSLISTHDRGIQASVHLNEQQM

SVFEKDEHLLRFATPTGQD

>NattASAT3_OIS97679.1

MVASSLISLISKKIIKPSIPTPLRIHKLSFVDQVFHYYIPVSFFYPKNGNDKSYEPTHVSQLLEKSLSKT

LSFYYPYAGRLKDSSTVDCNDAGVELSHVRVHCPMSEIFNHPYTNAEHVVFPVAQPHGHDEGNLAITQVS

HFDCGGIAVGGCLSHKIGDGCTASSFFYNWGVLSRDISANLSRPHFVEDSIYPQSSDPSIVSKSIKSEEP

VGYLGKRFVFPAAKVNALKAIFSMDSGVCNPTRTEVVTALLYKCAVAARRVNFGSFRPSSLFQVADMRQR

LNPSLSHDASGNIISGFLVPITNEREMNCPRLVSEMRKEKEKLVDKDNIKKNAFISKVLDSLEKGKNNEF

DDYFCSSVIALPLYKTDFGWGRPAKVSMATGPISKLFFLMDNQSGGGVEALVMLDKQDMSIFEKDPELLE

FAIPAPNL

>NattASAT3_OIS99435.1

MVGLSTIISKKIVKPLSPTPSTQRWHKLSLLDQGIGNFYMGLVLFYPKHQVDTFQIEPKQLSTLLENSLS

KAITYYYPWAGSLRGNATIDCDDTGAEFLEVEVNCPMDKVLYHQDSSIKNIAFPQGVPLRNAADGACLIV

SQLSHFECGGIAISVCLSHKVGDARSALFFVRDWAVLTRDPNAELVCPPYFVQDSLIPSPSDGPLAFPVI

VSKPEERIELEKRFYLSASKIRALKDLVAAESGVENPTTTEVVSALVYKSAAMFNLGSSHEQSRLILLSD

IRKEISHSVPSTSIGNILTVFSTPIYNNKEDLKVSTLVADIRKSKHELSSRDNFKENEWASGILHAYNTG

DGTYRQSNCDVYRCSSASNIPFQDLDFGWGKPIRATMPSSPFSKMFYMMSTHDRGIEVIINLNEQQMSAF

ENDKDLLEFATPAHVED

>NattASAT3_OIT03546.1

MASIAQSPIGTTTEKINKPFSATTISLPLVSIISEKLIKPSSPTPPTKRWHKLSLIDQAFSNSYIPFSLF

YTKQQLDAISKNNNPTQISHLLEESLSKILSTYYPYAGRLKDNTVVDCNDTGAEFIEVQISCPISQTLNW

HNSAVEDLLFPQGLPWSNSADRGLVVVQLSYFSCGGIAISMCISHKTGDGCSGYNLFRDWSEITSDPNFS

KPSLHYVEQSIFPPLSSGPFLSPLFMSNKHDCVQRKYIFSNEKLLNLKNKVAAESEVQNPTRTEVVSALI

FKRAVAAAKANLGFFQPSSMVQAVDLRAQIGLPPNAIGNLLTICPTSIITEQSMTISKLVSEIRKSKELA

YNKDNINDNIFVALLLELANSKQEYHDNGPNAYQITSLVKFALHEIDFGWGKPTKVSIANGLNNKLAILM

GNQSGGLDAFVTLSEQDMSVFERDPELLEFASLVPSC

>NattASAT3_OIT04893.1

MSASNLVSSVCKKIIKPYFPTPTSLRCHKLSYLDQMLGGIYVPFAFFYPKLSNTWSDKPSNIISEHLENS

LSKVLSHYYPFAGKLNENVSVDCNDGGVEFFTTKINCPMAKILNDPFVDKEDLVYPKGIPYAYSYEGSLA

MIQLSHFNCGGIAISACLTHKVGDGYTMANFIHDWATIARNPSSKIQSPQFNAATIFPPTKDMINLPEVV

PKGEECCFKSFAFSSSKLAALKARVINNSNIQNPTTSEIVSAFIYQRAMATKKKTSGSICPSVLVQAMNL

RPPLPKNLMGNIFSFFSMLTTEEKEMDLPKVVGKLRAAKEELRQKYKNAKADEVLPITIELYEEAYDNFF

NNSCYDIYRFSSINKFSFYNVDFGWGKPEKVSMSSNCPVKNMIVVMDNKSRDGVEVLSYMKEQDMSALEK

DEELLEFASPSS

>NattASAT3_OIT05194.1

MAASALVSLSKKIIKPFSPTPFSERIYKLSFIDQFNSTQYIPLVFFYSKNKDNVVTPSIEPSDMCKVIEN

SLSKTLSAYYPFAGTLRDNVHVECNDIGADFFKARFDCPMSEILKSPDTNVKELVYPKDIPWNVVTSNRK

LVTVQFNQFDCGGIALSACVSHKIGDMCTVSKFLQDWAAIARDSNLKLCPQFIGSSIFPPTNEPVNEPPI

QKCVTRRLVFSNHTLRSLLSESSQVKNPTRVEILTALTYKCGMKVNSCSFKPSILFQTVNLRSLIPLPDD

TPGNFSSSLFVPTYNEEEMNLSRLVSQLRKEKEQLVANYKMCKGGQDLVSTTIRPFQEIRKLFKDMDFDM

YRCSSLANYPINDLDFGWGNPNKISLADGVFRNTFLLYDNKTGDQIEAQISLDEESTMSAFLREMEQLLE

FEILL

>NattASAT3_OIT06250.1

VKPSKPTPPHLKNHKLCFLDQIQPPTRIKIVLFYENTTLAQNQNILKNALSETLTLFYPLAGELKEGSTF

VDCNDTGVAYVEAQAKTHLSHFLENPDLSQLKKLMIDVECHLVSFQVTRFDCGGMAIGASMFHLVSDANT

MSSFLKTWANIARERIMPLASLDFTSSSLLFPPQNHLPQELSAFLKTLFFKEGDQFLQRRFVFDSKAIAA

IKMKSGSNSNVPKPTRVEALTAFIWKHMMLASEAVKGSSSPSCMLNHAVNLRPRLHPPLPNAFGNVVWFA

TTFYESEITTSDNIDLSHLVGLVREVFETFSNGENLKELEGDASFTYLVNLATDTINSFDKVDNYMFTSW

CRFGLDEVDFGWGKPIWVAPLFDPTCSLGHVQIVILVESGRSSDGIEAWI

>NattASAT3_OIT19645.1

MNNNYVDCNDIGAGYLDVRVSCPMFQVLNHAYNDATHTVYPQDLPLSSSLNRSPLMVQLTRFDCGGIAVS

LCISHQITDGCIHVSFFMIGQLQPDCWISDHLFSLMQIVSSQ

>NattASAT3_OIT20175.1

MKMPTSTILSKKLIKPSSPTPSTQKWHKLSLLDQAFSNIYIPFAFFYTKKQLDSVSIHPTQISHLLEDSL

SKLLVSYYPYAGRLKDNTIVNCNDVGADFLRVEINIPISEALQKHNNVIEEMVFPQDLPWSNGTNRGLVV

AQLSYFNCGGIAISLCISHKVGDGHSNYNFFHDWAKLTREPNGPKPVLQYVQESIFPPPITGTPFLSPVC

TSNKDDCIQRKFIFPIQKLNKLKSLVASTSNVQNPTRNEVVSALIFKCIVAAVKLLNSGKSFQPWIMSQS

FDLRHQIDLPPNTIGNILTSTYISIASEKYMTLANIVTEMRKEKQRVSDRDNVEDNLFLKKLLELATEEK

SFTKVQNNMCNFSSLVKFPVHEIDFGWGKPAKMNIANGLHNKVIFLVGNLSGGIDAFVSLSAKVMSGFEK

DKELLEFATAVPTF

>NattASAT3_OIT22778.1

MNIKVKLISTEMIQPTSPTPNHLKNFNHCLLDQLIPAPYAPIILFYPNLGDIKVHEKSALLKKSLVDTLS

RFYPLAGRFKDELSIDCNDQGVNYVTTKVNCHLIAFLNEPNLESIGQFLPCQPPFKVLAAGDYVTNIQIN

VYECGGIAIGLCIAHKILDGVGLSTFLKYWAGLACSSNEVQCPNLMANYIFPAEDLWLRDTSMTMWSSLF

KKGNFVTRRFVFNASAINNLKTMSTSSHIKHPTKVEVVSSFIWKCSIAASKQKNCLNSLLTHIVNLRKRA

APSLPENIHGNLLWLSSAKNTAKHEMGLPDLVNQVRKSISRIDDHYIKRLRSDEGFNLMRKSL

>NattASAT3_OIT27081.1

MAALSVVSLISKKIIKPSIPTPLSLRSHKLSFVDQVIHMCTPIAFFYPKVGNYAYKSYEPTQVSQLLEKS

LSKTLSFYYPYAGRLKNGSSVDCNDMGVELSTVRVHCPMSKIFKHPYTEAEHVVFPQAKPHDHNEGNLAI

AQVSHFDCGGIAIGACLSHKIGDGCTASNFLYNWGVLSRDINATPSPRFVGESIYPQSNDPSVISSIKYE

EPIGYLGKRLIFPATKVNALKAKISRESGVKNPSRTEVVSALLYKCAVAARRANNVGSFKPSTLFQLANM

RQRLNPPLSHHACGNVISGFFVKTTNERELNYPRLIREMRKEKEKLHVHENNATKNAIISEVVDSLEKGK

TPFHSSEVDDYACTSVLLFPVYNTDFGWGRPARVSMATGPFNRYFFLMDNQSGGGVEAIVMLGEQDMAIF

ERDPELLEFAIPIPNH

>NattASAT3_OIT29898.1

MAASALLSSPSLVSISDKSFIKPSTLTPSTVRLHKLSLVDQSFSNMYIPFAFFYPKQQKEESNNSQLSHI

ADLLQRSLSQTLVSYYPYAGKLRDNATVECNDMGAEFLSVQIKCPMSEILNHPHATDAESIVFPKDLPWK

NNYEGGNLLVVQLSKFDCGGIAISICLSHKIGDGCSVINFLNDWASVTRDHRFVVPSPRFVADSIFSSLN

GPLIAPHIMSSNVSECVQKRLVFPTAKLDALRAKIAVESGVENPTRSEVVSALVYKCATKAAVASTSANI

HPSKLVHYLDVRTMIKPRLPRSAIGNLLSVFSTAATQDIELPRLVHNMRKEVEVAYKKDQVQQNELLLEV

VESMKKGKLAFEEKDENCTTPTTMYFCSNLCKFPFYSVDFGWGKPERVCLGTGPFANFFFLKDYQTGRGV

EARVTLQKQHMSAFERDEELLQFASSSAPSL

>NattASAT3_OIT33147.1

MGFLCANLKNSLAVEIMSKKLVKPSSPTPTHLQSYKLSFFDQLAIRMHVPIVLIYHNLNNSITNELLEES

LSKTLTHVYPSAGRINKDRRVVDCLDQGVEFIIAKVNCQLEDFLEQARKDIDLANHFWPQGIKDVDDNYD

FAITPLVFVQVTRFECGGLALSVAAEHIAIDGFTNMKFIYEWAKVCRLGIPTSTTTDIFNYDLGDIFPAR

DTSRILKPLASLAIPKDTITYVAKRFVFNEASISKLRNKIASGVLSFKPSRVEIVTALLWRALIRASQAK

NGRLRPSLMSFPVNLRGKASLPKLSDTFGNFAVEVPVVFTPNETKMELHNLIALIRDATDKTMVSSAKAS

NDELVSMAANLYNMTQEWEANEEVDEFTCSSLCRFPMKEADFGLGKPCWMTFGLRQSQVFWLYDADFGSS

IAAQVDLNESLMHYFERDQDLNTFTILNN

>NattASAT3_OIT35573.1

MVESKILSRKMIKPFSPTPSSLSRHNLSFIDQIATCAYSPVVAFYSKPTNNTSQILENSLSKVLSTYYPF

AGRMMDNYEYVDCNDTGAELLNVRISCPMSEILNHTYNDVSDVVFPQDLPWSSSYSNGSLLVVQLSHFDC

GGIAISVCLSHKIADGYSLLKFLKDWAATTRHDLDFKPSAQFDAHSFFPPVAMNDLPAVMPDILREPQQR

VSRIYHFSSSSLDTLKDIVAMNSHVLNPTRVEVATALLHKCGAAVSMANFSVFQPSLLLHIMNFRPPLPQ

NTIGNACFFFGSIAATEDEIQLPHFVAQLRKAKQYLQDKSKDPHQLASHLLEKVKEQANKSGENKFDLYM

CSSLCNLGVYKIDFGWGEPIRVTLARNPMKNNFIFLDPPNGDGINVLITLTEADMLTFQSNKELLEFASP

LV

>NattASAT3_OIT37273.1

MDPTKISNILENSLSQTLSLYYPYAGRLKDNTIVDRNDAGAEFVQVQINSPISESLQWHNNAIEEMIFPQ

GLPWSNCTNRALVVAQISYFNCGGIVVSMCISHKVGDGHSGYNFFMDWAITSREPNLSTKPCLYYVEKSI

FPPPPNGPFLSPLCTSNKDESIHRRYTFLAKNLMNLRPQ

>NattASAT3_OIT37557.1

MAISRLVSLSQKIIKPSSPTPFSHRIHNLSLMDQMGTHSYIPFSFFYPKQDTASSLEPTKVSQILEKSLS

KVLTAYYPFAGRVRDNSFVECNDMGVDLSQVRIDCPMSSIFNHPRTDIDKLIFPKDPWNLSTGSLIVAQL

NHFECGGIILSACISHKVVDGYSMTNFLRDWALVARDSEAKPSPLFNGASVFQPANYSAPPQVADPSRKQ

NASKRYHFSASKLKALKARSQIPPTTVEAVTALLCKCANTPTFKPSLLIQAVNLQGTNNDALVPAGLVGN

AILPYVASAANEEDLKWQRLIGELRERKEKVHDMLKDIKSEDILFSRVSELATQINERTSSNDFSIYRFS

SLRKFPFHDINFGWGRPTRVDAATFPVNMFLLLDNQNGDGVEVLVNLEEGEMSVFESNEELLQFASPSSG

L

>NattASAT3_OIT39828.1

MSASNLVSSVCKKIIKPYFPTPTSLRCHKLSYLDQMLGGIYVPFAFFYPKLSNTWSDKPSNIISEHLENS

LSKVLSHYYSFAEKLNENVSVDCNDGGVEFFTTKINCPMAKILNDPFVDKEDLVYPKGIPYAYSYEGSLA

MIQLSHFNCGGIAISACLTHKVGDGYTMANFIHDWATIARNPSSKIQSPQFNAATIFPPTKDMINLPEVV

PKGEECCFKSFAFSSSKLAALKARVINNSNIQNPTTSEIVSAFIYQRAMATKKKTSGSICPSVLVQAMNL

RPPLPKNLMGNIFSFFSMLTTEEKEMDLPKVVGKLRAAKEELRQKYKNAKADEVLPITIELYEEAYDNFF

NNSCYDIYRFSSINKFSFYNVDFGWGKPEKVSMSSNCPVKNMIVVMDNKSRDGVEVLSYMKEQDMSALEK

DEELLEFASPSS

>NattASAT3_like_XP_019226719.1

MSKAITYYYPWVGSLRDNATIDCDDTGVEFLEVEVNFPMDKCEGIAISVCLSHKVGDARNTLFFVRDSPA

LTGDLNAKLVYSPFINQGSLVLSPSDSPLSFPAILSKLEECIELEKRFYFSASKIRALKDLVVAELSVEN

PTTTEVVSALVYECAVIINLGSSHEQSRLVLLSDIQKEISHSVPPTSINNILTIFTAPIYNNKGDLRVSN

LVADI

>NattASAT3_like_XP_019235181.1

MYIPIAFFYPRPLHHNINKSSKQELAQVLENSLSKSLTSYYPFAGKLKDNVAIECNDMGAKFLNVELNCS

MSDVVNLPDTGPEYLTFPKNLPWNTSYDKGSNFVVAQLSHFKCGGIAVSACLSHKLGDGSISSISAYKPP

YIAKRYVFSCSKISALKEEVMASESGVQNPTRNEVVTALLYKCIMAASRSNSGGIFKPSVLNQMVNLRPR

LNPPLPNNSAGNFVTTISIKSTSDIEQMSRARVNSRVEKGKRTTQEETVDNYW

>NtabASAT3_like_NP_001313112.1

MASIDQSPIGTTPENINKPFSATTISLPLVSIISEKLIKPSSPTPHTKRWHRLSLIDQAFSNSYIPFSLF

YTKQQLDAISKNNNPTQISHLLEESLSKILSTYYPYAGRLKDNNVVDCNDTGAEFVEAQISCPISQTLNW

HNSATEDLLFPQGLPWSNSADRGLVAVQLSYFNCGGIAISMCISHKIGDGCSGYNLFRDWSEITRDPNFS

KPSLHYVEQSVFPPPSSGPFLSPLFMSNKHDCVQRRYIFSNEKLLHLKNRVAAESEVQNPTRTEVVSALI

FKRAVAAAKANSGFFQPSSMVQAVDLRAQIGLPPNAIGNLLTICPTSITNEQSMTISKLVSEMRKSKELA

YNRDNIFVALLLELANSKQEYHDNGPNAYQITSLVKFALHEIDFGWGKPTKVSIANGLNNKLAILMGNQS

GGLDAFVTLSEQDMSVFERDPELLEFASLVPSC

>NtabASAT3_like_XP_016432602.1

MAAVPSQRISIPEKTIVKPSSPSPSSLNPILKNVRAHRKFLFSGSKLGALRAIVAVESGVKNPSRSEVVS

AIMFKFVTKAASRINNNSGTFRPSMMLNDVDIRPLVVPPLPQNSIGNLLSVFLLVVTKEDEMKIATLVCN

LRKELEAVYKKDPVKQNALILDGENQEEYHHHLVRGRIACLRGVEALVTLEEKQMLALER

>NtabASAT3_like_XP_016440226.1

MAASALLSSLVSICDKSFVKPSTLTPSTLRLHKLSFVDQSLSNMYIPVAFFYPKPQREESNNSQLSHIAD

LLQTSLSQTLVSYYLYAGILRDNATVECNDMGAEFLSVKINCPMSEILNHPHATDAESIVFPKDLPWKNN

YEGGNLLVVQLSKFDCGGIAISACLSHKIGDGCSVINFLNHWANVTRDRRFVVPSPRFVGDSIFSSLNGP

LIAPQIMSNVSECVQKRLIFPIAKLDALRAKIAVESGVENLTRAEVVSALIYKSATKAAASCSANIQPSK

LVHYLNVRTMIKPRLPRRAIGNLLSVFSTAATQDIELPRLVHNLRKEVEVAYKKDQVDENELVLEIVDSM

KKGKLPFEEKDENCTTTTMYFCSNLCKFPFYNVDFGWGKPERVYLGEXQTSSTTKCNWKSLIRVLHSSYS

VSGRGVEARVMLQKQHMSAFECDEELLEFASSSVPSL

>NtabASAT3_like_XP_016442548.1

MVESTILSKKMIKPSSPTPSSLSRHNLSFIDQIASSAFSPIVVFYSKPTNNTRQILENSLSKVLSSYYPF

AGRIKDDYEYVECNDTAISGCLSHKIADGYSLLKLLKDWAATARLELDFKPSTQFDVHSFSSSSLDRLKD

IVAMNSQVQNPIRVEVATALLHKCGSAVSIANLGVFQPSLLCHLMNFLPPLPQNTIGNACYFFGSTAAKE

NETQSQLRKAKQYIQDKSKDPNQLASHLLEKFKERANKSEKNKFDFYMCSRLCNLGVYKIDFGWGEPIRV

TLARNPMKSNFIFLDSPNGDGINVLITLTEADMLIFQSNKELLEFASPVV

>NtabASAT3_like_XP_016447325.1

MTGLSTIISKKIVKPLSPTPSTQRWHKLSLLDQGIGNFYMGLVLFYPKHQVDTFQIEPKQLSTLLENSLS

KALTYYYPWAGSLRDNATIDCDDTGAKFLDVEVNCPMDKVLYHQDSSIKNIAFPQGVPLRNAADGACLIV

SQLSHFECGGIAISVCLSHKVGDARSALFFVRDWAALTRDPNAELVCPPYFVQDSLIPSPSDGPLAFPVI

VSKPEERIELEKRFYFSASKIRALKDLVAAESGVENPTTTEVVSALVYKSAAIANLGSSHEQSRLILLSD

IRKEISHSVSSTSIGNILTIFSTPIYNNKEDLKVSKLVADIRKSKHELSNRDNFKENDWASAMLHAYKTG

EGTYRQINCDVYRCSSASNIPFKDLDFGWGKPIRATMPSSPFSKMFYMMSTHDKGIEVIINLNEQQMSAF

ENDKDLLEFATPAHVED

>NtabASAT3_like_XP_016447861.1

MTSSRLISVSEKIIKPFSATPSPLRHYKLSLIDQLMSTMYIPIAFFYRRPLEHKIINKSSKLEVSQTLEK

SLSKTLSSYYPFAGKLKDNLSIECNDMGAKFLNVELNCSMSEVVNLPDTGPEYLAFPKNLPWNSTSYYEG

TNYFVVAQLSHFRCGEIAVSACLSHKIGDGCTEINFMNDWATIARSATSSNYAISRLSTPQWIGASIFPP

TYDDHSSAISRITPNTLPYITKRYVFSSSKLSALKKFMASDSGGVQNPTRNEVVTALLYKCANMATSRSS

SGSGLFKPSALIQVVNLRPRLNPPLPNNSAGNLISSILIKSTDEEQMKVARIVQDLRKEKEQLNMKHDLV

NQNGVVLSALEYLNDTMDVYCCSSLCNYQLYNVDFGWGKPERVVVTNGTINFFLLSDDKNGDGIEVLVSL

GKEVMSEFNNNKELLEFAFPVSN

>NtabASAT3_like_XP_016447896.1

MGVLSRLHKLCLPGQAFTPSPIPLVFLYSKQQVDTIFPNGPKQQLSKFLEISLSKTLTCYYPWAGRLKDN

ATVECNDVGAEYFEVQINSPMDEVVHHSDSRIKDMVTFPQGATWGNCTDRALAIAQLIHFECGEIAISVC

LSHKVGDGCSCYWFLRDGASLTRGPNAKITSTPYFVEDSVIPSPLPDGPLVSPNIITGSLDGCFQSPLCS

HNYTLLRTLKGETIDRLLVDN

>NtabASAT3_like_XP_016451255.1

MAISRLVLLSQKIIKPSSPTPFSHRIHKLSLMDQMGTRTYMPISFFYPKQDTAISLEPTKVSQILEKSLS

KVLTAYYPFAGRVRDNSFVECNDMGVDLSQVRIDCPMSSIFNQPRTDIEKLIFPKDPWSTSTDSLIVAQL

NHFECGGIVLSACISHNVADGYSMTNFLRNWALVARDSEAKPSPLFNGASIFQPTNYSAPQVADPSRKQN

ASKRYHFSASKLKALKARSQIPPTTVEAVTAFLCKCANTPTFKPSLLIQAVNLRGTSNDALVPAGLVGNA

ILPYVVSAANEEDLNLQRLIGELREGKEKVHNMLKYIKSEELLCSRVSELATQINEQTSNNDFSIYRFSS

LRKFPFDDINFGWGRPTRVDIATFPVNMFLFLDNQNGDGVEVLVNLEEGEMSVFESNEELLQFASPSSGL

>NtabASAT3_like_XP_016453041.1

MGVASRLLSTVSKKTIKPFSPTPSTQRLHKLCLPGQAFPPNHIPLVFFYPKQQVDTIFPNGHKQRLSKFL

EISLSKTLTCYYPWAGRLKDNATVECNDVGAEYFEVQINSSMDEVLHHSDSRIKDLVTFPQGAPWGKCMD

RALAIAQLSHFECGGIAISVCLSHKVGDACSCYCFLRDWASLTRDPNTKIASSPYFVEDSVIPSPLPDGP

LASPVSKKKQECIEKTFDFSALKISPLKAFVEAESGVQNPTRNEVVNALVYKCAANAASIHQSHQLVQYT

NIRPSVSPPLPPTSIGNLLTVFSTPIYNSSREGDFNLSKIVTDIRNSKQKLLSRDNIKENEWAWEIVNAY

KTGKGTFQQRNCDVYVSTSLCQFPFHDLDFGWGKPSRARLGGHLISKFFSLMSTHDGGIQVSVHLNEQQM

SVFENDQHLLQFATPTG

>NtabASAT3_like_XP_016462941.1

MKKPSSTILSKKLIKPSSPTPSTQKWHKLSLLDQAFSNIYIPFAFFYTKKQLDSVSIHPTQISHLLENSL

SRLLVSYYPYAGRLKDNTVVDCNDVGADFLRVQINIPISQALEKHNNVIEEMVFPQDLPWSNCTNRGLVV

AQLSYFTCGGIAISLCTSHKVGDGHSSYNFFHDWAKLTREANGPKPVLQYVQESIFPPPITGTPFLSPVC

TSNKDDCIQRKFIFSLQKLNKLKTLVASTSNVQTPTRNEVVSALIFKCIVAAVNGKSFEPWIMSQSFDLR

HQIDLPPNSIGNILTSTYISIASEKYMTLANIVTEMRKEKQRVSDRDNVEDNLFLKKFLELATEEKSYSK

VQNNMCNFSSLVKFPVHEIDFGWGKPAKMNIANGLHNKVIFLVGNLSGGVDAFVSLSAKVMSAFEKDKEL

LEFATAVPTF

>NtabASAT3_like_XP_016464881.1

MAASGLLSSPSLVSICDKSFIKPSTLTPSTLRLHKLSFVDQSVSNMYIPVALFYPKPQREESNNSHLSHI

ADLLQTSLSQTLVSYYPYAGILRDNATVECNDMGAEFLSVKINCPMSEILNHPHATDAESIVFPKDLPWK

NNYKGGNLLVVQLSKFDCRGIAISACLSHKIGDGCSVINFLNHWVNVTRDRRFIVPSPRFVGDSIFSSLN

GPLIAPQIMSNVSECAQKRLVFPTAKLDALRAKIAVESGVENPTRAEVVSALLYKSATKAAASCSANIQP

SKLVHYLNVRTMIKPRLPRSAIGNLLSMFSTAATQDIELPRLIHNLRKEVEVAYKKDQVEQNELVLEIVE

SIQKGKLPFEENDENCTTPTTMYFCSNLCKLPFYSVDFGWGKPERVCLGTGPFKNFFFLKDY

>NtabASAT3_like_XP_016467639.1

MATASALLSSPSLVSICDKSFIKPSCLTPPKLRLHKLSFVDQFLSNQYIPVAFFYPKPQREESNNNSQLS

HIADLLQRSLSQTLVSYYPYAGKLRDNATVECNDIGAEFLSVQIKYPMSEILNHPHATDAESIVFPKDLP

WKNNYEGGNLLVVQLSKFDCGGIAISACLSHKIGDGSSVINFLNDWANVTRDHTFVVPSPRFVGDSIFPS

QNGPLIAPQSMSNVSECVQKRLIFPAAKVDALRAKIAVESGVENPTRAEVVSALLYKCATKSKAAASSSG

NIQPSRLVHYLNVRTMMKPRLPRSTIGNLLSVFSTAATQDIELPRMVRNLRKEVEVAYKKDQVEQNELVL

EVVESMKKGKLPFEENDENSTYFCSNLCKFPFYSVDFGWGKPERVCIANGPSKNYFFLKDYKTRRGVEAQ

VMLQKQHMSAFECDEELLQFASSSSGV

>NtabASAT3_like_XP_016469851.1

MKAAQVGHRERDNKLANAEKRRQKPSRTGAAPQKQLCGRRSRMGEVKQEPQKRYYLAKKEWHYLGLKIAV

ESGVENPTRAEVVSALLYKCATKAAAISSSGNIQPSKLVHYLNVCMMIKPRLPQSAIGNLLSMFSAAATS

TQDIELPRMVRNLRKEVEVAYKKDQVEQNELVLEVVESMKKGKLPFEENDENSTYFCSNLCKFPLYSVDF

GWGKPERVCLANGPSKNYFFLKDYKTGRGVEARVMLQKQHMSAFECDEGLLQFASSSGV

>NtabASAT3_like_XP_016474114.1

MAASALLSSLVSICDKSFVKPSCLTPPKVRYHKLSFVDQSLSNMYIPFAFFYPKQQKEESNNSQLSHLAD

LLQTSLSQTLVSYYPYAGILRDNATVECNDMGAEFLSVKINCPMSEILNHPHATDAESIVFPKDLPWKNN

YEGGNLLVVQLSKFDCGGIAISACLSHKIGDGCSVINFLNHWANVTRDRRFVVPSPRFVGDSIFSSQNGP

LIAPQTMSNVSECVQKRLIFPTAKLDALRAKIAIESGAENPTRAEVVSALLYKSATKAAASSSANIQPSK

LVHHLNVRTMIKPRLPRSAIGNLLSMFSTAATQDIELPRLVHNLRKEVEVAYKKDQVDENELVLEIVDSM

KKGKLPFEEKDENCTTTTMYFCSNLCKFPFYSVDFGWGKPERVCLGTGPFKNFFFLKDYQTGRGVEARVM

LQKQHMSAFECDEELLQFASSSLPSL

>NtabASAT3_like_XP_016477109.1

MCASNLVSSVCKKIIKPYFPTPTSLRCHKLSYLDQMLGGIYVPFAFFYPKLSNTWSDKPNNVISEHLENS

LSKVLSHYYPFAGKLNENVSVDCNDHGVEFFTTKIDCPMSKILNNPYADKEDLVYPKGIPYTDSYGSLAV

VQLSHFNCGGIAVSACLTHKIGDGYTIANFIHDWATIARNPSSKIQSPQFNAATIFPPTKDMVNRHEVVP

KGEECSFKSFAFSSSKLVALKTRVINNSNIQNPTTTEIVSAFIYQRAMATKKKTSGSICPSVLVQAMNLR

PPLPQNLMGNIGSFFSIITTEEKEMDLPRLVGKLREAKEEVRKKYKHDKTDEFLPITIDLFREANDSFFN

NSWYDIYRFSSINKFPFYNVDFGRGRPEKVSMSSNGPVKNMFLLMDNKSRDEVEVLSSMKEQDMSALEKD

EELLEFASPSS

>NtabASAT3_like_XP_016477482.1

MAVSRIISVSEKIIKASSTTPSPLKNYKLSLLDQVMSTMYIHIAFFYPRPLHHNINKSSKQELAQVLENY

LSKTLTSFYPVAGKLKDNVAIECNDDMGAKFLNVELNCSMSDVVNLPDTGPEYLAFPKNLPWKTSYDKGS

KFAVAQLSHFKCGGIAVSACLSHMLGDGCTVTNFMNDWATIARNQENQQNQGDPHNINTQVPTPRDSPRQ

SRESTPDGSQINEHAQFENVEAVDKALQKLIVAQVNKAVEALVNQLHVATSTPTPNNNTLENPCSELVNS

GSGGTPSESQEREPGNGICQIRGDQHTARAINSVADTSTRNEGK

>NtabASAT3_like_XP_016479613.1

MAASGLLSSPSLVSICDKSFIKPSTLTPSTLRLHKLSFVDQSVSNMYIPVALFYPKPQREESNNSNLSHI

ADLLQTSLSQTLVSYYPYAGILRDNATAECNDMGAEFLSVKINCPMSEILNHPHATDAESIVFPKDLPWK

NNYEGGNLLVVQLSKFDCGGIAISACLSHKIGDGCSVINFLNHWANVTRDRRFVVPSPRFVGDSILSSTS

LSQTLVSYYPYAGILRDNATVECNDMGAEFLSVKINCPMSEILNHPHATDAESIVFPKDLPWKNNYEG

>NtabASAT3_like_XP_016480914.1

MAASALVSLSKKIIKPFSPTPSSERIYKLSFIDQFNSTQYCPLVFFYPKNKGNVVTPSIEPSDMCKVIEN

SLSKTLAAYYPFAGTLRDNVHVECNDIGADFYKARFDCPMSEIVKSPDRNVKEMVYPKGIPWNIVTSNRK

LVTVQFNQFDCGGIALSTCVSHKIGDMCTISKFLQDWATIARDPNLKLCPQFIGSSIFPPTNEPVNEPPI

QKCVTRRLVFSNHTLKSLLSEPSQVKNPTRVELLTALLYKCGMKANSSSLKPSILFQTVNLRSFIPLPDN

TAGNFSSSLFVPTYNEEEMMLSRLVSQLRKEKEQLVANYKNCKGGQDLVSTTMRPFQEIRKLFKDMDFDM

YRCSSLANYPLYDVDFGWGKPNKISIAEGVFRNVFLLYDNKTGDEVEASVCLDEESTMSAFLREMEQFLQ

FEISSEEIKMEARCLLLKMELSRSHPTAKFNLQAPALSSSFFHLCIFVLVACFTYPIVCSSNSERYQAAL

FVFSDSVFDPGNNNYSYHYYSVQRKDSHIYMQLEKDL

>NtabASAT3_like_XP_016486234.1|NtomASAT3_like_xp_009630197.1

MAVLSTIISKKIVKPLSPTPSTQRWHKLSLLDQGIGNFYMGLVLFYPKHQVDIFQNGPKQLCILLENSLS

KAITYYYPWAGSLRDNATIACDDTGAEFLEVQVNCPMDKVLYHQDSSIKNITFPQGVPLRNAADGACLIV

SQLSHFECGGIAISVCLSHKVGDARSALFFVRDWAALTREPNAELVCPPYFVQDSLIPSPSDGPLTFPII

VSKPEERIELEKRFYFSASKIRALKDLVSAESGVENPTTTEVVSALVYKSAAIANLGSSHEQSRLILLSD

IRKEISHSAPPTSIGNILTIFSTPIYNNKGDLRVSKLVADIRKSKHELSNRDNFKENEWASAMLHAYKTG

DGTYRQSNCDVYRCSSASNIPFQDLDFGWGKPIRATMPSSPFGKMFYMMSTHDRGIEVIINLNEQQMSAF

ENDKDLLEFATPAHVED

>NtabASAT3_like_XP_016486892.1

MRDVLCTELPFPFSNSNADVYNSISICISHKIADGYSLSKFLNDWAATNRELDFEPSTQFDAASFFPPMD

DPPVIPDVVREQCVSRMFNISSYSLGKLKDIVSTNSGIQLALKLLQHLFINVDSTTTTEDEIELAHVVAQ

LRKAKQHQRDKLKDMSPDKIALHALESINVGVNIILERNYDPYMYSSLCNKGLYKTDFGWGKPISVTLAR

SPMKNNIVFLDDPSGEGINALITLTEADMLIFQSNKELLEFASPVAYSPERGPSSNPT

>NtabASAT3_like_XP_016489441.1

EFLVVSALLSSPSLVSICDKSFIKPSTLTPSTLRFYKLSFVDQSFSNMYIPVAFFYPKPQREESNNSQLS

HIADLLQTSLSQTLVSYYPYAGILRDNATVECNDMGAEFLSVKINCPMSEILNHPHATDAESIVFPKDLP

WKNNYEGGNLLVVQLSKFDCGGIAISACLSRDGCSVINFLNHWANVTRDRRFVVPSPRFVGECVRLIFPT

VKLDALQAKIAVESGVENPTRAEVVSALLYKSATKAAASSSANIQPSKLVHYLNVRTMIKPRLPRSAIGN

LLSMFSTAATQDIELPRLVHNLRKEVEVAYKKDQVDENELVLEIVDSMKKGKLPFEEKDENCTTTTMYFC

SNLCKFPFYSVDFGWGKPERVYLGTGPFKNFFFLKDYQTGRGVEARVMLQKQHMSAFECDEELLEFASSS

VPSL

>NtabASAT3_like_XP_016489747.1

MVSVSKMLSTISKKIIKPSSPTPSTQRCHKLSLLDQRPYNYMPLVFFYPKHQVATIPNGPKQLPNLLANS

VSKTLTCYYPWAGSLKDNATIDCDDSGVEFFEVQISSQMDKVIHHPNSNIKELTFRQVGDACSAYYFLRD

WAALTRNPNAKISPYFAEDSLFPSPFDGPLVASVIESKNKDCIQKRFVFSDSKLSALKAYIADESGLQNP

TRSEVVNALLYKCAAFAVSGSSGSFQRSQLIQYSNLRERISPPLPPSTIGNILTVFSTPIYNNERDLRLP

KLVRDIRKYKNNLSSKNNLEENEWVVEMLDAYRTGKELFNQRNCDVYLCSSTLMYEYEKLDFGWGRPAKG

SLGNVGQFSKNFILMNTPDRGVEAFVNLNEQEMCLFDKDKDLLEFPTPIDRDNLN

>NtabASAT3_like_XP_016490033.1|NtomASAT3_like_XP_009616124.1

MKMPSSTILSEKLIKPSSPTPSTQRWHKLSLLDQAFSNIYIPFAFFYTKKQLDSVSINPTQISHLLENSL

SKLLVSYYPYAGRLKDNTIVDCNDVGADFLRVEINIPISQALQKHNNVIEEMVFPRDLPWSNCTNRGLVV

AQLSYFNCGGIAISLCISHKVGDGHSSYNLFHDWAKITREPNGPKPVLQYVQESIFPPPITGTPFLSPVC

TSNKEDCIQRKFIFPIQKLNKLKALIAATSNVQNPTRNEVVSALIFKCIVAAVKVVNSGKSFQPWIMSQS

FDLRHQIDLPPNSIGNILTSTYISIAAEKYMTLANIVTEMRKEKQRVCDRDNVEDNLFLKKFLELATEEK

NFLKVQNNMCNFSSLVKFPVHEIDFGWGKPAKVNIANGLHNKVVFLVGNLSGVDAFVSLSAKVMSVFEKE

KELLEFAAAVPTF

>NtabASAT3_like_XP_016494255.1

MASIAQSPIGTTPENVNKSSATTISLPLVSIISEKLIKPSSPTPHTKRWHKLSLIDQAFSNSYIPFSLFY

TKQQLDAISKNNNPTQISHLLEESLSKILSTYYPYAGRLKDNTLVDCNDTGAEFIEAQISCPISQTLNWH

NSAVENLLFPQGLPWSNSADRGLVVVQLSYFGCGGIAISMCISHKIGDGCSGYNLFRDWSEITRDPNFSK

PSLHYVEQSIFPPPSSGPFLSPLFMSNTHDCVQRRYIFSNEKLLNLKNKVAAESEVQNPTRTEVVSALIF

KRAVAAANANSGFFQPSSMVQAVDLRTQIGLPPNAIGNLLTICPTSITNEQSMTISKLVSEMRKSKELAY

NRDNINDNIFVALLLELANSKQEYHDNGPNAYQITSLVKFALHEIDFGWGKPTKVNIANGLNNKLAILMG

NQSGGLDAFVTLNERDMFVFQRDPELLEFASLVPSC

>NtabASAT3_like_XP_016495009.1

MVALSVVSLISKKIIKPSIPTPLSLKSHKLSFVDQVLHISIPIAFFYPKVGNYAAYKSYEPTLVSQLLEK

SLSKTLSFYYPYAGRLKNDSSVDCNDTGVELSNVRVHCSMSEIFKHPYTEAEHVVFPQAKPHDHSKGNLA

IAQVSHFDCGGLAIGACLSHKIGDGYTASNFLYNWGVLSRDISATPSPRFVGESIYPQCNDPSDVSSIKY

EEPTGYLGKRLIFPATKVNALKANISLESGVQNPTRTEVVSALLYKCAVAARRANNVGSFKPSTLFQVAN

MRQRLNPPLSYDACGNVISGFLVKTTNERELNCPRLIREMRKEKEKLHDQQKNATKNTIISEVVDSLEKG

RTPFHSSEVDYYVCSSVLVFPFYNMDFGWGRPARVSMATGPFNKYFFLMDNQSGGGIEAIVMLGEQDMAI

FERDPELLEFAIPIPNH

>NtabASAT3_like_XP_016495885.1

IASSAYSPIVAFYSKPTNNTSQILENSLSKVLSSYYPFAGRIKDNYKYVDCNDTAAEFLNVRISCPMSEV

LNHAYNDAINVVFPPDLPWTSSFGRSPLVIQLSHFDCGGIAVSDWAATARHELDFEPSTQFDAHSFFPQI

DDLPVVPDIMREPQRCVSRIYHFSSSSLDRLKDIVAMNSQVQNPTRVEVATALLHKCGATVSMANCGMFQ

PSVLFHIMNFRPPLPLNTIGNVLCFFSSVAVKEDEIQLPQFVAQLRKAKKTSSRKVEQSKSASITCS

>NtabASAT3_like_XP_016500728.1

MVESTIVSRKMIKPSSPTPSSLSRHNLCFLDHIATSAYAPIAVFYPKPTNNISQILENSLSKVLSSYYPF

AGIITDNKYVDCNDTGAEFLNVRISCSMSEILDHAYNDAIDVVFPPDLPWTSSFGRSPLVVQLSHFDCGG

IAISICLSHKIVDGYSLFKFLKDWAATSQHLDFKPSTQFDANSFFPLMDDPPVLLRDIVREPQRCVSRIY

HFSSSSLDRLKEIVAMNSQVQNPTRVEVATALLHKCGASVSMTINSGVFQPSLLFHVMNFRPPLPLNTIG

NACSAFASAAILENEIQVPNFVAQLREAKKHLQDKLKDPEQLTSYVLETIKGTTNKTEKNIVFDFYMCTS

LCNFGLHNIDFGWGNPIRVTVATNSMKNHFIFMDSPSGDGIDVLITLTEADMLIFHNNKELLEFASPVVL

PQE

>NtabASAT3_like_XP_016504229.1

MSNVSECVQKRLIFPATKLDALRAKIAVESGVENPTRAEVVSALLYKSATKAAASSSANIQPSKLVHYLN

VRTMIKPRLRRSAIGNLVSTFSTTVGTEAFRCNRTCGQVVASASRCALKRSAQVRKMLRRCGIAPAREGA

QMRAEGCEALVA

>NtabASAT3_like_XP_016504553.1

MIFQLYPILFVNLSDICLSRIYHFSSSSLDRRKDIVAMNSQVPNPTRVEVAAALLYRCGAAVSMANFGVF

QPSILFHVMNCRPPLPQNTIGNALCFFISLAVTEDEIQVPQFVAQLRKAKKYLQEKLNDPNQLAPHVLEK

IKETTNKSEKNM

>NtabASAT3_like_XP_016505083.1

MDIDDSLAAAHINIDLLYIKFISSFRKHTAIFFFCLETQYSNHFYLKMCASNLVSSVCKKIIKPYFPTPT

SLRCHKFSYLDQMVGGIYVPFAFFYPKLSNTWSDKPSNVISEHLENSLSKVLSHYYPFAGKLNENISVDC

NDRGVEFFTTKIDCPMAKILKDPYSDKEDLVYPKGIPCTYSYEGSLAVVQLSYFNCGGIAVSACLTHKVG

DGYTIANFIRDWATIARNPSSKIQSPQFNAASIYPPTKDMVNLPEIVPKGEECSSKSFAFSSSKLAALKS

MVINNSVVQNPTATEIVSAFIYQRAMATKKKTSGSICPSVLVQAMNLRPPLPKTLMGNIGSFFSVITTEE

KEMDLPRVVGKLREAKQEVRKKYKHDKTDEFLPMTIELFREANDSFFNNSCYDIYRFSSINKFPFYDIDF

GWGKPEKVSLASNGPVKNMFVLIDNKSRDGMEVVSCMKEQDTSALERDEELLKFTSPSN

>NtabASAT3_like_XP_016506125.1

MAASRILSVSQKIIKPSFPTPLLRRHNLSFLDQISLSIYAPTAAFYSKPANNNISQILENSLSKVLSSYY

PFAGRIKAISICISHKIADGYNLSKFLNDWAATTRELDFEPSTQFDAASFFPPMDDPPVIPNVVREQCVS

RMFNISSYSLGKLKDIVSTNSEVQNLTRIEVATALVHKCGMSVSMANSGLFRPSLVTHLMNLHPHFPLNT

IGNAYCFLNSATTTEDEIQLPHIVAQLRKAKQHQRNMLKDMSSDKIALHALESINAGVNIIIEKKYDPYM

YSSLCNLGLYKIDFGWGKPINVTLARSPMKNNIVFLDNPSGEGINTLITLTEADMLIFQSNKELLEFAYP

VVYSPE

>NtabASAT3_like_XP_016512747.1

MSEIKRLIKNKSQPDRMATVQDLSIESNHLKKEVSELKASNVALDERIFKLENSDTTIDKELGTNNTETN

IGTETSKGKTAGKLLAANDYAEDEFLQKLQSKVTTRRPIKWDEVVAPQNTGNTEVFHFSIPMSFFYPKNR

NDKSYEPTHVSQLVEKSLSKALSFYYPYAGGLKDSSFVDCNDVGVELSHVRVHCPMSEIFNHPYTDAEHV

VFPVVQPYGHNEGNLAITQVSHFDCGGTAVGGCLSHKIGDGCTASSFLYNWGVLSRDISANLSRPHFVED

SIYPQSSDPSILSKSIKSEEPGYLGKRFIFPAAKVNALKAIFSMDSGVCNPTRAEVVTALLYKCAVAARR

ANFGSFRPSSLFQVADMRQRLNPSLSQDASGNIVSGFLVPITNEREMYCPRLVSEMRKEKEKLVDKDNIN

KNAFISKVIASLEKGKNNESDDYYCSSLNAFPHYETDFG

>NtabASAT3_like_XP_016514860.1

MVESMILSRKMIKPSSPTDISRHNLSFIDHIASSAYVPVAAFYSKPENNISQILENSLSRVLSSYYPFAG

IIMDNKYVDCNDTGAEFLNVRISCPMSEILDHAYNDVTDVVFPQYLPWSSYSNGSLLVVQLSHFDCGGIA

VSVCLSHKIADGYSLCKFLNDWAATARHESDFKPSTQFDAYSFFPLMDDPPVLRDIVREPQRCVSRMYHF

SSSSLGRLKDIVSTNSQVQNPTRVEVATALLHKFGAAVSMANFGMFQPSILSHLMSFRPPLSRNTIGNAC

YFFGSIASTEDEIELPHFVAQLRNAKQYLQDKSKDPNTLASHVLENVKERANKSEKSTKFDFYMCSSLCN

LGVYKIDFGWGEPIRVTLARNPMKNNFIFLDSPNGDGINVLITLREADMLIFESNKELLEFASPLV

>NtomASAT3_like_XP_009588593.1

MAISRLVLLSQKIIKPSSPTPFSHRIHKLSLMDQMGTRTYMPISFFYPKQDTAISLEPTKVSQILEKSLS

KVLTAYYPFAGRVRDNSFIECNDMGINLSQVRIDCPMSSIFNQPRTDIEKLIFPKDPWSTSTDSLIVAQL

NHFECGGIVLSACISHNVADAYSMANFVRDWALVALDSEAKPSPLFNGASIFQPANYSAPQVADPSRKQN

ASKRYHFSASKLKALKARSQIPPTTVEAVTAFLCKCANTPTFKPSLLIQAVNLRGTSNDALVPAGLVGNA

VLPYVVSAANEEDLNLQRLIGELREGKEKVHNMLKYIKSEELLCSRVSELATQINEQTSSNAFSIYRFSS

LRKLPFRDINFGWGRPTRVDIATFPVNMFVLLDNQNGDGVEVLVNLDEGEMSGFESNEELLQFASPSSGL

>NtomASAT3_like_XP_009590647.1

MTSSRLISVSEKIIKPFSATPSPLRHYKLSLIDQLMSTMYIPIAFFYRRPLEHKIINKSSKLEVSQTLEK

SLSKTLSSYYPFAGKLKDNLSIECNDMGAKFLNVELNCSMSEVVNLPDTGPEYLAFPKNLPWNSTSYYEG

TNYFVVAQLSHFRCGGIAVSACLSHKIGDGCTEINFMNDWATIARSATSSNYAIIRLSTPQWIGASIFPP

TYDDHSSAISRITPNTLPCITKRYVFSSSKLSALKKFMASDSGGVQNPTRNEVVTALLYKCANMATSRSS

SGSGLFKPSALIQVVNLRPRLNPPLPNNSAGNLISSILIKSTDEEQMKVARIVQDLRKEKEQLNMKHDLV

NQNGVVLSALEYLNDNMDVYCCSSLCNYQLYNVDFGWGKPERVVVTNGTINFFLLSDDKNGDGIEVLVSL

GKEVMSEFNNNKELLEFASPVSN

>NtomASAT3_like_XP_009596353.1

MAVSRIISVSEKIIKPSSATPSPLRNYKLSLLDQVMSTMEYCYNNLDKIDQFNDIEQMIRARVIQELRKG

KEQLKKKHHVKQNGLLSALQYFDEGIRPDNTNDNIVDVCWFTSLCNYQLYNVDFRWGRPERAVVASGGLN

SIFLSDDREGDGVEALVSLDKAVMSEFNTNKELLEFASVLLN

>NtomASAT3_like_XP_009603396.1|NtomASAT3_like_XP_033512582.1

MAASGLLSSPSLVSICDKSFIKPSTLTPSTLRLHKLSFVDQSVSNMYIPVAFFYPKPQREESNNSHLSHI

ADLLQTSLSQTLVSYYPYAGILRDNATVECNDMGAEFLSVKINCPMSEILNHPHATDAESIVFPKDLPWK

NNYEGDNLLVVQLSKFDCGGIAISACLSHKIGDGCSVINFLNHWANVTRDRRFVVPSPRFVGDSILSSLN

GPLVAPQIMSNVGECVQKRLIFPAAKLDALRAKIAVESGVENPTRAEVVSALLYKSATKAAAASSANIQP

SKLVHYLNVRTMIKPRLRRSAIGNLVSTFSTAAMQDIELPRLIHNLRKEVEVAYKKDQVDENELVLEIID

SMKKGKLPFEEKDENSTTTTMYFCSNVCKFPLYSVDFGLGKPERVCLANGPSKNYFFLKDYKTGRGVEAR

VMLQKQHMSAFECDEELLQFASSSLPSL

>NtomASAT3_like_XP_009604796.1

MAASALLSSLVSICDKSFVKPSCLTPPMVRYHKLSFVDQSLSNMYIPFAFFYPKQQKEESNNSQLSHLAD

LLQRSLSQTLVSYYPYAGILRDNATVECNDMGAEFLSVKINCPMSEILNHPHATDAESIVFPKDLPWKNN

YEGDNLLVVQLSKFDCGGIAISACLSHKIGDGCSVINFLNHWANVTRDRRFVVPSPRFVGDSIFPSQNGP

LIAPQTMSNVSECVQKRLIFPTAKLDALRAKIAIESGAENPTRAEVVSALLYKSATKAAASSSANIQPSK

LVHHLNVRTMIKPRLPRSAIGNLLSMFSTAATQDIELPRLVHNLRKEVEVAYKKDQVDENELVLEIVDSM

KKGKLPFEEKDENCTTTTMYFCSNLCKFPFYSVDFGWGKPERVCLGTGPFKNFFFLKDYQTGRGVEARVM

LQKQHMSAFECDEELLQFASSSVPSL

>NtomASAT3_like_XP_009604797.1

MAASALLSSLVSICDKSFVKPSCLTPPMVRYHKLSFVDQSLSNMYIPFAFFYPKQQKEESNNSQLSHLAD

LLQRSLSQTLVSYYPYAGILRDNATVECNDMGAEFLSVKINCPMSEILNHPHATDAESIVFPKDLPWKNN

YEGDNLLVVQLSKFDCGGIAISACLSHKIGDGCSVINFLNHWANVTRDRRFVVPSPRFVGDSIFSSLNGP

LIAPQIMSNVSECVQKRLI

>NtomASAT3_like_XP_009606246.1

MAAVPSQRISIPEKTIVKPSSPSPSSLNPILKNVRAHRKFLFSGSKLGALRAIVAVESGVKNPSRSEVVS

AIMFKFVTKAASRINNNSGTFRPSMMLNDVDIRPLVVPPLPQNSIGNLLSVFLLVATKEDEMKIATLVYN

LRKELEAVYKKDPVKQNALILDGENQEEYHHHLVRGRIACLRGVEALVTLEEKQMLALENDEEFLEFATP

VTSY

>NtomASAT3_like_XP_009607076.1

MGVLSRLHKLCLPGQAFTPSPIPLVFLYSKQQVDTIFPNGPKQQLSKFLEISLSKTLTCYYPWAGRLKDN

ATVECNDVGAEYFEVQINSPMDEVVHHSDSRIKDMVTFPQGATWGNCTDRALAIAQLIHFECGEIAISVC

LSHKVGDGCSCYWFLRDGASLTRGPNAKIASTPYFVEDSVILSPLPDGPLVSPSGMSTLIYNSSREGDFN

LSKVVADIRNSKQKLLSRDN

>NtomASAT3_like_XP_009629726.1

MIKPRLRRSAIGNLVSPFSTAATQDIKLPRLIHNLRKEVEVAYKKVQVDENVLVLEIVDSMKKGKLPFEE

KDENSTTTTYFCSNVCKFPLYSVDFGLGKPERVCLANDPSKNYFFLKDYKTGRGVEARVKDYKLLQFASS

SVPSF

>NtomASAT3_like_XP_009631587.1

MASIDQSPIGTTPENINKPFSATTISLPLVSIISEKLIKPSSPTPHTKRWHRLSLIDQAFSNSYIPFSLF

YTKQQLDAISKNNNPTQISHLLEESLSKILSTYYPYAGRLKDNNVVDCNDTGAEFVEAQISCPISQTLNW

HNSATEDLLFPQGLPWSNSADRGLVAVQLSYFNCGGIAISMCISHKIGDGCSGYNLFRDWSEITRDPNFS

KPSLHYVEQSIFPPPSSGPFLSPLFMSNKHDCVQRRYIFSNEKLLHLKNRVAAESEVQNPTRTEVVSALI

FKRAVAAAKANSGFFQPSSMVQAVDLRAQIGLPPNAIGNLLTICPTSITNEQSMTISKLVSEMRKSKELA

YNRDNIFVALLLELANSKQEYHDNGPNAYQITSLVKFALHEIDFGWGKPTKVSIANGLNNKLAILMGNQS

GGLDAFVTLSEQDMSVFERDPELLEFASLVPSC

>NtomASAT3_like_XP_018631230.1

MCASNLVSSVCKKIIKPYFPTPTSLRCHKFSYLDQMVGGIYVPFAFFYPKLSNTWSDKPSNVISEHLENS

LSKVLSHYYPFAGKLNENISVDCNDRGVEFFTTKIDCPMAKILKDPYSDKEDLVYPKGIPCTYSYEGSLA

VVQLSYFNCGGIAVSACLTHKVGDGYTIANFIRDWATIARNPSSKIQSPQFNAASIYPPTKDMVNLPEIV

PKGEECSSKSFAFSLSKLAALKSMVINNSVVQNPTATEIVSAFIYQRAMATKKKTSGSICPSVLVQAMNL

RPPLPKTLMGNIGSFFSVITTEEKEMDLPRVVGKLREAKQEVRKKYKHDKTDEFLPMTIELFREANDSFF

NNSCYDIYRFSSINKFPFYDIDFGWGKPEKVSLASNGPVKNMFVLIDNKSRDGMEVVSCMKEQDTSALER

DEELLKFTSPSN

>NtomASAT3_like_XP_033507977.1

KATIGEYAELAIDNLSFIDQIASSAYSPIVAFYSKPTNNTSQILENSLSKVLSSYYPFAGRIKDNYKYVD

CNDTAAEFLNVRISCPMSEVLNHAYNDAINVVFPPDLPWTSSFGRSPLVIQLSHFDCGGIAVSVCLSHKI

ADGQSLXKFLKDWAATARHELDFEPSTQFDAHSFFPQIDDLPVVPDIMREPQRCVSRIYHFSSSSLDRLK

DIVAMNSQVQNPTRVEVATALLHKCGATVSMANCGMFQPSVLFHIMNFRPPLPLNTIGNVLCFFSSVAVK

EDEIQLPQFVAQLRKAKKHLQEKLNNPNQLASHVLEKIKATTNKPEKNKVDFYMCSSLCNFELYNIDFGW

GKPIRVTLATNSMKNHFIFLDSPNGNGIDALITLKEADMLIFQSNKELLQFASPFVLPQE

>NtomASAT3_like_XP_033512495.1

MDDPPVIPNVVREQCVSRMFNISSYSLGKLKDIVSTNSEVQNLTRIEVATALVHKCGMSVSMANSGLFRP

SLVTHLMNLHPHFPLNTIGNAYCFLNSATTTEDEIQLPHIVAQLRKAKQHQRNMLKDMSSDKIALHALES

INAGVNIIIEKKYDPYMYSSLCNLGLYKIDFGWGKPINVTLARSPMKNNIVFLDNPSGEGINTLITLREA

DMLIFQSNKELLEFAYPVVYSPE

>NtomASAT3_like_XP_033514428.1

MAASALVSLSKKIIKPFSPTPFSERIYKLSFIDQFNSTQYCPLVFFYPKNKGNVVTPSIEPSDMCKVIEN

SLSKTLAAYYPFAGTLRDNVHVECNDIGADFFKARFDCPMSEIVKSPDRNVKEMVYPKGIPWNIVTSNRK

LVTVQFNQFDCGGIALSTCVSHKIGDMCTMSKFLQDWAAIARDPNLKLCPQFIGSSIFPPTNEPVNEPPI

QKCVTRRFVFSNHTLKSLLSEPSQVKNPTRVELLTALLYKCGMKANSSSLKPSILFQTVNLRSFIPLPDN

TAGNFSSSFFVPTYSEEEMKLSRLVSQLRKEKEQLVGNYKNCKGGEDLISTTMRPFQEIRKLFKDMDFDM

YRCSSLANYPLYDVDFGWGKPNKISIAEGVFRNVFLLYDNKTGDEVEASVCLDEESTMSAFLREMEQFLQ

FEISSEEIKMEARC

>NtomASAT3_like_XP_033515849.1

MAASALLSSLVSICDKSFVKPSCLTPPMVRYHKLSFVDQSLSNMYIPFAFFYPKQQKEESNNSQLSHLAD

LLQRSLSQTLVSYYPYAGILRDNATVECNDMGAEFLSVKINCPMSEILNHPHATDAESIVFPKDLPWKNN

YEGDNLLVVQLSKFDCGGIAISACLSHKIGDGCSVINFLNHWVNVTRDRRFIVPSPRFVGDSIFSSLNGP

LIAPQIMSNVSECAQKRLVFPTAKLDALRAKIAVESGVENPTRAEVVSALLYKSATKAAASCSANIQPSK

LVHYLNVRTMIKPRLPRSAIGNLLSMFSTAATQDIELPRLIHNLRKEVEVAYKKDQVEQNELVLEIVESI

QKGKLPFEENDENCTTPTTMYFCSNLCKLPFYSVDFGWGKPERVCLGTGPFKNFFFLKDY

>NtomASAT3_like_XP_033516706.1

MSPKDETEQEYMSRVPYANVVVFPQDLPWSSYSNGSLLVIQLSHFDCGGIAVSVCLSHKIADGYSLLKFL

KDWAATTRHELDFKPSTQFDAHSFFPPMAMNDLPAVMPDIVRETQRRVSRMYHFSSSSLGRLKEIVAMNS

QVQNPTRVEVATALLHKCEAAVSMANLGVFQPSLLCHLMNFRPPLPQNTIGNACYFFGSIAATEDEIQLP

HFVAQLRKAKQYLQDKSKDPHQLASHLLERVKERANKSEKNKFDFYMCSSLCNLGVYKIDFGWGEPIRVT

LARNPMKNNFIFLDSPNGNGINVLITLTEADMLIFESNKELLEFTSPVV

>NtabASAT3_like_XP_016453912.1

MAALPLLSSDSLVSIRDKSFIKPSTLTPSTLRFHKLSFVDQAVSNMYIPVAFFYPKPQRE

ESNNSQLSHIADLLQRSLSQTLVSYYPYAGILRDNATVECNDMGAEFLSVKINCPMSEIL

NHPHATDAESIVFPKDLPWKNNYEGGNLLVAQLSKFDCGGIAISACLSHKIGDGSSVNNF

LNDWANVTRDRRLVPSPRFVGDSIFSSLNGPLIAPQIMSNVSECVQKRLIFPAAKLDALR

SKIAVESGVENPTRAEVVSALLYKCATKAAAASSSANIQPSKLVHYLNVRTMIKPRLRRS

AIGNLVSTFSTAAMQDIELPRLVHNLRKEVEVAYKKDQVDENELVLEIVDSMKKGKLPFE

EEDENSTATMYFCSNVCQFPLYSVDFGWGKPERVCLANGPCKNYFFLKDYKTGRGVEARV

MLQKQHMSAFECDEELLQFTSS

>HnigASAT5_c40105_g1_i1

MEIISKEFIKPISPTPTHLKSYQYSILDQFAPSNYIPILIYYPPPSKNQEPNNATAINTSVSKQLKKSLSQILAHYYFLAGRLNIANNCVDCNDEGVPFLEAFVRNKNLEDILNNLHDENKLKESLLPLTIESFTSTLLLSIQVSFFECGGMILGISATHKVLDTSSLRTFLTDWTSLARNDDPNFVISPPVPAIAKLLPPINELPPSPNTEAAISKEACVTNVYVFSPPSILDLKKKIVSEDVLKPTRVEVVTSVLWKFLTANSKRPSLLCHLVNMRNRVDPPMHAHCIGNFVGFMVASKEESEKDVANMVVALRKGLSEFDDQYAKRLKGDEAIGTTLELLKYSLTLLSKKNEVDLYHFSSWCGYSFYDVDFGWGKPIMFSIIEANIKNMVFLMDTKDGGVEAWVTLTEENMKILEKNSTELLTYATLKASFLKKE

>HnigASAT4_c56915_g1_i1

MAELEIQIHLRKMVKPSTPTPNHLTTHKLSLFDQVAPHIFVPILFHYLPNNECNMEEITERCDKLQKSLAESLTRFYPLAGRLSKDNLSIHYNDEGVEYVESKVNTNLAEFLHERLKIELLNDFLPTDLPSSFLLRVQVNVFNCGGIVIGINISHIVADGFTLGSFVKEWAHLSQTGTSKGCLTSFDHLPVVFPPRALSGPQVSPPSNIGPKIVTKRFVFDALAISKLKDRINSSSTFTKPTRLVVVLSFIWKVLVGNSSAKHGHSRDSSLLFPINLRGKSNLPSIEHALGNFYVTEVATLEANQSRKELTDFVNVVGSKTRDTCVSIAKASADDIASMFVNCGMEVIKKLGQGDKIDIYPSTSWCRFPWYEADFGWGKPFWVSSVSKPFEVIILNDTKDSDGIEAWISLKENEMIEFERDPEILSSTSKVAFDSSMWIK

>HnigASAT1_c58659_g1_i1

MAASALLSLSKKIIKPFSPTPFSERIYKLSFIDQFNTTQYNPLAFFYPKNKGVPSIDPNDMCKVIENSLSKALAAYYPFAGTLRDNIYIECNDIGADFYKARFDCPMSEILKSQDRNVKEIVYPKDVPWNIVTPSRKLVVVQFNQFDCGGIALSACVSHKIGDMHTFYKFMHDWAAISLCSNVNICPQFIGSSVFPPTNEAVNEPPREQCVTKRLLFSNHALKSLLPGSSEVKNPTRVEILTALLYKCSMRANSSGLFKPSMLFQTINLRPIIPLPENTPGNFSSSLFVPTYTEEEMKLSRLVSELRKGKEKYFDDYRKCKEGHQDMVSTTTRPYQEIRALFKDNDFDLYRCSSLVNYGWHGLDFGWGMPNRVSMADVKLRNIFMLFDNNTEDHVEAQVSFDKESKMSAFLREIEQVLEFKLHA

>HnigASAT3_c61998_g1_i2

MSLSSIVSSACKIIKPYSPTPTSLRCYKLSYLDQMLGGIYVSFALFYPKLSYTCLNTPSNLSEHIEKSLSKVLTHYYPFAGKMNGNVSVDCNDSGVEFFTTKIDCPMSKIQDDPYADKEDLVYPKGIPMTYSYDGSLAVFQLSYFNCGGIAISACLTHKLGDAYTMTNLLSDWAAISRNPSSKQPSRQFNGASFFPPTKEDLSKESNSIVPEREECSSTSFAFSSSKLAALKARFINQPKVYNPTDTEVVSAFIYQRAMATKKVTSGSIRPSVLHQAANLRPPCPKNTMGNIGSLFAILTKEEKEMDVARVVGQLQKAKEELRQKYKHANIEELLPITIGQFKDANDLCVNECYDLYRFSSVTKFPIYDIDFGWGKPEKISLSSNGPIKKLFSLDG
